# 2.6-Å resolution cryo-EM structure of a class Ia ribonucleotide reductase trapped with mechanism-based inhibitor N3CDP

**DOI:** 10.1101/2024.10.09.617422

**Authors:** Dana E. Westmoreland, Patricia R. Feliciano, Gyunghoon Kang, Chang Cui, Albert Kim, JoAnne Stubbe, Daniel G. Nocera, Catherine L. Drennan

**Affiliations:** Howard Hughes Medical Institute, Massachusetts Institute of Technology, Cambridge, MA 02139; Department of Biology, Massachusetts Institute of Technology, Cambridge, MA 02139; Department of Chemistry, Massachusetts Institute of Technology, Cambridge, MA 02139; Department of Chemistry and Chemical Biology, Harvard University, Cambridge, MA 02138

**Keywords:** Radical chemistry, proton-coupled electron transfer, enzyme inhibition, cryogenic electron microscopy

## Abstract

Ribonucleotide reductases (RNRs) reduce ribonucleotides to deoxyribonucleotides using radical-based chemistry. For class Ia RNRs, the radical species is stored in a separate subunit (β2) from the subunit housing the active site (α2), requiring the formation of a short-lived α2β2 complex and long-range radical transfer (RT). RT occurs via proton-coupled electron transfer (PCET) over a long distance (~32-Å) and involves the formation and decay of multiple amino acid radical species. Here, we use cryogenic-electron microscopy and a mechanism-based inhibitor 2′-azido-2′-deoxycytidine-5′-diphosphate (N_3_CDP) to trap a wild-type α2β2 complex of *E. coli* class Ia RNR. We find that one α subunit has turned over and that the other is trapped, bound to β in a mid-turnover state. Instead of N_3_CDP in the active site, forward RT has resulted in N_2_ loss, migration of the third nitrogen from the ribose C2′ to C3′ positions, and attachment of this nitrogen to the sulfur of cysteine-225. To the best of our knowledge, this is the first time an inhibitor has been visualized as an adduct to an RNR. Additionally, this structure reveals the positions of PCET residues following forward RT, complementing the previous structure that depicted a pre-turnover PCET pathway and suggesting how PCET is gated at the α–β interface. This N_3_CDP-trapped structure is also of sufficient resolution (2.6 Å) to visualize water molecules, allowing us to evaluate the proposal that water molecules are proton acceptors and donors as part of the PCET process.

**Significance Statement:** Several FDA-approved cancer drugs target human ribonucleotide reductase (RNR), a radical enzyme that produces the requisite deoxyribonucleotides for DNA biosynthesis and repair. Human RNR is a class Ia enzyme that requires radical transfer (RT) from a β2 subunit to an α2 subunit on every round of turnover. Long-range RT is both a remarkable feature and an Achilles heel, given that inhibitors can intercept the radical species. Here we present a cryogenic electron microscopy (cryo-EM) structure of the best studied class Ia RNR, the enzyme from *E. coli*, in which α2 and β2 subunits have been trapped together using a mechanism-based inhibitor. This structure provides insight into both the mechanism of RNR inhibition and the mechanism of long-range RT.

## Introduction

Ribonucleotide reductases (RNRs) are essential enzymes in DNA biosynthesis and repair. They convert all four ribonucleotides (CDP, UDP, ADP, GDP) into the corresponding deoxyribonucleotides (**Fig. 1A, B, S1**) (1, 2). The 2′ hydroxyl group is lost in the form of water with reducing equivalents coming from the oxidation of a pair of cysteine residues (**Fig. 1A, S2B**). To afford the chemically challenging removal of the hydroxyl group from a five-membered ring, radical-based chemistry is employed. An amino acid (thiyl) radical is transiently formed on the α subunit (3). This cysteine (Cys439 in *Escherichia coli*) is not near the protein surface, instead sitting in a loop in the center of a 10-stranded α/β barrel fold (4). Three different strategies have evolved for generation of this transient thiyl radical on a relatively buried cysteine. Class II RNRs use adenosylcobalamin, which is small enough to enter the barrel and generate the thiyl radical directly (5). Class III RNRs employ a post-translationally installed glycyl radical species that can generate the thiyl radical species through short-range radical transfer (~3.5 Å) (6–8). Finally, class I RNRs employ a second subunit (β) for radical storage (9), necessitating a long-range radical transfer (~35 Å) between α and β subunits for thiyl radical generation (4, 10). Whereas some microorganisms encode all three RNR classes and/or multiple RNR subclasses, humans only use a class Ia RNR. To the best of our knowledge, the long-range (~35-Å) radical transfer that is required by class I RNRs is unparalleled in biology and is a focus of this study.

**Figure 1.**
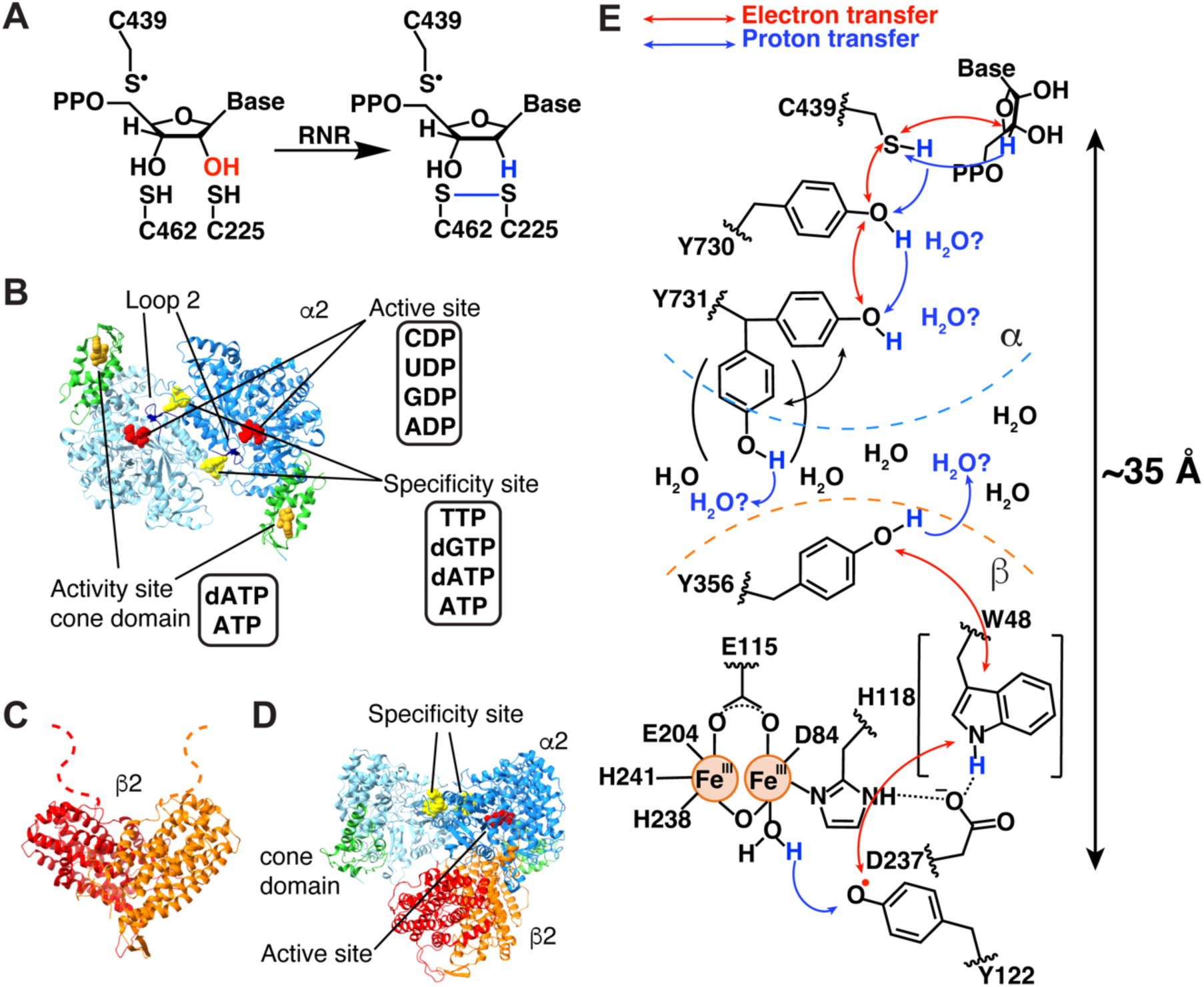
*E. coli* class Ia RNR reaction and structures. **(A)** Reaction of *E. coli* class Ia RNR. **(B)** The α2 subunit of *E. coli* class Ia RNR contains three nucleotide binding sites. The activity site (gold) in the cone domain (green) upregulates RNR activity when ATP is bound and downregulates RNR activity when dATP is bound. The specificity site (yellow) at the dimer interface dictates which nucleotide (red) is bound in the active site by modulating the hydrogen bonding interactions of loop 2 (navy blue). **(C)** The β2 subunit contains the Fe_2_O-Y122● cofactor responsible for the initiation of radical transfer. **(D)** The active state of class Ia RNR is an asymmetric α2β2 complex. **(E)** A proposed PCET pathway in *E. coli* class Ia RNR. Electron transfer involving W48 has not been experimentally demonstrated. Additionally, the role of Y731α flipping in facilitating PCET has not been determined.

This long-range radical transfer (RT) from a diiron-tyrosyl radical cofactor in β to α has been investigated extensively using the *E. coli* class Ia RNR as the model system (11). Uhlin, Eklund, and coworkers provided the first structures of the α2 and β2 subunits of *E. coli* class Ia RNR (4, 12). α2 structures revealed the locations of the active sites, of the specificity sites that allosterically regulate the preference of the enzyme for each of its four substrates, and of the activity sites that allosterically regulate overall enzyme activity (**Fig. 1B, S1**) (13, 14). The β2 structure (**Fig 1C**) revealed the proximity of the stable radical in β (Y122β•) to the diiron site and led to a symmetric docking model of α2β2 (4), which was the working model of an active RNR for almost three decades (**Fig. S1B**). Subsequent research by Stubbe, Nocera, and Bennati using unnatural amino acids and spectroscopy confirmed that radical transfer occurs over long distances and involves proton-coupled electron transfer (PCET) and the formation of multiple transient amino acid radical species (2, 11, 15–17). These RT residues include Y356 on β’s highly flexible C-terminal tail and α residues Y730 and Y731 (**Fig. 1E**) (18, 19).

The first experimental structure of the active α2β2 complex of class Ia RNR came 30 years after the docking model, obtained by Kang, Drennan, and co-workers in 2020 (**Fig. 1D**) (20). The active state was trapped by using a doubly substituted β2, E52Q/(2,3,5)-trifluorotyrosine122(F_3_Y122)-β2 and wild-type α2 and was solved by cryogenic-electron microscopy (cryo-EM) to 3.6-Å resolution. This structure enabled the first structural visualization of the residues known to participate in PCET along the radical transfer pathway (~32 Å, as measured from the structure) (20). This cryo-EM structure also demonstrated that α2β2 is asymmetric, with β2 interacting with only one of the α2 subunits, in contrast to the symmetric docking model (**Fig. 1D, S1B)** (4). Consistent with the observation that one product molecule was produced when wild-type α2 is mixed with E52Q/(2,3,5)-F_3_Y122-β2 (21), the cryo-EM structure showed one α protomer to be in a post-turnover state (α′) and the other in a pre-turnover state (α). α′ made no significant contacts with β2, had density for a Cys462-Cys225 disulfide, and had an empty active site; i.e., it was a post-turnover state. In contrast, α made substantial contacts with β2 and had reduced cysteines and substrate GDP in its active site, with allosteric effector TTP in the neighboring specificity site; i.e., it was in a pre-turnover state. Apparently, during the time course of the cryo-EM grid preparation, β2 reacted with α′, producing one dGDP, and flipped over to α, where it was trapped pre-turnover (**Fig. S2A**). The catalytically essential C-terminal tail of β, which is disordered in all crystal structures, was also disordered on the α′ side. However, on the α side, the β tail was fully ordered and wrapped in the α active site where it contributed residue Y356β to a fully ordered PCET pathway.

One surprise of this structure determination was that no polar residues were found within 3.5 Å of Y356β that could act as a proton acceptor in the oxidation of Y356β to Y356β• or as a proton donor in the subsequent reduction of Y356β• to Y356β as the electron moves along the 32-Å PCET pathway. This observation suggested that water molecules might be the direct proton acceptor/donor for Y356β (**Fig. 1E**). The 3.6-Å resolution cryo-EM structure also showed a distance of 8.3 Å across the β–α interface between Y356β and Y731α (**Fig. S2C**) (20). Although this distance is reasonable for electron transfer, it has been suggested that Y731α could flip into the interface to provide a shorter distance (22). Although the cryo-EM map for Y731α showed no indication of an alternative flipped position, the structure provided only one snapshot and that snapshot was pre-turnover and was obtained with a double-substitution of β2. It was therefore possible that β2 modification altered the conformational orientations of side chains relevant to PCET.

Here, we report the use of mechanism-based inhibitor 2′-azido-2′-deoxycytidine-5′-diphosphate (N_3_CDP) to trap a wild-type α2β2 complex of *E. coli* class Ia RNR. We obtain a 2.6-Å resolution cryo-EM structure that enables us to investigate postulated water channels connecting the PCET pathway to bulk solvent, to resolve water molecules both along the PCET pathway and at the α–β interface, and to consider proposals for how PCET across the α–β interface is gated. We additionally explore inhibition of RNR by N_3_CDP and probe the proposed asymmetric mechanism for catalysis that involves β2 swinging between α′ and α subunits.

## Results

Our 2.6-Å resolution structure of a trapped α2β2 complex was obtained using wild-type protein, N_3_CDP, dATP as the specificity effector, and ATP as the activity effector (**Fig. 2A**). We collected 6007 movies at 0.415 Å/pix super-resolution on a Titan Krios 300 kV electron microscope equipped with a Gatan GIF K2 camera and processed these movies using RELION-4.0 (23) to obtain the 2.6-Å resolution structure that we present here (**Table S1, S2, Fig. S3-S6)**. Water molecules were added by comparison to known crystal structures of the individual α2 (24–28) and β2 (27, 29, 30) subunits (**Tables S3-S6**) and by using a combination of the “Find Waters” function in Coot (31) (**Table S7**) and manual inspection (**Table S8**).

**Figure 2.**
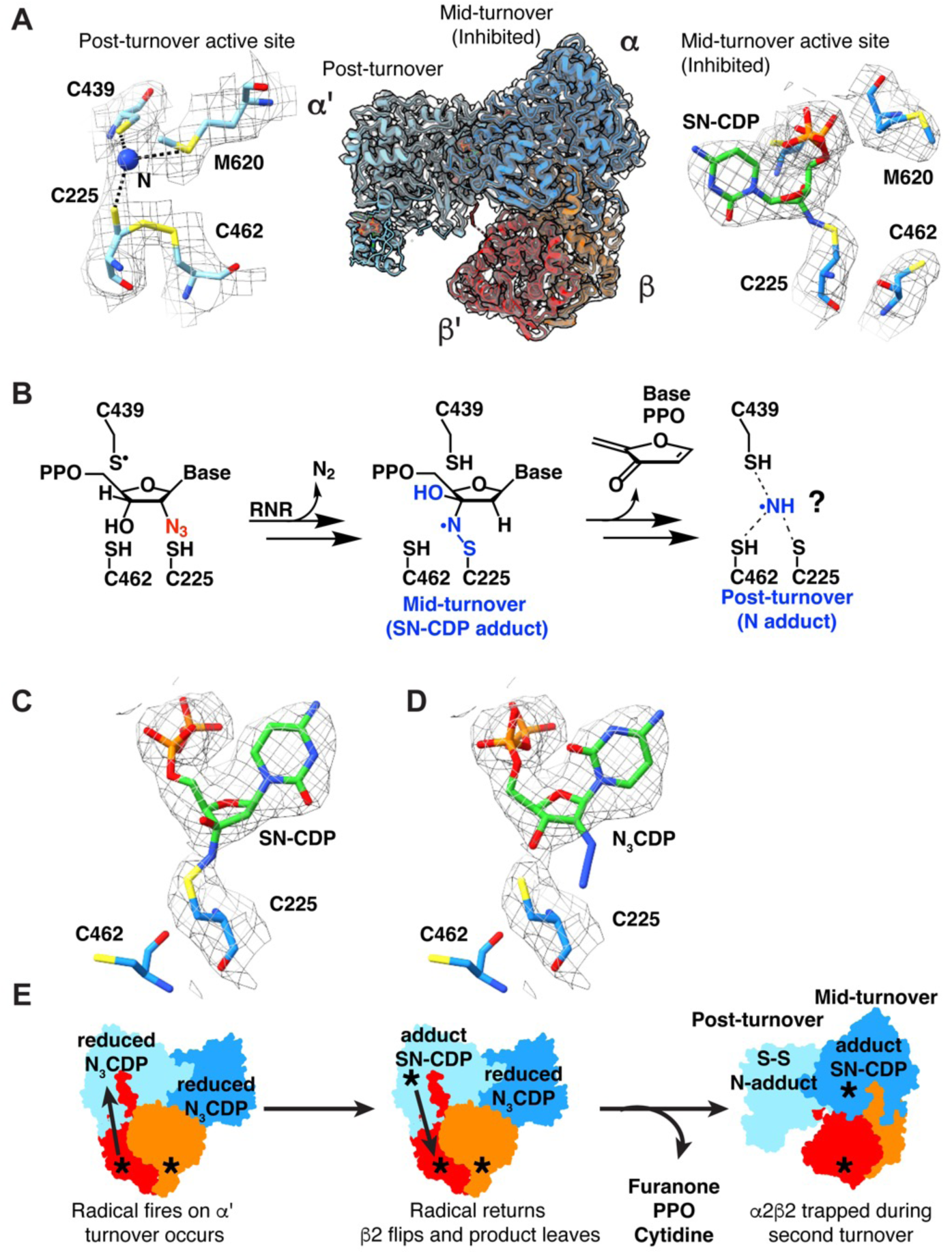
The two sides of the N_3_CDP-inhibited asymmetric complex are trapped at different stages of turnover. **(A) (Left)** 2.6-Å cryo-EM map (grey mesh; sd level 5) for α′ active site after reaction with N_3_CDP. A single nitrogen atom adduct (blue) is modeled. Dashed lines indicate distances of ≤ 3.5 Å. **(Middle)** Cryo-EM map (grey mesh; sd level 5) of active state of RNR trapped with N_3_CDP substrate. The α2β2 model is asymmetric, where α′ (light blue) and β′ (red) have undergone turnover. α (dark blue) and β (orange) have not yet released product and display a fully intact radical transfer pathway. **(Right)** Cryo-EM map (grey mesh; sd level 5) for α active site trapped with adduct from reaction with N_3_CDP. **(B)** Proposed reaction with N_3_CDP that first leads to the SN-CDP adduct, followed by release of the indicated decomposition products, potentially with a nitrogen species of some sort left behind. **(C)** Cryo-EM map (grey mesh; sd level 3) modeled with sulfur-nitrogen-CDP (SN-CDP) adduct. **(D)** Cryo-EM map (grey mesh; sd level 3) modeled with N_3_CDP. **(E)** Schematic depiction of reactions with N_3_CDP that could have led to the obtained cryo-EM structure.

### Overall structure is asymmetric with each active site displaying a different timepoint of RNR inhibition by N_3_CDP

Our overall structure is consistent with the asymmetry observed in the prior cryo-EM structure (**Fig. 1D**, **S7**) (20): β2 is primarily contacting α and not α′ (**Fig. 2A, middle, Table S9**). Also, as observed before, we find that the α active site is occupied whereas the α′ active site has no nucleotide substrate bound. These structural observations are consistent with an α′ active site that reacted completely with one N_3_CDP such that the nucleotide-based reaction products dissociated and β2 has flipped to become trapped on α, with α not having completed its turnover of N_3_CDP (**Fig. 2A**). The reaction of *E. coli* class Ia RNR with mechanism-based inhibitor N_3_XDP (X = U or C) has been previously studied (32–39) (**Fig. 2B**). Briefly, reaction with N_3_CDP requires α2 and β2, establishing N_3_CDP as a mechanism-based inhibitor (32). N_2_ is lost and an adduct with a nitrogen-centered radical species is formed (37–39). The chemical structure of the adduct was predicted from spectroscopic data to contain a covalent bond between the sulfur of Cys225 and a nitrogen atom bound to the 3′ carbon of the CDP ribose (**Fig. 2B, S8**) (39). Consistent with this prediction, we observe continuous density in the cryo-EM map between Cys225 and the 3′ position of the CDP ribose (**Fig. 2A “mid-turnover” right, 2C**). Refinement of an “SN-CDP” (sulfur-nitrogen-CDP) adduct to Cys225 (see Methods) shows that the adduct fits the density with geometric parameters that agree with the spectroscopic data (39) (**Figs. S8, S9**). In contrast, N_3_CDP is not consistent with the cryo-EM map (**Fig. 2D**). We interpret this result as an indication that β2 fired a radical into the α subunit, causing the loss of N_2_ and the formation of the SN-CDP adduct, and therefore we refer to this structure as “mid-turnover.”

As previously reported (36), the SN-CDP adduct is subject to decomposition, the mechanism of which is not fully established. However, products of the decomposition have been observed to include 2-methylene-3(2H)-furanone, inorganic diphosphate, and cytidine (**Fig. 2B**) (37, 39). Ammonia or an N atom-based adduct is the expected final product (**Fig. 2B, “post-turnover”**) (37, 39). The cryo-EM map for α′ is consistent with a post-turnover state: the density in the active site is too small for a nucleotide, furanone, diphosphate or cytidine, but large enough for an N atom or ammonia (**Fig. 2A, left**). Also indicative of a post-turnover state is partial occupancy for a disulfide between Cys225 and Cys462 (**Fig. 2A, left**). Accordingly, we have modeled a disulfide with partial occupancy between Cys225-Cys462 and have modeled the extra density between Cys225 and Cys439 as a single N atom based on the biochemistry (37, 39); our density cannot distinguish the type of atom or the number of atoms conclusively (**Fig. 2A “post-turnover” left**).

Additionally supporting a post-turnover state of α′ is the observation that loop 2, which communicates between the specificity site in α and the active site in α′, has flipped away from the active site as would be expected when product departs (**Fig. S10A**). Loop 2 of α′ no longer contacts the specificity effector dATP (**Fig. S10D**), which is bound similarly here as observed before (**Fig. S10A, B, S11**), and resembles the loop 2 conformation from an oxidized (post-turnover) *E. coli* class Ia RNR crystal structure (25) (**Fig. S10B**). In contrast, loop 2 of α beautifully mirrors the conformation of loop 2 from a previous reduced (pre-turnover) structure with CDP-dATP substrate-effector bound (27) (**Fig. S10C**, **S10E**, **S11**). Collectively, these observations indicate that α′ is post-turnover and α is mid-turnover (**Fig. 2E**). This asymmetry is consistent with our previous structure (20). However, when E52Q/(2,3,5)-F_3_Y122-modified β2 was used instead of wild-type β2, the radical was unable to fire into the second active site (21), which yielded a post-turnover structure in α′ and a pre-turnover (not mid-turnover) structure in α.

### Presence of allosteric activator ATP does not result in substantial changes to the positioning of the cone domains within the α2β2 state

ATP is proposed to act as an allosteric activator of *E. coli* class Ia RNR by displacing the allosteric inhibitor dATP, thereby destabilizing the α–β interface that is present in the inactive α4β4 state (28). ATP could, in theory, additionally increase RNR activity by stabilizing the α–β interface that is present in the active α2β2 state. Thus, we were interested to compare this α2β2 structure, determined in the presence of ATP, to the previous structure determined in ATP’s absence. As expected (28), we find two ATP molecules bound in the allosteric activity sites in the N-terminal cone domain of α (**Fig. S12**). One ATP is bound to the same site that dATP occupies in the inactivated α4β4 ring structure (site 1) and the other ATP is in a site that is specific for ribonucleotides (site 2) (**Fig. S12C-E**) (28). Despite the presence of these two ATP molecules, there are no substantial changes in the cone domain’s interactions with β compared to the α2β2 structure without ATP (**Fig. S7**). The interface between the cone domain and β is small, approximately 500 Å^2^ as determined by PDBePISA (40). These structural data are consistent with the proposal that ATP activates *E. coli* class Ia RNR by freeing β2 from the inactive α4β4 state rather than by stabilizing the active α2β2 state (28, 41).

### Water molecules at the α–β interface may play a role in communicating the presence of a bound cognate substrate-effector pair to β-tail binding residues

In an active α2β2 complex, the β-tail (residues D342 to L375) extends from β into the α active site, running along a strand of a hairpin (residues V642-V655) that is part of the α subunit (**Fig. 3A**); the β-tail makes a sharp turn as it reaches the substrate and runs back down toward the β subunit before making another turn. Following this turn, the β-tail contributes residue Y356 to the PCET pathway and extends for another 19 residues to the end of the β chain. The β-tail makes twenty-six of the direct interactions at the α–β interface whereas residues outside of the tail make eighteen (**Fig. S13, Table S10**). The β-tail also makes most of the water-bridged interactions across the interface (17 of 23 interactions) (**Fig. S13B, Table S11**). The binding pocket for the β-tail is transient. The hairpin in α that contacts the β-tail is known to re-position toward the active site upon the binding of cognate substrate-specificity effector pairs to α (**Fig. 3A**). In particular, substrate-effector binding causes the barrel that houses the active site to clamp, securing the substrate and triggering the hairpin to re-position (27). Since the hairpin doesn’t contact the substrate or effector, the molecular rationale for this movement was not clear initially. However, when the first α2β2 structure was determined, this hairpin was observed to be responsible for forming the β-tail binding pocket. Thus, we understood that the hairpin must move to create the binding site for the β-tail, allowing β to bind when a cognate substrate-effector pair is present but not otherwise, thereby providing a molecular safeguard against unwanted radical chemistry. We didn’t, however, understand the molecular basis for the hairpin movement as there were no hydrogen bonds or direct contacts made between the hairpin and either the substrate-effector pair or loop 2.

**Figure 3.**
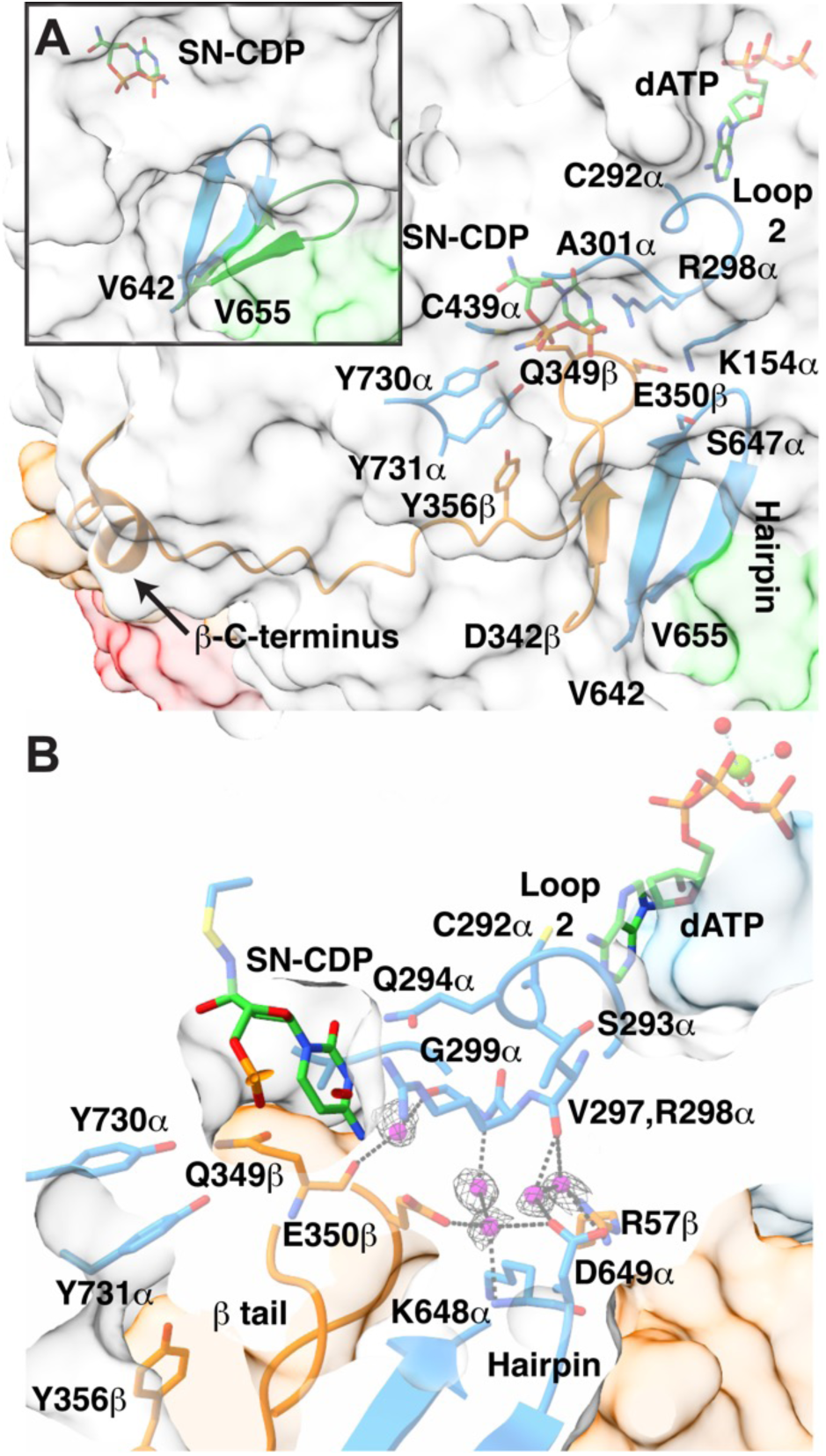
The β-tail binding pocket interactions. **(A)** The β-tail (orange ribbon) binding pocket is comprised of a hairpin (residues V642α-V655α, blue ribbons) on one side, Y730α and Y731α on the other side, and the SN-CDP adduct (green carbons) and loop 2 (C292-A301) on top. Specificity effector dATP (green carbons) is also shown. The protein surface of α is in light gray (cone domain surface in green), β surface in orange, β′ surface in red. Side chains are shown as sticks, with carbons colored orange for α and blue for β. (**Inset**) Hairpin in α shifts towards the active site (blue ribbon, this work) relative to substrate-free α2 (green ribbon, PDB 2R1R). **(B)** Water molecules (magenta spheres with gray mesh; sd level 1) mediate interactions between the β-tail (orange Cα trace), the hairpin (blue ribbons), and loop 2 (blue sticks). The SN-CDP adduct and specificity effector dATP are shown as sticks with carbons in green. Side chains colored as in A. Dashed black lines are potential hydrogen bonding interactions with distances ≤ 3.5 Å.

These structural data finally resolve this mystery. This high-resolution structure shows water molecules bridging between backbone atoms of loop 2 residues V297α and G299α and hairpin residues K648α and D649α (**Fig. 3B**), which finally explains how an ordered loop 2 stabilizes the flipped-in orientation of the hairpin and thus the β-tail binding pocket. This new structure also shows through-water contacts from loop 2 residue G299α to the β-tail (backbone of Q349β and side chain of E350β), and from E350β of the β-tail to the hairpin (**Fig. 3B**). In addition to hydrogen bonding to bridging water molecules, E350β has been observed to hydrogen bond to the side chains of K154α and S647α (20); the latter depending on the side chain conformation of the S647α (**Table S10**). Substitution of E350β has previously been shown to disrupt radical transfer between subunits (42) and now the cryo-EM data indicate E350’s key role in positioning the β-tail for radical transfer. The net result of these interactions (**Fig. 3B**) is that the binding of the β-tail is afforded when a cognate substrate-effector pair is bound and loop 2 is positioned appropriately. If a mismatched substrate-effector pair is bound, a mispositioned loop 2 won’t stabilize the β-tail binding pocket via this water network, hindering turnover on a disfavored substrate. Additionally, when substrate is reduced to product, structural data show that loop 2 is flipped out of the active site (see **Fig. S10** and above discussion of the loop 2 in α′). Such a movement of loop 2 would destabilize β2 binding, facilitating β2 departure and subsequent product release. Thus, the interactions shown in **Fig. 3B** not only provide a means by which substrate-effector binding enables β2 binding, these interactions also provide a means to trigger β2 release.

### N_3_CDP-inhibited structure provides view of intact wild-type PCET pathway that includes water molecules

The goals of determining a higher resolution structure of α2β2 were to identify the proton donor/acceptor to PCET residues, including Y356β, and to investigate the effect, if any, of using unnatural amino acids to trap the α2β2 complex. Our 2.6-Å resolution structure shows a fully intact PCET pathway with residues lined-up as previously described (**Fig. 4A**) (20). E52β, which was substituted (E52Q) in the previous structure, is similarly positioned. The distances between the residues along the PCET pathway (Y122β-[W48β]-Y356β-Y731α-Y730α-Cys439α) are also similar, except for the distances between Y356β/Y731α and Y731α/Y730α, which are shorter in the new structure due to shifts in the positions of Y356β and Y731α. (**Fig. 4A**). Excitingly, our 2.6-Å resolution structure has allows us to visualize water molecules along the PCET pathway (**Fig. 4B, S14A, Table13**). The following residues appear to be involved in stabilizing this PCET water network: R236β, D237β, E52β, V353β, Q349β, N322α, R411α, and E623α (**Fig. 4B**). Of the 20 water molecules shown along the PCET pathway, four have been previously seen in the β2 crystal structures (27, 29, 30) and nine have been previously seen in the α2 crystal structures (24–28) (**Fig. S14**). Only seven water molecules along the PCET pathway have not been previously visualized in the individual α2 or β2 subunits (**Fig. S14**), indicating that the majority of PCET water molecules remain bound when the α2β2 complex dissociates.

**Figure 4.**
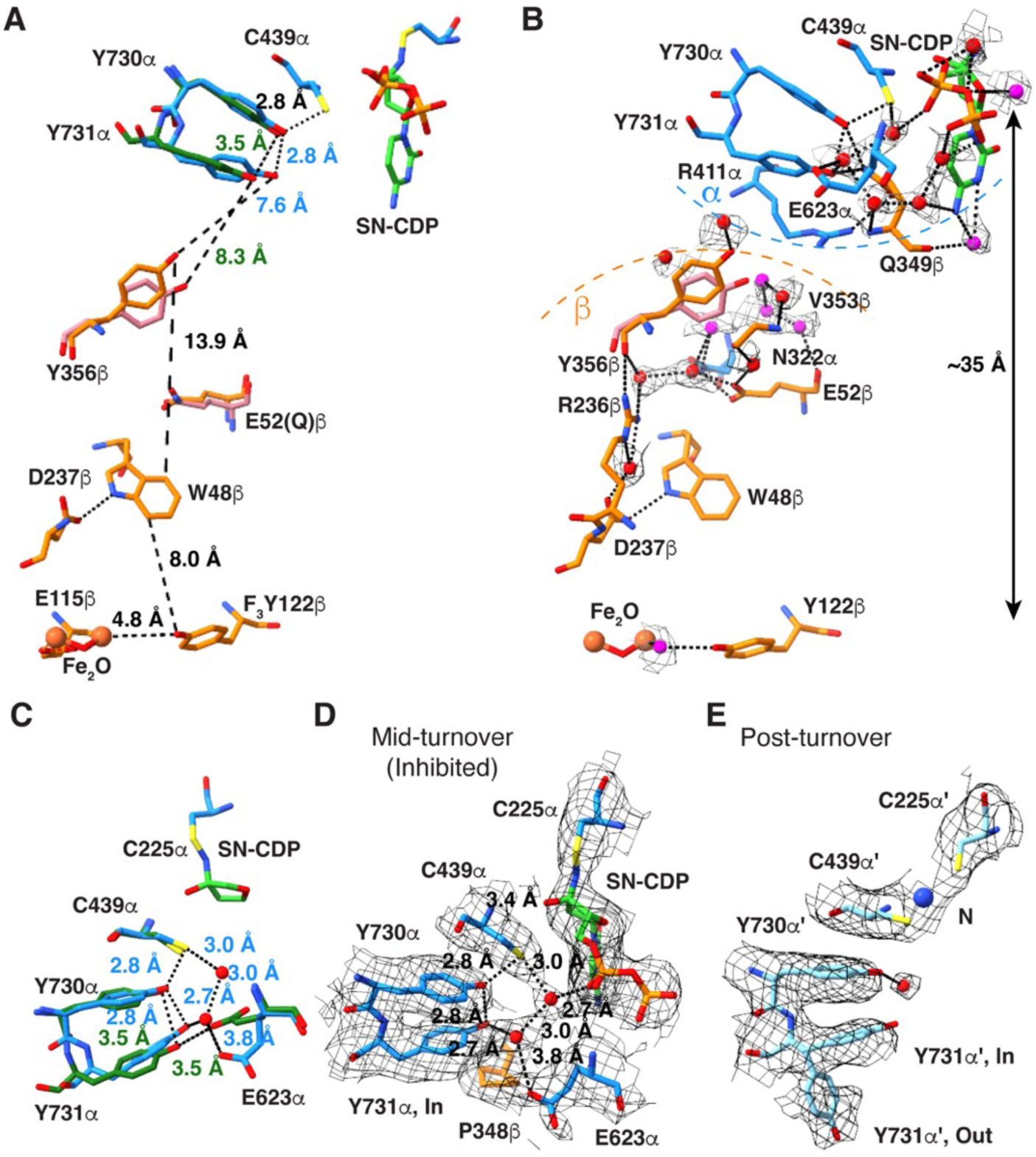
Water molecules line the PCET pathway. **(A)** The PCET pathway from this work (β in orange and α in blue) showing previous positions of E52Qβ, Y356β (dark pink) and Y731α (dark green) from the 3.6-Å resolution model (PDB 6W4X) shown. **(B)** The same PCET pathway as shown in **(A)**, but with water molecule density (gray mesh; sd level 1) and additional residues involved in hydrogen bonding interactions with the water molecules shown. Previously observed waters are red spheres (see **Tables S3-S6**) and newly observed waters are magenta spheres. Only the ribose ring of SN-CDP is shown for clarity. Select hydrogen bonds are indicated by dashed black lines (distance ≤ 3.5 Å) with additional interactions shown in **Fig. S14**. (**C**) Superimposition of Y731α, Y730α, and E623α from the previous structure (PDB 6W4X) (dark green) with the current structure (dark blue). Distances ≤ 3.8 Å are labeled in dark green (previous structure) and blue (current structure). Both water molecules, Water1027 (upper) and Water1014 (lower), have been observed previously. **(D)** The active site of α, which is inhibited mid-turnover by N_3_CDP. P348 of the β tail is shown with carbons in orange, α residues with carbons in dark blue, SN-CDP with carbons in green, and water molecules Water1027 (upper) and Water1014 (lower) as red spheres. Black dashed lines indicate potential hydrogen bonds (distance ≤ 3.8 Å). Cryo-EM density is shown as gray mesh (threshold set to sd level 3). **(E)** The active site of α′, with cryo-EM density for both the “flipped in” and “flipped out” conformations of Y731 (threshold set to sd level 3). The N-adduct in the active site is modeled as a single nitrogen atom (blue sphere) and the water molecule is shown as a red sphere.

For turnover to occur, the radical fires from β to α, starting with the reduction of the stable tyrosyl radical Y122β-O● to Y122β-OH. Given that there are no polar residues capable of proton donation near Y122β, it has been proposed that an Fe-bound water molecule is the proton donor (11, 43). However, poor density around the diiron center has hindered the ability to visualize a bound water in the *E. coli* class Ia RNR. Notably, here we see density for water bound to Fe1 that is in hydrogen bonding distance (3.2 Å) to the hydroxyl group of Y122β (**Fig. 4B, S15**). The next PCET step is either oxidation of W48β or Y356β. Definitive support for a Trp radical intermediate in the PCET pathway has not been obtained, but the structural positioning of W48 midway between Y122β and Y356β is suggestive of a role in radical transfer (20). Y356β has been well studied (11) and is known to form a transient Y356β-O● radical species. A key question has been the source of the proton acceptor/donor that allows for Y356β-OH to be oxidized to Y356β-O● and then reduced back to Y356β-OH as the radical moves from β to α. The initial structure of the α2β2 complex showed no polar residue nearby. The substituted E52Qβ was closest, at 6.9 Å, raising the possibility that a wild-type structure would show a closer distance between E52β and Y356β. However, this is not the case. E52β is positioned similarly to E52Qβ, but Y356β has moved ~2.1-Å, away from E52β and closer to Y731α (**Fig. 4A**). Instead of a polar residue in direct contact with Y356β, water molecules are positioned to contact Y356β (**Fig. 4B, Fig. S14**), supporting the proposal that water molecules are the direct proton donor/acceptors of Y356β and thus play a direct role in PCET, consistent with electron nuclear double resonance (ENDOR) and electron paramagnetic resonance (EPR) spectroscopies and DFT studies (44–46).

### N_3_CDP-inhibited structure allows us to evaluate proposed water channels between the protein surface and PCET residue Y356β

Notably, Y356β sits at the α–β interface where we previously predicted that water channels could provide water molecules from bulk solvent to interact with Y356β (20). With our improved resolution, we can identify that there are two water channels, which we will refer to as pink and yellow channels, that connect bulk solvent with Y356β (**Fig. 5 and S16**). Additional water channels that branch from the pink and yellow channels will be described elsewhere. The pink and yellow channels lie on opposite sides of Y356β and do not intersect (**Fig. 5**). The yellow channel is long (35 Å), connecting bulk solvent near loop 2 in α to Y356β, E52β, and to the water network that goes all the way to the diiron center in β (**Fig. 5A**). The pink channel (20-Å long) links Y356β to bulk solvent without contacting other PCET residues in β. Interestingly, the position of Y356β in the pre-turnover structure, which is farther from the α interface, is positioned to contact a water molecule from the yellow channel (water594 in Chain C in **Tables S4, S13, S15, S16**), and Y356β in the mid-turnover structure, which is closer to the α interface, contacts a water molecule from the pink channel (water507 in Chain C in **Tables S4, S13, S15, S16**) (**Fig. 5C**).

**Figure 5.**
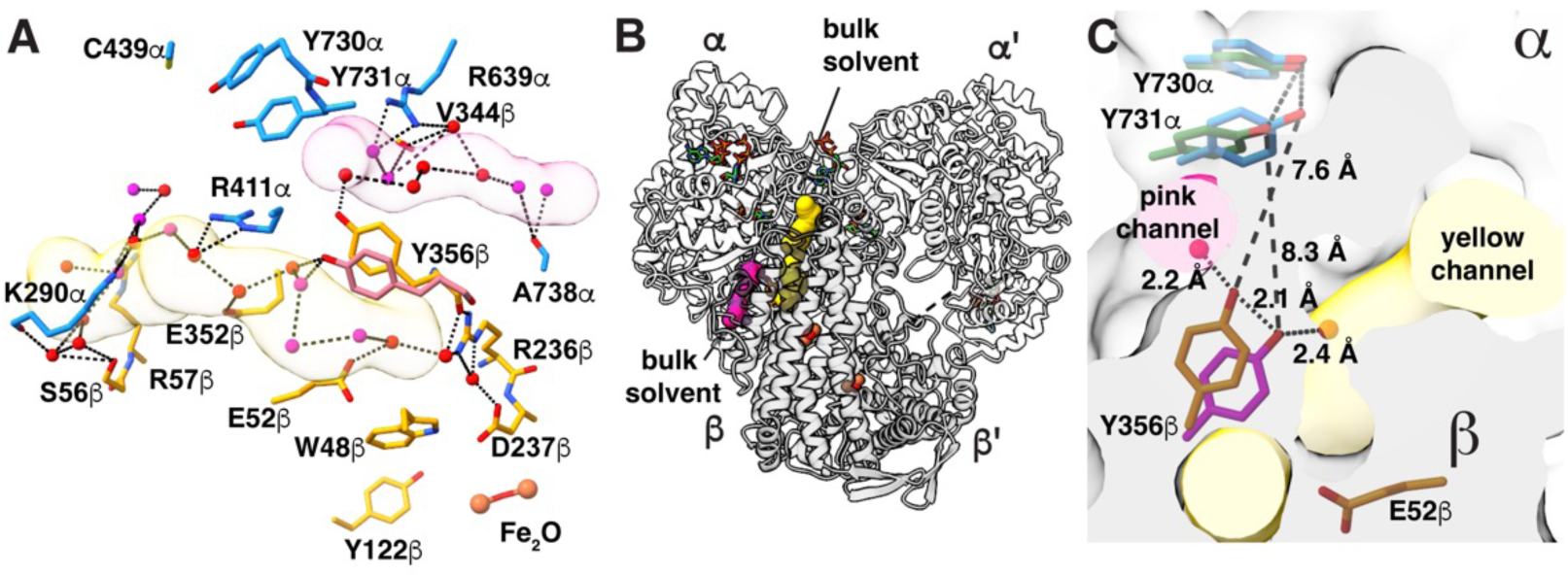
Two water channels provide access to bulk solvent from Y356β. **(A)** Two water channels originate near Y356β; pink and yellow (**Fig. S16**). Water molecules are shown as red and magenta spheres, α residues are shown with carbons in dark blue, and β residues with carbons in orange. **(B)** Ribbon drawing of the α2β2 structure indicating the location of the two water channels shown in **A**. **(C)** Overlay of Y356β/Y731α, the residues involved in PCET across the α–β interface, from the previous α2β2 structure (dark green and magenta, PDB 6W4X) and current structure presented (dark blue and orange). The α2 subunit has a light gray surface and the β2 subunit has a dark gray surface.

### Structure of mid-turnover PCET pathway provides insight into radical transfer across the α–β interface

Although a distance of 7-8 Å between Y356β and Y730α is short enough for facile electron transfer (ET) between subunits (47–49), it has been suggested that a side chain flip of Y731α could decrease this distance (see **Fig. 1E**) (22). Y731α is known to be flexible in the absence of β (17, 26, 45, 50). Both long (8-Å) and shorter (3-4-Å) distances have been observed spectroscopically between Y356β and a fluorinated-Y731α (45). An important caveat of this latter experiment is that it was done using a 2,3,5-F_3_Y122-substituted β2, which only fires on the α′ side (20, 21). Thus, it is unclear whether the experiment is measuring distances when the PCET pathway is intact or distances as β2 is starting to swing from α′ to α. Here, the cryo-EM maps support a single (unflipped) conformation for Y731α when β is bound to α, and support two conformations (flipped and unflipped) in post-turnover α′ (**Fig. 4C-E**). When the PCET pathway is intact, P348β resides directly below the Y731α side chain (**Fig. 4D**), restricting, if not preventing, Y731α side chain movement through van der Waals packing interactions (**Fig. S17**). Although these structural data cannot rule out the transient movement of the Y731α side chain, the design of the β-tail binding pocket with P348β directly below Y731α seems antithetical to such a mechanism.

Deprotonation of Y731α by a general base is another means to facilitate ET between Y356β and Y731α (49). In our previous pre-turnover structure (20), a putative general base, E623α was observed to be within hydrogen-bonding distance to Y731α, and computational analysis suggested that E623α may play a critical role in PCET (51). Notably, in this mid-turnover structure, we observe that the E623α side chain has flipped away from Y731α, and a water molecule (Water1014 in Chain B) has taken the place of glutamate’s carboxylate (**Fig. 4C**). A survey of previous structural data (20, 24–28) shows that the position of E623α observed in this mid-turnover structure is the one found in all structures except for the one structure of an α subunit that was trapped pre-turnover. It is intriguing that E623α has a unique conformation after the PCET pathway has formed but prior to forward RT, and that this E623α conformation appears ideal for deprotonation of Y731α.

In addition to the E623α rearrangement, our mid-turnover structure shows a slight movement of Y731α toward Y730α (**Fig. 4C**), which would bring the proton of Y730α-OH even closer to Y731α-O● and facilitate protonation of Y731α-O● by Y730α-OH. This proton transfer is expected to be coupled to ET between Y731 and Y730, formation of Y730α-O●, and thus forward RT. Movement of the E623α side chain creates space for Water1014 to occupy a position near Y731α, creating a circular hydrogen-bonding network that connects Water1014 to Water1027 in Chain B and to PCET residues Y731α, Y730α, C439α and substrate (**Fig. 4D, S18**). This circular hydrogen-bonding network limits the options for proton positioning on the PCET residue side chains due to both steric and electrostatic considerations (**Fig. S18**). To avoid unfavorably close proton-proton interactions with water molecules, the proton of the C439α side chain sulfhydryl can be expected to point toward the Y730α hydroxyl, and the proton of the Y730α hydroxyl toward Y731α (**Fig. S18**). We suspect that such a restriction of proton positioning would allow for facile colinear PCET back and forth between Y731α-Y730α-C439α-substrate (**Fig. S18**). In other words, once Y731α-O● forms, the availability of the Y730α-OH proton will help ensure forward RT (**Fig. S18A**). Likewise, once the product radical forms, the availability of the C439α proton to the product radical, and the availability of the Y730α proton to C439α will facilitate reverse RT (**Fig. S18B**). Notably, this circular water network is broken post-turnover, and a single water molecule remains, bound to Y730α (**Fig. 4E**).

## Discussion

The ability of enzymes to harness radical species to perform challenging chemical reactions continues to fascinate scientists. Although initially described for an adenosylcobalamin-dependent enzyme in the 1960s (52), the enzyme design principles involved in controlling radical chemistry are still enigmatic. Arguably the most impressive example of exquisite control is the reversible ~32-Å-long radical transfer that occurs between transiently associating subunits in class Ia RNR. Whereas most radical enzymes use a metallocofactor to directly generate a single transient substrate-based radical species, class Ia RNR uses a metallocofactor to generate a stable tyrosyl radical species and then generate one transient amino acid radical species after another to move a radical species over long distances, both in the forward direction and in reverse. At each step, the pathway must be designed such that RT continues in the correct direction, allowing for the radical to reach all the way to the active site in the α subunit before returning to its storage position in β. Additionally, the radical transfer must be gated such that the radical only fires when the preferred substrate, as determined by an allosteric specificity effector, is bound in the active site, since unbalanced deoxyribonucleotide pools lead to mutator phenotypes (53, 54). Finally, the PCET pathway must be designed so that it can be reliably assembled and disassembled on every round of turnover. How RNR meets these challenges has been the topic of many studies (2) and the focus of this work.

Here we have taken advantage of the “resolution revolution” of cryo-EM to obtain long-awaited structures of class Ia RNR in which the α2 and β2 subunits are trapped together allowing for the visualization of an intact PCET pathway. This work would not have been possible using X-ray crystallography, since our “trapped” α2β2 complexes are stable on the order of 15 minutes (21), not the length of time needed for the growth of diffraction quality crystals. Use of a doubly-modified E52Q/(2,3,5)-F_3_Y122-β2, wild-type α2, and substrate GDP and specificity effector TTP to trap an α2β2 complex led to a 3.6-Å resolution structure (20). Use of a mechanism-based inhibitor, N_3_CDP, specificity effector dATP, and activity effector ATP with wild-type enzyme has now led to a higher resolution (2.6-Å) structure. This new structure has allowed for the visualization of water molecules along the PCET pathway and of water channels at the α–β interface. It has also provided a view of a mid-turnover PCET pathway, i.e., a state in which forward RT has occurred, and a depiction of an α2β2 complex that is composed of the wild-type enzyme. These structural data have exceeded our initial goals; they have yielded unexpected insight into the workings of this fascinating enzyme.

Although our purpose in using N_3_CDP in these studies was not to study the mechanism of RNR inhibition, the resulting structure has been very informative in this regard. A hallmark of N_3_CDP inhibition of RNR is the appearance of the nitrogen-centered radical signal. Fitting of the resulting spectroscopic data suggested the formation of an SN-CDP adduct in the active site of *E. coli* class Ia RNR (39). This finding was surprising given that formation of such a species would seem to require loss of dinitrogen from an N_3_CDP radical species, migration of the third nitrogen from the 2′ to 3′ position of the ribose, and formation of a cysteine sulfur-to-nitrogen bond. These reactions are non-trivial and are not established steps in the mechanism of class Ia RNR with substrate (2). Notably, our cryo-EM data support the previously proposed SN-CDP adduct as the source of RNR inhibition by N_3_CDP.

Despite the use of a different method to trap the α2 together with β2, the same overall asymmetric complex was observed. As seen with our first trapped structure (20), α′ of the N_3_CDP-treated α2 dimer has turned over, β2 has shifted from α′ to α, and is trapped on α with the PCET pathway intact. This structure adds to the growing evidence (2) that the active complex of a class Ia RNR is asymmetric, with β2 positioned for RT with only one α subunit at a time. For each RT event, the binding of a cognate substrate-specificity effector pair to α leads to β2 association via the binding of one β-tail in a transient binding pocket formed in α. This higher resolution structure has expanded our model for how the binding of a cognate substrate-specificity effector pair is signaled to β2. Our data suggest that the binding of a cognate substrate-specificity effector pair secures loop 2 into the active site. This ordering of loop 2 leads to the ordering of water molecules that stabilize the conformation of the hairpin that forms the β-tail binding pocket. Lower resolution structures showed a correlation between substrate-effector binding and hairpin positioning, but this work provides the molecular basis for the correlation and demonstrates the importance of water molecules for α–β association.

Following RT from α′ to β′, product must leave and β2 must swing over to α. This higher resolution structure has also expanded our model of the structural features that facilitate the departure of β2 from α′ following the first turnover. As previously noted, the disulfide formation that accompanies turnover shrinks the active site (20, 25). Active site compaction should disrupt interactions between product and loop 2. We see here that loop 2 flips away from the active site on α′, adopting the same conformation that was previously observed in an oxidized crystal structure of α2 (25). The bridging water molecules are gone, and the flipped loop 2 conformation now occupies a position that would be unattainable when β is bound. It was previously noted that disulfide formation should favor product release and β2 departure (20) but the involvement of a loop 2 conformational change in β2 departure is made clear in this current structural study.

Use of wild-type enzyme in this structural study allowed for an analysis of whether the previously employed amino acid substitutions affected the local structure near the sites of the substitutions in the prior study. Although we found no noticeable differences in the position in this structure of residues Y122β (previously F_3_Y122β) or E52β (previously E52Qβ), use of wild-type β2 over the doubly modified β2, had an unexpected advantage. Wild-type β2 allows us to obtain a structure of the PCET pathway in which the radical fired but did not return. Thus, we can now compare a pre-turnover PCET pathway with a mid-turnover PCET pathway. We can also now visualize water molecules along the PCET pathway.

Structural comparisons of pre-turnover and mid-turnover structures have allowed us to suggest a modified mechanism for forward RT that does not involve the flipping of Y731α toward Y356β (**Fig. 1E**, **6**). The importance of Y356β to PCET has been well established (11, 15), but up until 2020, this residue had never been visualized. Its location on the flexible β-tail resulted in a lack of density for Y356β and twenty other amino acids of the β-tail in all structures (27, 29, 30, 55). Y356β was finally visualized using cryo-EM in 2020 when the β-tail was trapped in its binding pocket on α in the pre-turnover structure. Our current structure is only the second time that Y356β has been visualized and now we see that the side chain can adopt an alternative position. Y356β is one of the two side chain rearrangements that is observed when comparing pre- and mid-turnover states. The side chain of E623α is the other.

**Figure 6.**
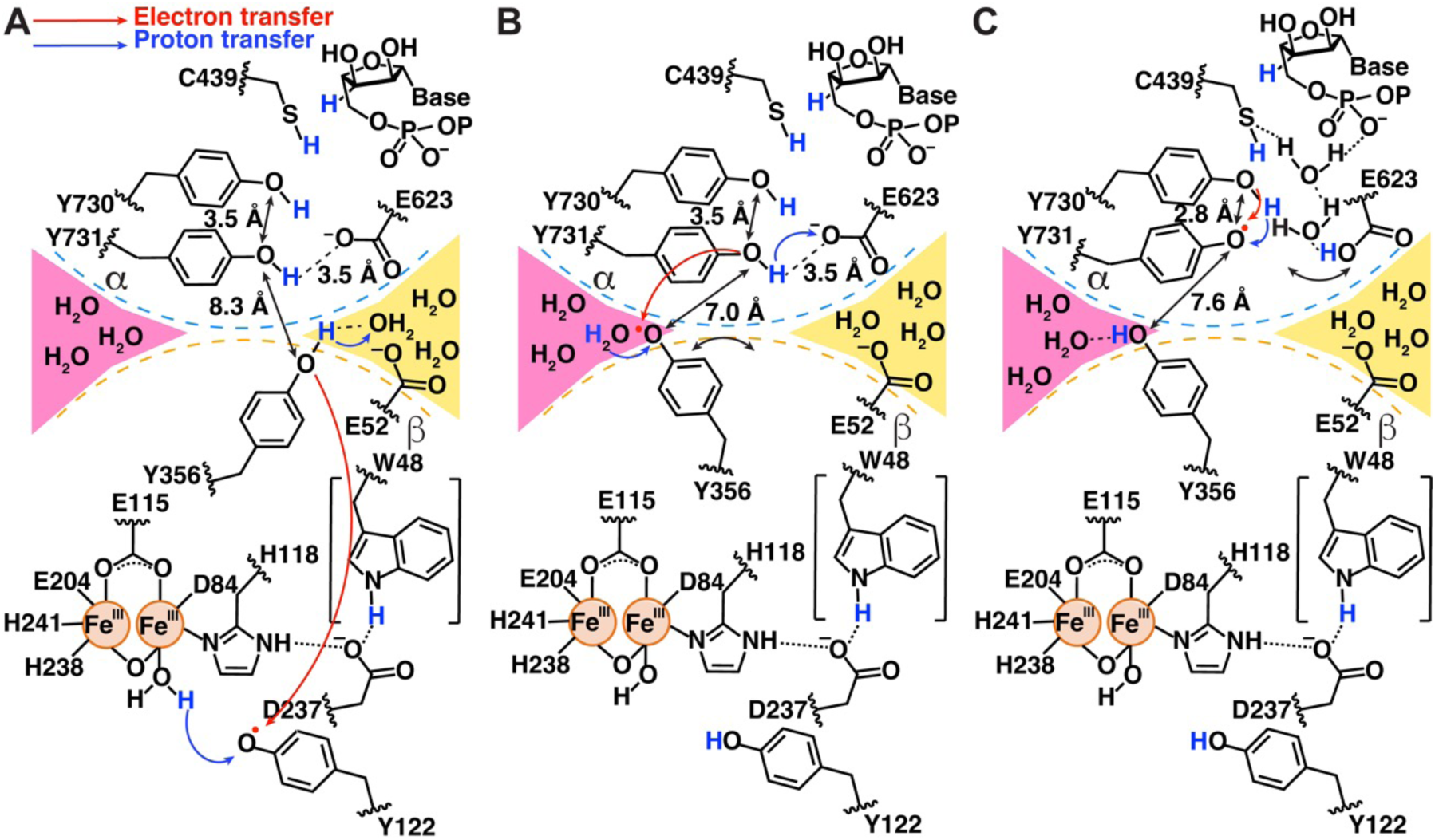
Schematic of side chain movements observed to accompany forward RT. **(A)** Scheme based on residue positions in the pre-turnover α2β2 structure (PDB 6W4X), with information about water channels from the current structure added. At the α–β interface, Y356β participates in hydrogen bonding interactions with a water molecule in the same water channel as E52β (yellow channel). **(B)** Scheme based on residue positions in the pre-turnover α2β2 structure (PDB 6W4X) except for Y356β, which is drawn as positioned in the mid-turnover structure. If Y356β undergoes the observed swinging motion before Y731α tilts closer to Y730α, the Y356β-Y731α distance would be a relatively short 7 Å. In the mid-turnover position, Y356β can hydrogen bond to water molecules in the pink channel. **(C)** Scheme based on residue positions and water molecule positions in this mid-turnover structure.

It is notable that in that pre-turnover structure, the side chain of Y356β points down toward the PCET pathway running within β, apparently positioned for forward RT starting from Y122β (**Fig. 6A**). Water molecules from the yellow channel, which also contact E52β (**Fig. 6A**), are available to serve as the proton acceptor for Y356β when Y356β is oxidized to form Y356β-O●. Loss of hydrogen bonding interactions between a yellow channel water and Y356β-O● may free Y356β-O● to swing toward the pink water channel (**Fig. 6B**). This side chain movement would be expected to favor forward RT compared to reverse RT, as the Y356β side chain rearrangement creates a longer distance between Y356β-O• and the yellow channel water proton acceptor and creates a longer distance to β residue W48 (12.1 Å becomes 13.9 Å) and a shorter distance to α residue Y731 (8.3 Å becomes 7.6 Å) (**Fig. 6B**).

The observation that there are two distinct, non-intersecting water channels available to Y356β, one to each conformation of this key PCET residue, is interesting as it potentially provides a molecular explanation for RT directionality at the α–β interface. The placement of the Y356β side chain near the pink channel in the mid-turnover structure suggests that a water from this channel is likely to be the proton donor for Y356β-O• reduction (**Fig. 6B**). Once re-protonated and reduced, Y356β can hydrogen bond to waters in the pink channel, stabilizing the side chain of Y356β in an orientation that is pointed up toward the PCET residues in α, thereby poised for reverse RT.

ET across the α–β interface in the forward RT direction is thermodynamically uphill (56) and would be facilitated by Y731α deprotonation. Here we propose that E623α is the proton acceptor for Y731α oxidation. E623α was first identified as important by Sharon Hammes-Schiffer in her computational studies of the PCET pathway that were based on the pre-turnover structure (22). E623α’s location near Y731α and Y730α suggested a role in facilitating PCET between these two residues. However, this new structure indictates that E623α flips away at some point during or after forward RT. This movement would seem to be inconsistent with a role in facilitating PCET between Y731α and Y730α, as this process occurs in both forward and reverse PCET and reverse RT has not yet occurred. Instead, the E623α side chain rearrangement suggests a different, but still critical, role for E623α. Here we propose that E623α acts as a general base for deprotonation of Y731α, favoring ET across the α–β interface and formation of Y731α-O●. In the role of general base, a benefit can be rationalized for the observed E623α rearrangement. By E623α shifting away from Y731α following deprotonation of Y731α, the transferred proton is not available to Y731α-O●, which can instead accept a proton from Y730α, favoring the continued forward direction of RT toward substrate (**Fig. 6C**). Reverse RT is likely to reset the protonation state of E623α by a through-water protonation of Y731α (**Fig. 6C**).

The proposal shown in **Fig. 6** provides an alternative mechanism for forward RT that does not require Y731α to access the α–β interface, and is one of several possibilities. Although these structural snapshots of the well studied *E. coli* class Ia RNR have offered tremendous insight into the molecular underpinnings of RNR chemistry, our work is not complete. Additional studies will be required for a more complete understanding of both forward and reverse RT. We suspect that RNR has even more surprises in store for us as we continue to dissect the workings of this fascinating enzyme.

## Materials and Methods

The α2 and β2 subunits of *Escherichia coli* class Ia ribonucleotide reductase (RNR) were prepared as previously described (57, 58). The 2′-azido-2′-deoxycytidine-5′-diphosphate (N_3_CDP) was synthesized by myosin cleavage of 2′-azido-2′-deoxycytidine-5′-triphosphate (N_3_CTP) using an adaption of a previous protocol (59). The reaction of 0.75 μM wt-α2, 3 mM ATP, and 25 μM dATP with 0.2 mM N_3_CDP in assay buffer (50 mM HEPES, 15 mM MgSO_4_, 1 mM EDTA, pH 7.6) was initiated by the addition of 1.5 μM wt-β2 and immediately transferred to an electron paramagnetic resonance (EPR) spectroscopy tube. The EPR spectra (**Fig. S3**), which were measured as described in the SI Materials and Methods, confirmed the formation of a nitrogen-based radical species upon reaction of RNR with N_3_CDP.

To prepare cryo-EM grids, Quantifoil R2/2 Cu 300 mesh grids were glow-discharged at −15 mA for 60 s. α2 (0.75 μM) was incubated with nucleotides (25 μM dATP, 3 mM ATP, and 0.2 mM N_3_CDP) in assay buffer for 2 min at room temperature. Following incubation, 1.5 μM β2 was added and the mixture was incubated for an additional 45 s at room temperature. The reaction mixture was then applied to the mesh grid (3 μL) and blotted for 5 s before plunging in liquid ethane. Cryo-EM data collection was performed at the Cryo-EM Core Facility at the University of Massachusetts Medical School at Worcester using a Thermo Fisher Titan Krios 300 kV electron microscope equipped with a Gatan GIF K2 camera. The cryo-EM data collection statistics are summarized in **Table S1**. Cryo-EM data processing was performed using the RELION-4.0 software suite (23), which was installed and configured by SBGrid (60). The workflow, FSC curve, and angular distribution plot are summarized in **Figure S4** and described in more detail in the SI Material and Methods. The 3DFSC (61) plot is shown in **Figure S5** and the 3D reconstruction parameters and statistics are summarized in **Table S1**.

The refined cryo-EM map was subjected to further sharpening by density modification in Phenix (62). The coordinates of the cryo-EM structure of the α2β2 *E. coli* class Ia RNR (PDB 6W4X) (20) were used as the starting model to dock into the final cryo-EM 3D reconstruction using UCSF ChimeraX (63). The cryo-EM map is of better quality in α than in α′, especially in the cone domain region. To build the SN-CDP adduct, we generated a parameter file for CDP using phenix.elbow (62) and manually modified it as described in the SI Material and Methods to be consistent with the spectroscopic data and the DTF model sent to us by the Bennati lab. The geometry of the final model of SN-CDP is given in **Fig. S8**, stereoviews of the SN-CDP binding compared to CDP binding is given in **Fig. S9**, and the fit of the SN-CDP adduct to the cryo-EM map is shown in **Fig. 2**. Water molecules (**Tables S3-S8, Fig. S16**) were added as described in the SI Material and Methods. The overall α2β2 model was refined by iterative rounds of real-space refinement using phenix.real_space_refine and the model quality was evaluated using MolProbity (64). The refinement and model statistics are summarized in **Table S1**. The residues visualized in the structure, of a total of 761 residues for the α subunit and 375 residues for the β subunit, are listed in **Table S2**. Figures were created with UCSF ChimeraX (63).

### Data, Material, and Software Availability

Coordinates and EM data have been deposited in the protein data bank (PDB) (PDBID 9DB2) (65), the electron microscopy data bank (EMDB) (EMD-46711) (66), and the electron microscopy public image archive (EMPIAR) (EMPIAR-12249) (67). All other data are included in the manuscript and/or SI Appendix.

## Acknowledgments

The authors gratefully acknowledge a grant from the Massachusetts Life Sciences Center for funding the Titan Krios at the Cryo-EM Core Facility at UMass Chan Medical School, which was used to collect the data cited in this study. The authors also thank Andreas Meyer and Prof. Marina Bennati for providing the coordinates of the DFT-modeled SN-CDP intermediate. We thank Dr. Edward Brignole, formerly assistant director of the cryo-EM facility at MIT, for his assistance with grid preparation, screening, and data processing.

## Author Contributions

CC prepared the protein samples and obtained the EPR data under the guidance of JS and DGN; AK prepared the N_3_CDP; PRK and GK carried out cryogenic-electron microscopy grid preparation, data collection and processing; PRK performed initial model building and refinement; DEW carried out additional data processing, model building and refinement; DEW and CLD performed the structural analysis and wrote the manuscript with edits from all authors.

