## Supplemental Information for "2.6-Å resolution cryo-EM structure of a class Ia ribonucleotide reductase trapped with mechanism-based inhibitor N3CDP"

### Supporting Information for
#### 2.6-Å resolution cryo-EM structure of a class Ia ribonucleotide reductase trapped with mechanism-based inhibitor $N_3$CDP

Dana E. Westmoreland[2†], Patricia R. Feliciano[1,2,3, †], Gyunghoon Kang[1,2,3], Chang Cui[3,4], Albert Kim[3], JoAnne Stubbe[2,3], Daniel G. Nocera[4], and Catherine L. Drennan[1,2,3]*

[1]Howard Hughes Medical Institute, Massachusetts Institute of Technology, Cambridge, MA 02139.
[2]Department of Biology, Massachusetts Institute of Technology, Cambridge, MA 02139.
[3]Department of Chemistry, Massachusetts Institute of Technology, Cambridge, MA 02139.
[4]Department of Chemistry and Chemical Biology, Harvard University, Cambridge, MA 02138

[†]Authors contributed equally.

*Catherine L. Drennan

**This PDF file includes:**

Supplementary Material and Methods
Figures S1 to S18
Tables S1 to S16
SI References

#### Supplementary Materials and Methods

##### Materials

For protein expression, purification, and spectroscopical expriments, Luria broth, phenylmethylsulfonyl fluoride (PMSF), 4-(2-hydroxyethyl)-1-piperazineethanesulfonic acid (HEPES), $MgSO_4$ and β-Mercaptoethanolwere purchased from Sigma-Aldrich; isopropyl β-d-1-thiogalactopyranoside (IPTG), L-arabinose, kanamycin (Km), ampicillin (Amp), (hydroxymethyl)aminomethane (Tris), and dithiothreitol (DTT) were purchased from GoldBio; imidazole, glycerol, and EDTA were obtained from Oakwood Chemicals; DEAE, Q-sepharose, Sephadex G-25 resins were obtained from GE Healthcare Life Sciences; Ni-NTA agarose resin was purchased from Qiagen. For the Cryo-EM grids preparation, deoxyadenosine triphosphate (dATP) and adenosine triphosphate (ATP) were purchased from Sigma-Aldrich. HEPES buffer, $MgSO_4$, and EDTA were purchased from Sigma-Aldrich.

##### Protein preparation

The α2 and β2 subunits of *Escherichia coli* class Ia ribonucleotide reductase (RNR) were prepared as previously described (1, 2). The protein concentrations were determined using the extinction coefficients at 280 nm of 189 $mM^{-1}$ $cm^{-1}$ for the α2 dimeric subunit and 131 $mM^{-1}$ $cm^{-1}$ for the β2 dimeric subunit (3).

##### Synthesis of 2'-azido-2'-deoxycytidine-5'-diphosphate ($N_3CDP$)

The 2′-azido-2′-deoxycytidine-5′-diphosphate ($N_3CDP$) was synthesized by myosin cleavage of 2′-azido-2′-deoxycytidine-5′-triphosphate ($N_3CTP$) using an adaption of a previous protocol (4). $N_3CTP$ was purchased from Trilink Biotechnologies and myosin from rabbit muscle was purchased from Sigma-Aldrich. To synthesize $N_3CDP$, 1 mM $N_3CTP$ was treated with 0.5 units of myosin in digestion buffer (25 mM potassium borate, pH 9, 5 mM $CaCl_2$, 50 mM KCl) at 25 °C for 30 min. Myosin was then removed by centrifugation using an Ultracel-10 KDa centrifugal filter (EMD Millipore). The filter was washed twice with 250 µL digestion buffer to maximize recovery. The flow-through was combined and loaded onto a DEAE Sephadex A-25 column (15 mL column volume) pre-equilibrated with 150 mL of 50 mM triethylammonium bicarbonate (TEAB) buffer (pH 7.6) at 4°C. The column was washed with 200 mL of 50 mM TEAB buffer and eluted with a linear gradient of 50-600 mM TEAB buffer (150 mL each buffer; pH 7.6). Fractions of 4 mL were collected and the fractions with absorption at 271 nm were combined and lyophilized. The product was redissolved in water and characterized by $^1H$ and $^{31}P$ nuclear magnetic resonance spectroscopy. The final concentration of $N_3CDP$ in solution was determined using the extinction coefficient at 271 nm of 9100 $M^{-1}$ $cm^{-1}$.

##### *E. coli* class Ia RNR reaction with $N_3CDP$

The reaction mixture containing 0.75 µM wt-α2, 3 mM ATP, 25 µM dATP and 0.2 mM $N_3CDP$ in assay buffer (50 mM HEPES, 15 mM $MgSO_4$, 1 mM EDTA, pH 7.6) was incubated in a 25 °C circulating water bath for 1 min. The total volume was 200 µL. The reaction was initiated by the addition of 1.5 µM wt-β2, immediately transferred to an electron paramagnetic resonance (EPR) spectroscopy tube in a 25 °C circulating water bath and manually quenched in liquid isopentane cooled by liquid nitrogen after 45 s. The EPR spectra were collected at 77 K on a Bruker EMX X-band spectrometer equipped with a liquid helium cryostat. The spectra (**Fig. S3**) were measured with a 9.4 GHz microwave frequency, a 32 µW power, a 1 G modulation amplitude, a 100 kHz modulation frequency, a 30 s scan time, and averaged over 128 scans.

##### Cryo-EM grid preparation

Quantifoil R2/2 Cu 300 mesh grids were glow-discharged at -15 mA for 60 s using a PELCO easiGlow cleaning system and subsequently used for cryo-EM grid preparation. To promote the formation of the α2β2 complex, 0.75 µM α2 was incubated with nucleotides (25 µM dATP, 3 mM ATP, and 0.2 mM $N_3CDP$) in assay buffer (50 mM HEPES pH 7.6, 15 mM $MgSO_4$, and 1 mM EDTA) for 2 min at room temperature. Following incubation, 1.5 µM β2 (2-fold excess to maximize complex formation) was added to initiate the reaction and the mixture was incubated for an additional 45 s at room temperature. The reaction mixture was then applied to the mesh grid (3 μL) and blotted for 5 s (Whatman filter paper #1) at 10 °C and 95 % humidity. The grid was then plunged in liquid ethane using a Thermo Fisher Scientific Vitrobot (Mk IV) cryo-plunger. Grids were stored in liquid nitrogen until data collection. The final protein solution contained 0.75 μM α2, 1.5 μM β2, 25 μM dATP, 3 mM ATP, 0.2 mM $N_3$CDP, 50 mM HEPES, pH 7.6, 15 mM $MgSO_4$, and 1 mM EDTA.

**Cryo-EM data collection**

Cryo-EM data collection was performed at the Cryo-EM Core Facility at the University of Massachusetts Medical School at Worcester using a Thermo Fisher Titan Krios 300 kV electron microscope equipped with a Gatan GIF K2 camera. The data collection parameters used were a defocus range of -1 to -2.3 μm, magnification of 105,000 X, with two shots per hole and pixel size of 0.415 Å (super-resolution, unbinned), 30 frames, and 1.6982 $e^-/Å^2$/frame dose. The collected data set contained 6,007 movies. The cryo-EM data collection statistics are summarized in **Table S1**.

**Cryogenic-electron microscopy data processing.**

The frames of each dose-fractionated image stack were corrected for beam-induced motion using MotionCor2 (5) within the RELION-4.0 software suite (6). The contrast transfer function (CTF) parameters of each motion-corrected micrograph were estimated using CTFFIND 4.1 (7) implemented in the RELION-4.0 software suite (6). Micrographs with a CTF fit resolution greater than 5 Å were discarded from further analysis. Based on this parameter, the data set containing 6,007 movies was truncated to 5,973 micrographs. These micrographs were used for particle selection using Topaz (8) implemented within the RELION-4.0 software suite (6). Initially, 6199 particles were automatically picked from a randomly generated subset of 20 micrographs using Laplacian-of-Gaussian picking within RELION-4.0 (particle diameter range 150 – 190 Å, box size of 576 pix downscaled to 288 pix). The automatically picked particles were classified into 50 2D class averages to exclude non-particle images from Topaz training. Nine of these 2D classes containing 2070 particles were used to train a neural network automated pipeline for particle picking in Topaz within the RELION-4.0 suite, which picked 1,029,442 particles (particle diameter of 170 pix, box size of 576 pix downscaled to 288 pix) from the 5,973 micrographs using a cut-off of -2.25 for positive signal (**Figure S4**).

These 1,029,442 particles were sorted into 200 2D class averages (mask diameter of 180 Å) (**Figure S4**). Of these, a total of 760,117 α2β2-like particles from 57 2D classes were used to generate an *ab initio* 3D model using no imposed symmetry (i.e. C1) and two classes in RELION-4.0. This set of 760,117 α2β2-like particles was used for 3D classification into four classes (**Figure S4**) using the 558,872-particle class of the two-class *ab initio* 3D model as a reference (**Figure S4**). Class 1 (440,549 particles) displayed an α2β2-like architecture with secondary structure features and the particles from this class following re-extraction at full scale (box size 576 pix) were used for unmasked reference-free 3D auto-refinement yielding a 3.4-Å resolution 3D reconstruction. To further improve map quality, a mask was generated and used to perform post-processing (map sharpening) yielding a 3.1-Å resolution 3D model. Following map sharpening, CTF refinement of the anisotropic magnification followed by the per-particle defocus was performed in RELION-4.0. Bayesian polishing was also performed in RELION-4.0. The refined particle set (440,549 particles) was used to re-run 3D auto-refinement and post-processing, yielding a final reconstruction at 2.6-Å resolution (FSC = 0.143 (**Figures S4** and **S5**)). The local resolution map was estimated using RELION-4.0 (6) (**Figure S6**). The 3D reconstruction parameters and statistics are summarized in **Table S1**.

**Model building, refinement, and validation**

The refined cryo-EM map was subjected to further sharpening by density modification in Phenix (9) using the two unfiltered half maps from the final round of 3D-refinement, a soft mask, a resolution of 2.6 Å and a molecular weight of 260690.0 Da. The coordinates of the cryo-EM structure of the α2β2 *E. coli* class Ia RNR (PDB 6W4X) (10) were used as the starting model to dock into the final cryo-EM 3D reconstruction using UCSF ChimeraX (11). The model was then fit into the map using phenix.dock_in_map. Since the cryo-EM structure of α2β2 *E. coli* class Ia RNR (PDB 6W4X) (10) did not have ATP molecules bound in the cone domains, coordinates from a two ATP bound crystal structure of α2 (PDB 8VHN) (28) were used to guide the cone domain model building. The cryo-EM map is of better quality in α than in α′. The α2β2 model was refined by iterative rounds of real-space refinement using phenix.real_space_refine and the initial round of real space refinement included simulated annealing to minimize model bias (9). The geometry restraints for ATP and dATP were generated by phenix.elbow. The cryo-EM map and model were validated using phenix.validation_cryoem (9), and the model quality was evaluated using MolProbity (12). The refinement and model statistics are summarized in **Table S1**. The residues visualized in the structure, of a total of 761 residues for the α subunit and 375 residues for the β subunit, are listed in **Table S2**. Figures were created with UCSF ChimeraX (11). Software packages were compiled by SBGrid (13).

**SN-CDP Modeling and Refinement**

Bennati and coworkers modeled their spectroscopic data of the nitrogen-based adduct (the "SN-CDP" adduct) formed during $N_3$CDP inhibition of RNR using density function theory (DFT) (14). This DFT model of the SN-CDP adduct had three simplifications: the 5′ diphosphate group was approximated by a hydroxyl group; the 2′ cytidine base was approximated as a primary amine; and the cysteine residue was approximated as a $SCH_3$ moiety **(Figure S8)**. Due to these simplifications, we could not exclusively use the DFT coordinates, which we obtained from the Bennati lab, to generate a parameter file for refinement of SN-CDP. Therefore, we generated a parameter file for CDP using phenix.elbow (9) and manually modified it to be consistent with the spectroscopic data and the DTF model. An initial round of real space refinement in phenix without restraints was used to place SN-CDP into the cryo-EM map. A second round of refinement was run restraining the distance between the nonbonded atoms N3′ and C1′ to 3.399 ± 0.010 Å and N3′ and C4′ to 2.425 ± 0.010 Å to be consistent with the N3′-H1′ and N3′-H4′ experimental distances obtained by electron spin echo envelope modulation (ESEEM) spectroscopy (**Fig. S8B, D**) (15, 16). The N3′ to Cys225 S distance was restrained to 1.634 ± 0.050 Å (**Fig. S8B**), consistent with the DFT model, in which Cys225 is approximated by a methyl sulfide (**Fig. S8A,B**) (14). The position of the Cys255 side chain, which is in a standard rotamer conformation, is not the same as the methyl sulfide position (**Fig. S8E**). Modeling of the Cys225 side chain to be positioned like the methyl sulfide, results in a non-standard rotamer for cysteine and places the side chain out of the density. Given the difference between a cysteine side chain and a methyl sulfide, we allowed angles involving the cysteine sulfur, C3′-N3′-S-Cys225 and N3′-S-βC-Cys225, to deviate from the angles for the DTF model with the methyl sulfide substituting for Cys225 (**Fig. S8D**). Given that the DFT calculations indicated that a range of dihedral angles for both S-N3′-C3′-C4′ and βC-S-N3′-C3′ could explain the spectroscopic data, no dihedral restraints were employed. The geometry of the final model of SN-CDP is given in **Fig. S8**, stereoviews of the SN-CDP binding compared to CDP binding is given in **Fig. S9**, and the fit of the SN-CDP adduct to the cryo-EM map is shown in **Fig. 2**.

**Water Molecule Modeling**

Water molecules were added automatically using Coot (17) at an rmsd level of 1.4 and a distance range of 2.2 – 3.5 Å to account for the average hydrogen bonding distance of water (18). These water molecules were inspected manually and the water molecules that were added to regions with smeared density were removed. Additional waters were added manually throughout the structure based on the following criteria: a density threshold of 0.235; 1.67 sigma and a hydrogen bonding distance of 2.2 – 3.5 Å to a polar residue or another water molecule. The majority of the water molecules identified were previously observed in crystal structures of α2 (19-23) or β2 (22, 24, 25) (see **Tables S3-S6**). Water molecules identified automatically by Coot but not observed in a crystal structure are shown in **Table S7**, and water molecules added manually but not observed in a crystal structure are shown in **Table S8**. To further validate the modeled water molecules, we ran Q-score analysis (26), which indicated an expected Q-score of 0.7674 (out of 1) at 2.6 Å resolution as the value below which water molecules may not be reliably resolvable. 198 of the 674 water molecules were below this expected Q-score. We chose to retain all water molecules below this threshold that had a corresponding water molecule in the crystal structures because visual inspection indicated that the Q-score analysis was under-identifying water molecules that are conserved, as has been previously observed (26). We re-examined all waters below a Q-score of 0.7674 that did not have a correlate in the known crystal structures and removed 33 waters that had a Q-score of 0.72 or below. For complete transparency, we have included all Q-score values in **Tables S3-S8**. The final number of water molecules in our structure is 624, including the 6 water molecules associated with $Mg^{2+}$ ions. Of the PCET pathway waters, only Water 511 in Chain C (the Fe1-bound water) is significantly below the Q-score threshold and, with the single exception of Water 918 in Chain B, which has been previously observed in three crystal structures (**Table S4**), all remaining PCET pathway waters are above our designated cutoff of 0.72. All water molecules within the described water channels are above this 0.72 cutoff.

**Water Channel Analysis**

Water channels in α2β2 were identified using Caver 3.0.3 (27). The minimum probe radius was set to 1.5 Å, which is 0.1 Å larger than the Coulombic radius of water, but 0.2 Å smaller than the van der Waals radius of water (28), and the desired radius was set to 2.2 Å, which is the minimum radius of a hydrogen bond to water (29). The shell depth was set to 5 Å, the shell radius was set to 3 Å, and the clustering threshold was 3.5. All protein residues and nucleotides were included in the calculation. For the yellow water channels (**Fig. S16A, B**), the starting point optimization was set to within 4 Å of Y356β. For the pink water channel (**Fig. S16C**), the starting point optimization was set to within 3 Å of the water that forms a 2.2-Å hydrogen bond with Y356β (**Fig. 4D** and **5C**, orange).

**Surface Area Analysis**

PDBePISA (30) was used to determine the surface area between individual subunit chains by uploading the structural model and selecting the interfaces option in the web interface. For the surface area of β2 buried by the cone domain of α2, residues 96-745 of chain B were manually deleted and the truncated model was uploaded to PDBePISA.

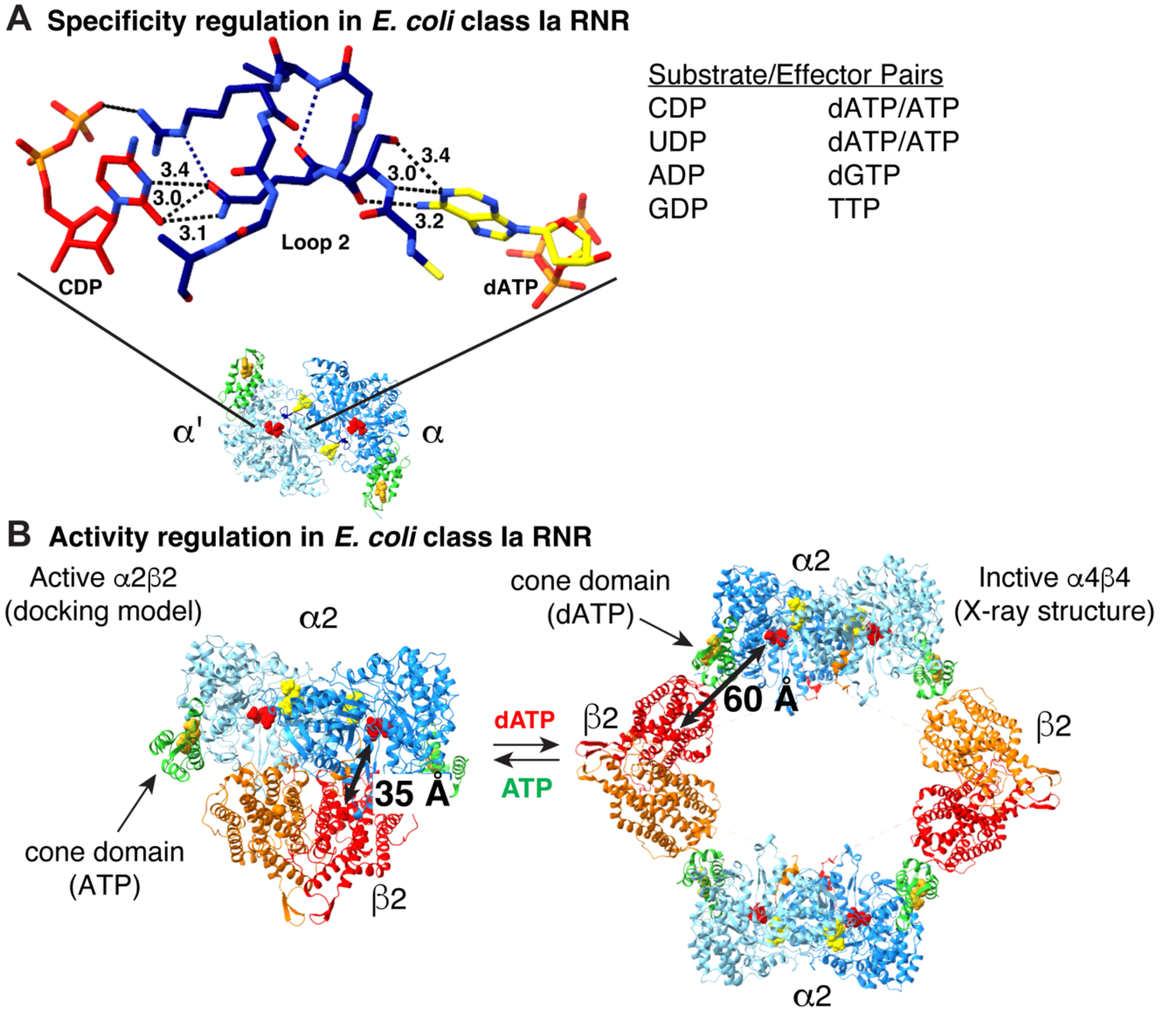


**Figure S1. Molecular basis of allosteric regulation of substrate specificity and enzyme activity in *E. coli* class Ia RNR. (A)** Preference for the four substrates of class Ia RNR (UDP, CDP, ADP, GDP) is modulated by allosteric specificity effectors (dATP/ATP, GTP, TTP) via a loop at the dimer interface called loop 2. Structure of the $\alpha 2$ subunit of *E. coli* class Ia RNR is shown with substrate (red carbons), specificity effector (yellow carbons), and loop 2 (navy blue carbons). The substrate/effector pairs are listed. **(B)** Overall RNR activity is regulated by allosteric activity effectors dATP (down-regulation) and ATP (up-regulation). At low dATP concentrations, dATP acts predominantly as a specificity effector to promote the reduction of CDP and UDP. At higher dATP concentrations, dATP binds to the activity site in the cone domain (green, residues 5-95) inducing the formation of an $\alpha 4\beta 4$ ring (PDB 5CNS), which is an inactive state of the enzyme. In this ring structure, the radical subunit, $\beta 2$, is held away from $\alpha 2$ so that the ~35-Å distance that the radical needs to travel is now ~60 Å and the route would go through solvent. Since no radical transfer can occur when $\beta 2$ is in an $\alpha 4\beta 4$ ring, the $\alpha 4\beta 4$ state is inactive. When ATP levels rise and ATP displaces dATP in the cone domain, the ring structure breaks apart and enzymatic activity is restored.

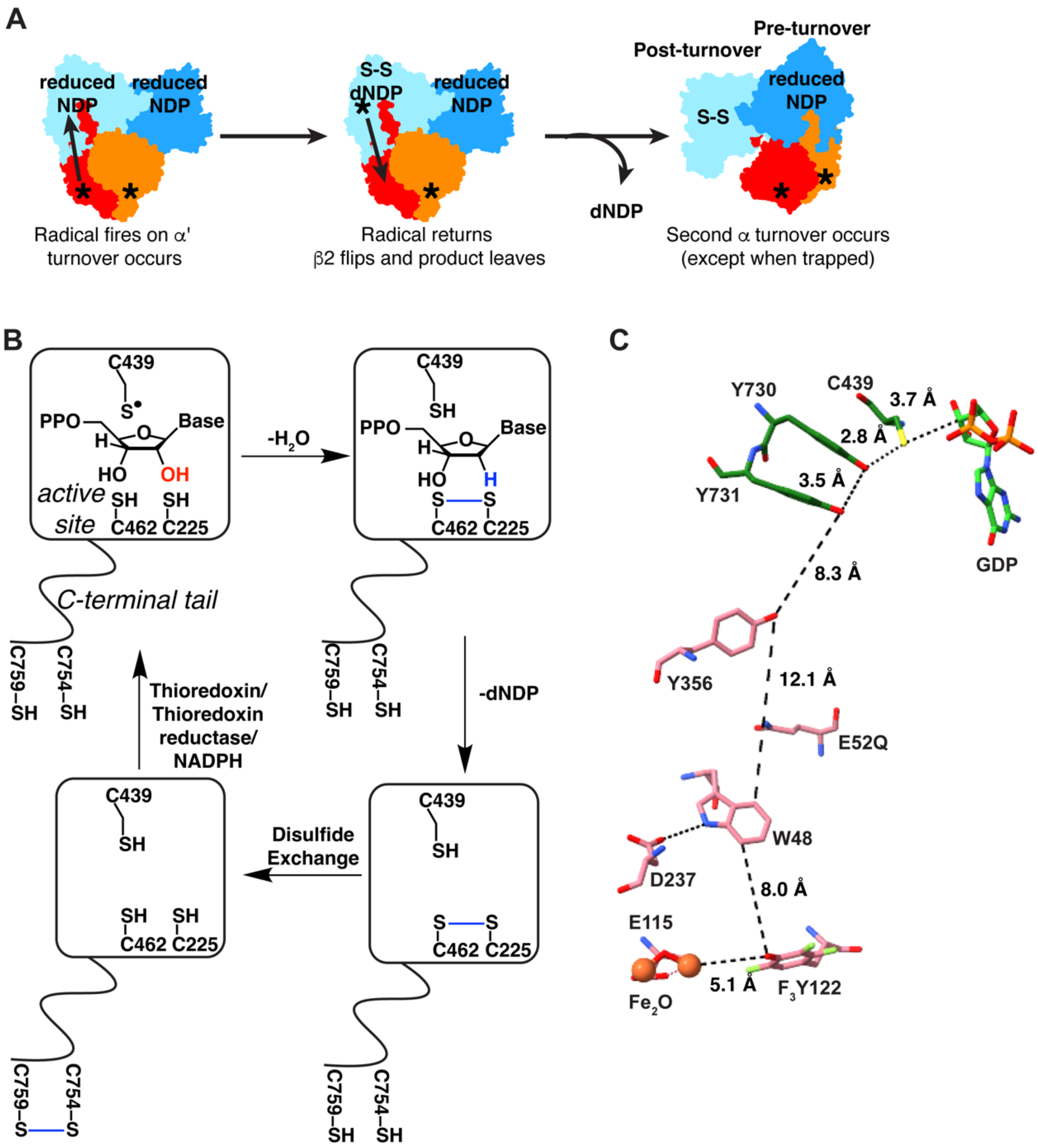


**Figure S2.** ***E. coli*** **class Ia RNR catalysis involves half-sites reactivity, PCET, and active site re-reduction. (A)** Schematic depiction for the half-sites reactivity of class Ia RNR that led to the previous 3.6-Å resolution cryo-EM structure. **(B)** Concomitant with nucleotide reduction in class Ia RNR, a pair of cysteines (C225, C462) in the active site are oxidized. This disulfide in the active site is re-reduced through two disulfide exchange steps, where another pair of redox-active cysteines (C754, C759) on the C-terminus of the α subunit reduces the disulfide in the active site and is subsequently re-reduced by thioredoxin/thioredoxin reductase/NADPH. **(C)** The PCET pathway of the previously published α2β2 structure at 3.6-Å resolution (PDB 6W4X) with wt-α2 and E52Q/(2,3,5)-$F_3$Y122-β2. The residues of the α subunit are shown with carbons in dark green and the residues of the β subunit are shown with carbons in dark pink. Interaction distances are indicated with dashed lines.

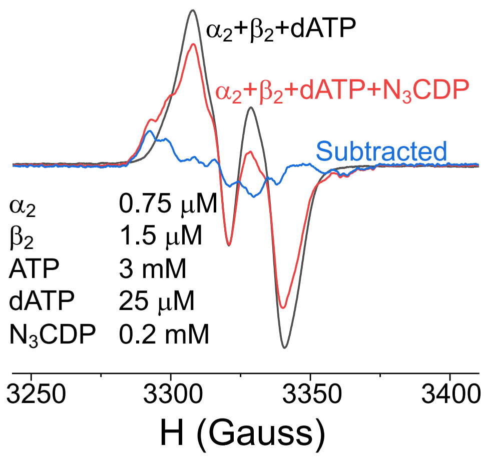


**Figure S3. Electron paramagnetic resonance spectrum before and after reaction with $N_3$CDP**. Spectrum collected using 0.75 μM α2, 1.5 μM β2, 25 μM dATP, and 3 mM ATP (black trace) and with 0.2 mM $N_3$CDP (red trace). The difference spectrum shown in blue was derived by subtracting the contribution of the unreacted Y122• from the composite spectrum (red trace). All samples were prepared in 50 mM HEPES pH 7.6, 15 mM $MgSO_4$, and 1 mM EDTA. In the absence of $N_3$CDP, the EPR signal is due to 100% Y122•. When $N_3$CDP is present, an N• signal contributes to the composite spectrum. Double integration of the composite (red trace) and difference (blue trace) suggests that the newly formed N• accounts for 25% of the total spin and the remaining 75% is attributed to Y122•.

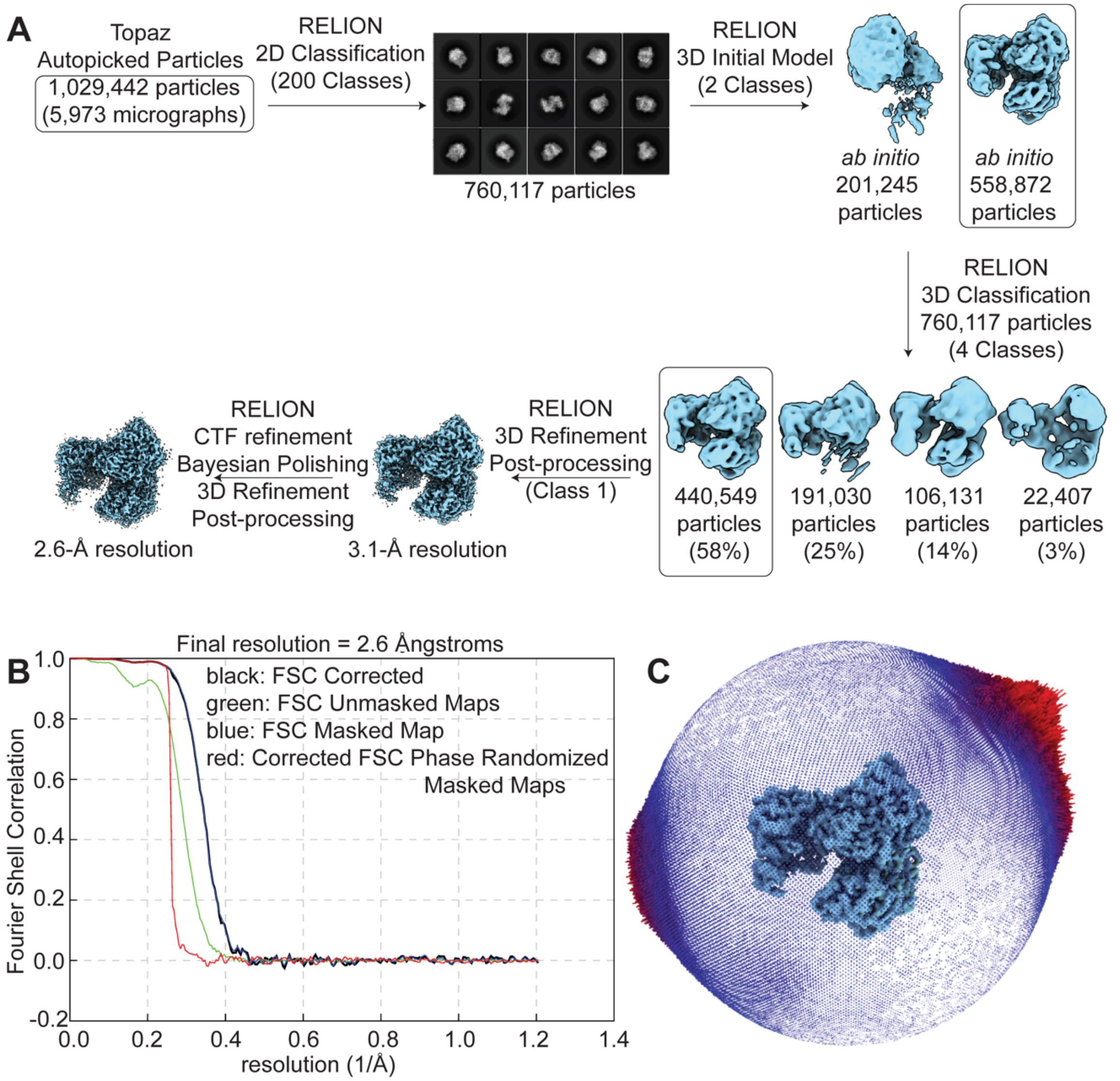


**Figure S4. Cryo-EM data processing workflow. (A)** Scheme describing the data processing steps to generate the final cryo-EM density map used for model building. See Methods for additional information and references to programs used. **(B)** Fourier Shell Correlation (FSC) plots for the final cryo-EM density map. **(C)** Particle orientation distribution for the final cryo-EM density map.

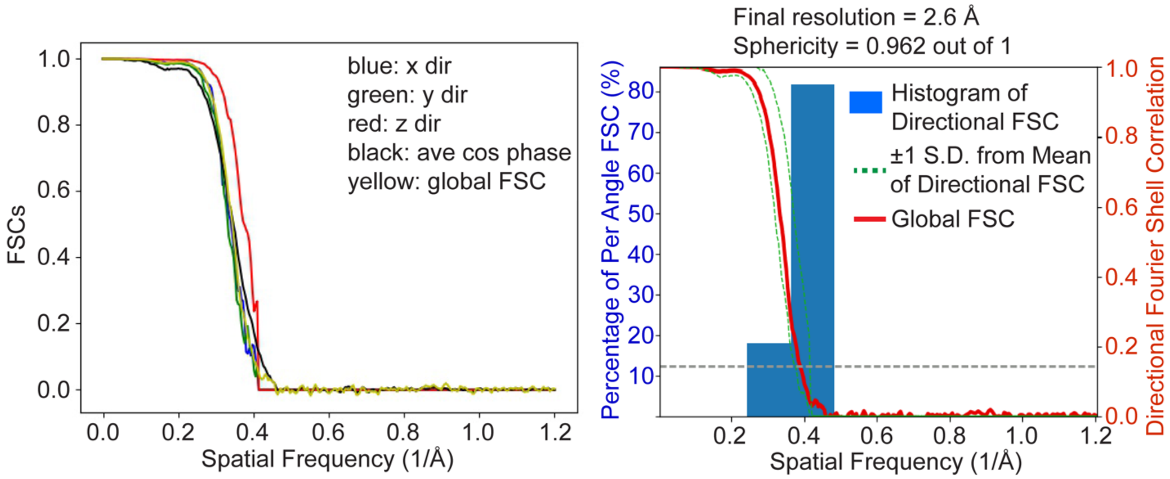


**Figure S5. The 3-dimensional Fourier Shell Correlation (3DFSC) plot for the final cryo-EM 3D reconstruction shows minimal preferred orientation in the final map.**

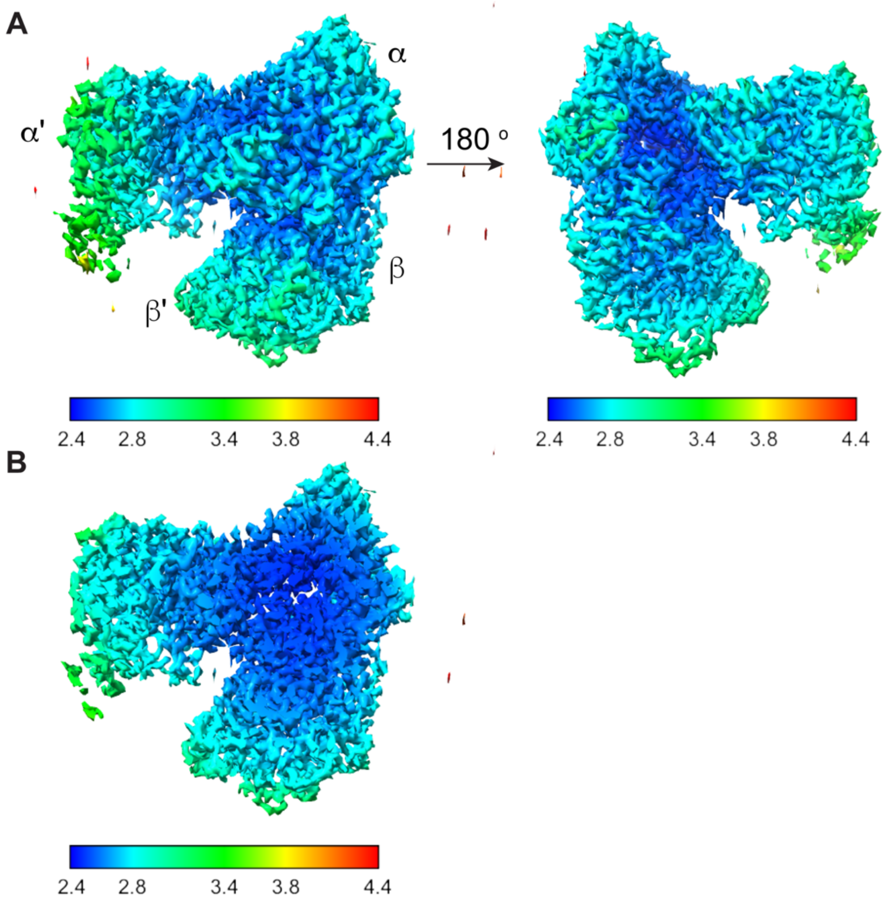


**Figure S6. Local resolution maps of the final cryo-EM density map. (A)** Two views, 180° rotations of each other, of the local resolution map of the final density map following sharpening by post-processing in RELION. The resolution ranges from 2.4 Å (blue) at the α2-β2 interface and core of the complex to 3.8 Å (yellow) at the N-terminus of one of the α monomers (α′ - chain A) to 4.4 Å in the solvent region. **(B)** Cross-sectional view of the final density map showing that the highest resolution (2.4 Å) is at the α2-β2 interface in the core of the complex.

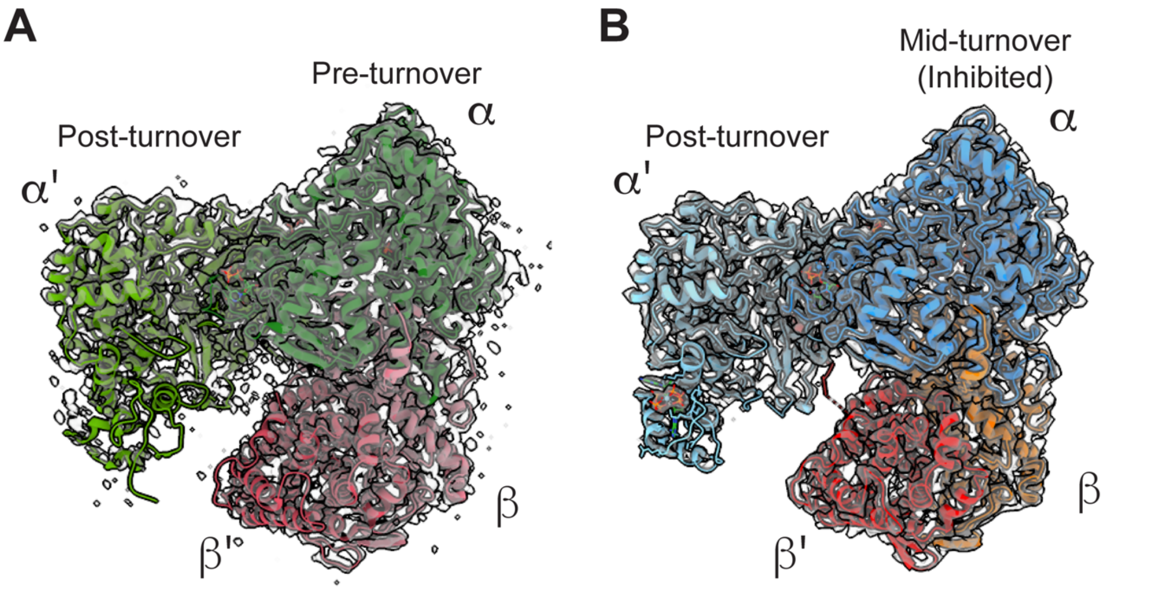


**Figure S7. Comparison of *E. coli* class Ia RNR α2β2 cryo-EM structures. (A)** The previously published 3.6-Å resolution cryo-EM map (transparent grey; sd level 5; PDB 6W4X) of the active state of α2 with E52Q/(2,3,5)-$F_3$Y122-β2 in the presence of GDP substrate and TTP specificity effector. The α2β2 model is asymmetric, where α′ (light green) and β′ (dark pink) have undergone one round of turnover and α (dark green) and β (light pink) are in a pre-turnover state which displays a fully intact radical transfer pathway. **(B)** Current structure duplicated from **Fig. 2A**.

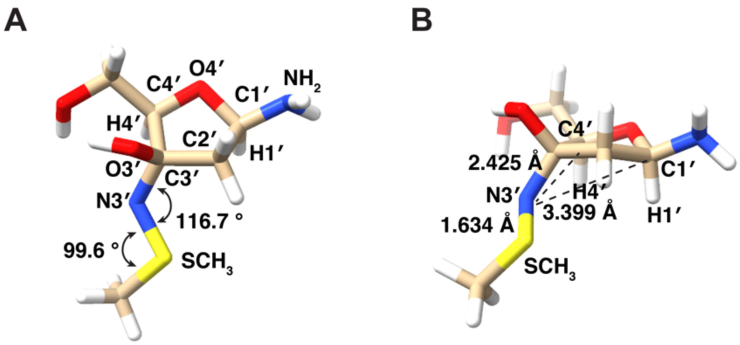


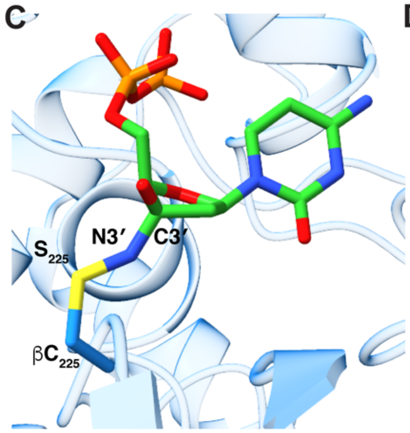


D

| Distance | Spectroscopy-Based DFT Model (Å) | Refined SN-CDP (Å) |
|---|---|---|
| S-N3′ | 1.63328 | 1.64 |
| C3′-N3′ | 1.46756 | 1.46 |
| N3′-C4′ | 2.425 | 2.42 |
| N3′-C1′ | 3.399 | 3.39 |

| Angle | Spectroscopy-Based DFT Model (°) | Refined SN-CDP (°) |
|---|---|---|
| O-C3′-N3′ | 108.2247 | 108.23 |
| C3′-N3′-S | 116.6424 | 142.31 |
| N3′-S-βC | 99.6103 | 118.60 |

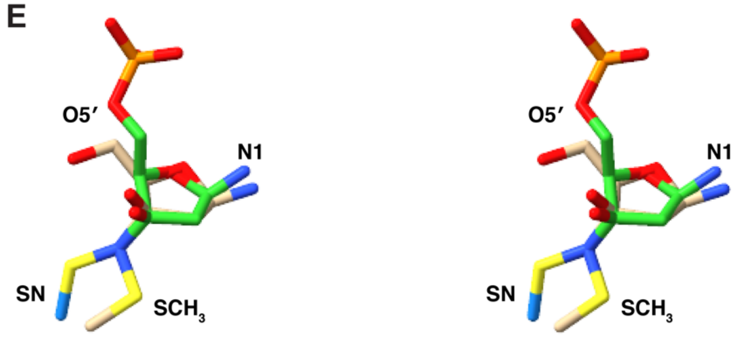


**Figure S8. Comparison of spectroscopy-based density functional theory (DFT) model of adduct and cryo-EM model of SN-CDP adduct. (A)** Top-down view of the DFT model of the SN-CDP intermediate. To simplify the DFT calculations, the 5′ diphosphate group of SN-CDP was approximated as a hydroxyl group, the 2′ cytidine base was approximated as a primary amine, and the cysteine was appropriately by a $SCH_3$ moiety (14). **(B)** Side view of the DFT model of SN-CDP. **(C)** Corresponding side view of the SN-CDP adduct following refinement into the experimental cryo-EM density. **(D)** Comparison between the distances (top) and angles (bottom) for the spectroscopy-based DFT model and the SN-CDP adduct as refined into the experimental cryo-EM density. **(E)** Stereoview of a superimposition of the spectroscopy-based DFT model (tan carbons) and the SN-CDP adduct as refined into the experimental cryo-EM density (green carbons).

**A** **Previous structure dATP:CDP**

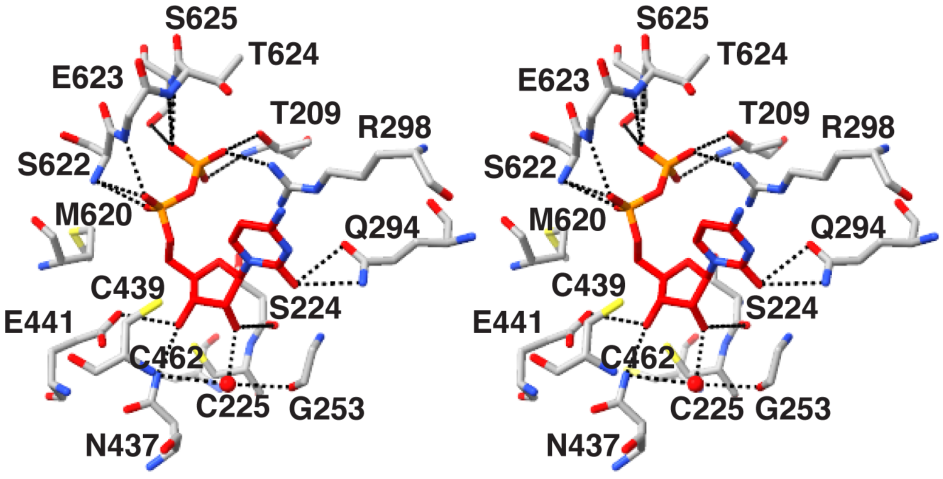


**B** **SN-CDP adduct**

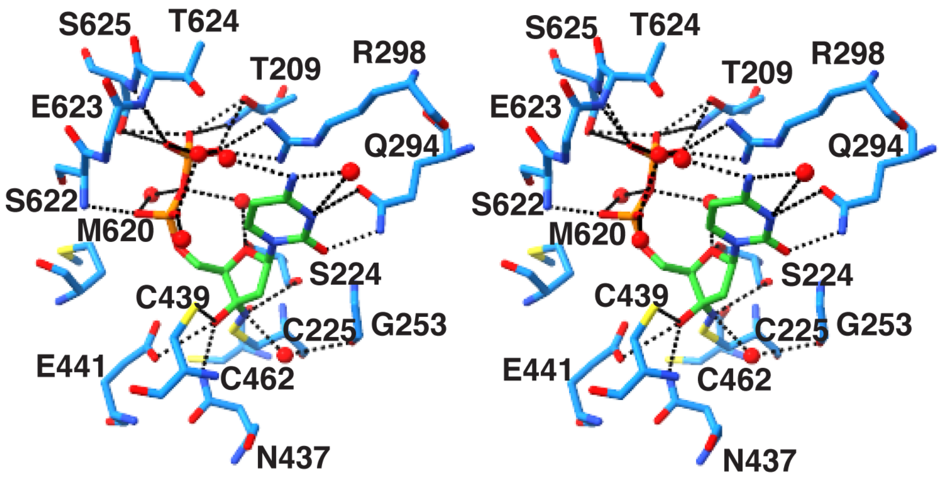


**Figure S9. Stereoview comparisons of CDP binding and of SN-CDP binding. (A)** Interactions between CDP substrate and the active site residues (gray carbons) from PDB 5CNS. Water molecule is shown as a red sphere. Hydrogen bonding interactions are shown with black dashed lines (distance ≤ 3.8 Å). dATP is bound in the specificity site, but is not depicted in the figure. **(B)** Interactions between the SN-CDP nucleotide (green carbons) and the active site residues (dark blue carbons) of $\alpha$ from this work. Water molecules are shown as red spheres. Hydrogen bonding interactions are shown with black dashed lines (distance ≤ 3.8 Å).

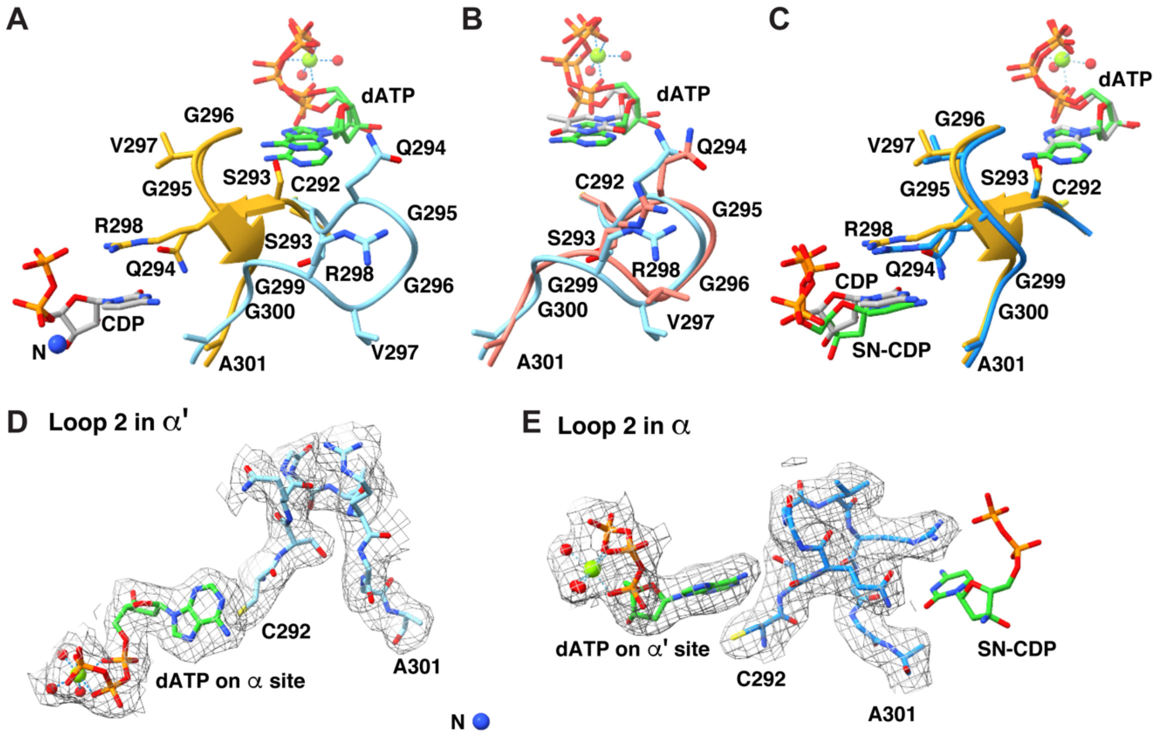


**Figure S10. Loop 2 comparisons. (A)** Superimposition of loop 2 in α′ (light blue, this work) and loop 2 from a previous structure with substrate-effector pair CDP-dATP bound (loop 2 in yellow, CDP with gray carbons and dATP with green carbons, PDB 5CNS). N-adduct (blue sphere) and dATP (green carbons) from current structure are also shown. **(B)** Superimposition of loop 2 in α′ (light blue, this work) and loop 2 from a previous oxidized structure (salmon, PDB 2R1R). dATP molecules are also shown with carbons in green (this work) and in gray (PDB 2R1R). **(C)** Superimposition of loop 2 in α (light blue, this work) and loop 2 from a previous structure with substrate-effector pair CDP-dATP bound (yellow, PDB 5CNS). SN-adduct and dATP (this work) are shown with carbons in green, and CDP and dATP (PDB 5CNS) are shown with carbons in gray. **(D)** Cryo-EM map (grey mesh; sd level 1) for loop 2 (light blue) of α′ and dATP bound to the adjacent specificity site. The dATP molecule is shown with carbons in green, the nitrogen atom is shown as a blue sphere, $Mg^{2+}$ is shown as a green sphere, and water molecules are shown as red spheres. **(E)** Cryo-EM map (grey mesh; sd level 1) for loop 2 of α (dark blue) and dATP bound to the adjacent specificity site.

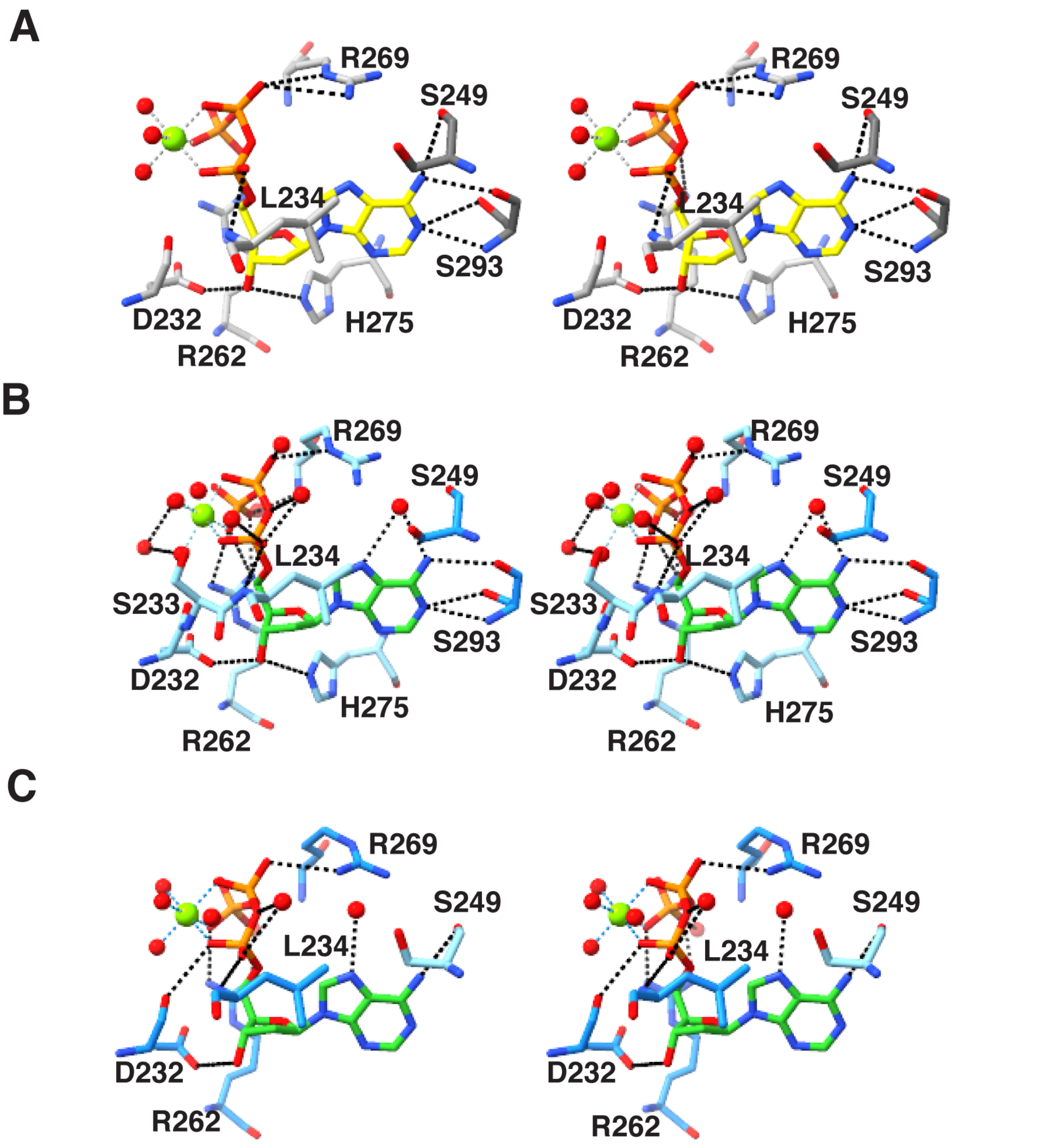


**Figure S11. Stereoview comparison of structures with dATP in the specificity site. (A)** dATP (yellow carbons) bound to specificity site at dimer interface from PDB 5CNS (one subunit in gray and another subunit in dark gray). Substrate CDP was bound in this structure, not shown for clarity. Water molecules are shown as red spheres and $Mg^{2+}$ as a green sphere. Black dashed lines are possible hydrogen bonding interactions with distance ≤ 3.5 Å. **(B)** dATP (green carbons) bound in the specificity site at the dimer interface ($\alpha'$ in light blue, $\alpha$ in dark blue) that communicates with the $\alpha$ active site containing the SN-CDP adduct, which is not shown for clarity. $Mg^{2+}$ is shown as a green sphere and water molecules as red spheres. Black dashed lines are possible hydrogen bonding interactions with distance ≤ 3.5 Å. **(C)** dATP (green carbons) in specificity site at dimer interface ($\alpha'$ in light blue, $\alpha$ in dark blue) that communicates with the $\alpha'$ active site containing the N-adduct, which is not shown for clarity. $Mg^{2+}$ is shown as a green sphere and water molecules as red spheres. Black dashed lines are possible hydrogen bonding interactions with distance ≤ 3.5 Å.

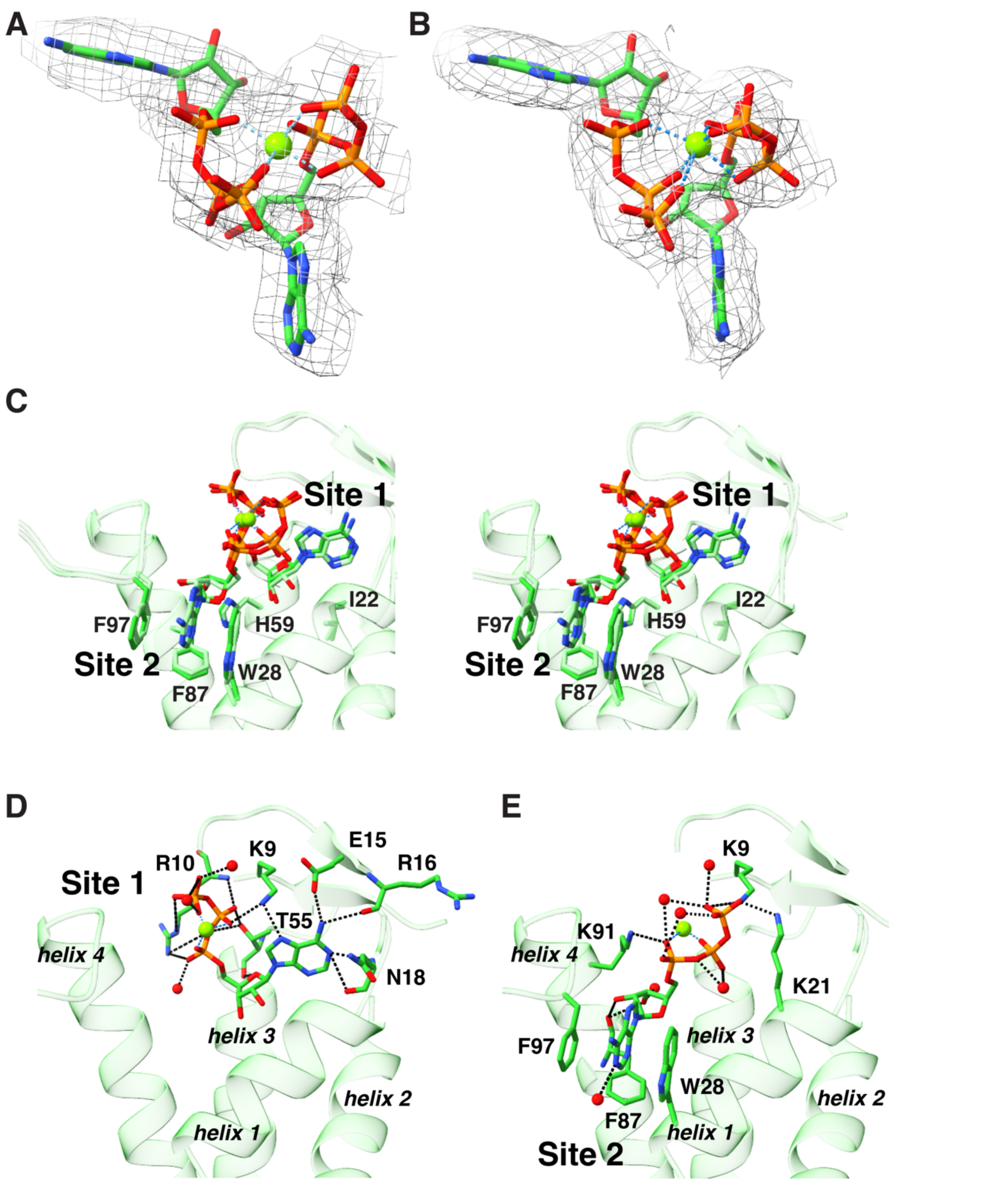


**Figure S12. Two ATP molecules are bound to the activity site in the cone domain in current structure. (A)** Cryo-EM map (grey mesh; sd level 1) for the two ATP molecules (green carbons) bound in the cone domain of $\alpha'$. **(B)** Cryo-EM map (grey mesh; sd level 1) for the two ATP molecules (green) bound in the cone domain of $\alpha$. $Mg^{2+}$ shown as green sphere. **(C)** Stereoview of the two ATP sites in the cone domain of $\alpha$ (green ribbons) with interacting residues indicated. **(D)** ATP molecule (green carbons) in site 1 in the cone domain of $\alpha$ (green ribbons). Dashed black lines indicated potential hydrogen bonding interactions (distance ≤ 3.5 Å). The helices of the cone domain are labeled. **(E)** ATP molecule (green carbons) in site 2 in the cone domain of $\alpha$ (green ribbons). Note: the quality of the cryo-EM map of the cone domain is better for $\alpha$ than for $\alpha'$.

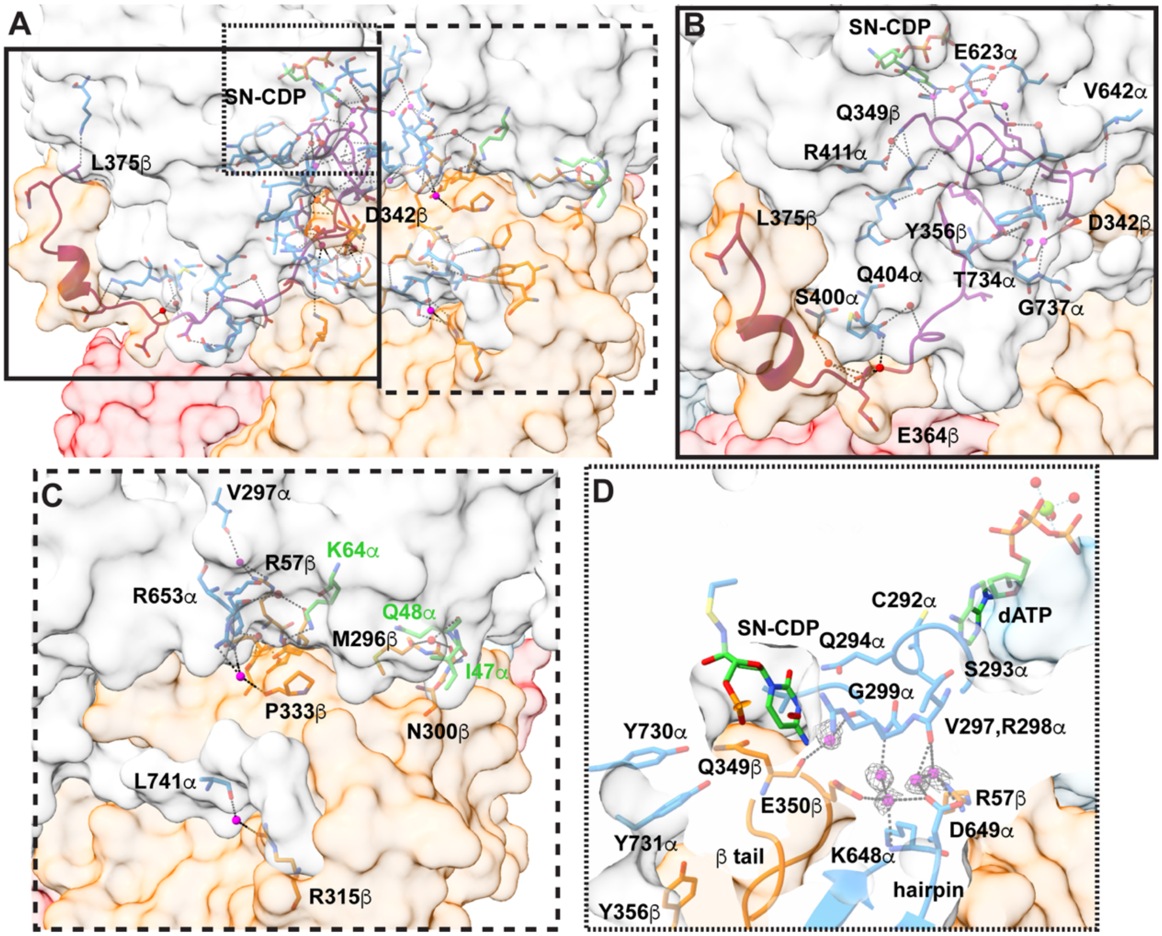


**Figure S13. Bridging water molecules participate in structuring the α–β interface. (A)** Surface view of the α–β interface showing amino acids participating in direct interactions and water-bridged interaction across the interface. ~3600 Å$^2$ of surface area on α (light gray) is buried by β (orange) with an additional ~300 Å$^2$ buried by β′ (red). The main interaction between β and α is made by the β-tail (residues D342 to L375). 13 of the 23 waters found at the α–β interface (**Table S11**) have been visualized previously in crystal structures of α2 or β2. Only ten water molecules are novel to the structure we describe here (shown as magenta spheres). α subunit residues are shown in dark blue, and the β subunit residues are shown in orange except for β-tail residues (D342-L375), which are shown in purple. The boxes indicate the corresponding regions shown in panels **(B)**, **(C),** and **(D)**. **(B)** Zoom-in on the interfacial interactions of the β-tail with α residues either through direct interactions or through a bridging water molecule. **(C)** Zoom in on the α–β interfacial interactions that do not involve the β-tail. Labels for cone domain residues are in green. **(D)** Zoom-in on loop 2 (C292-A301) interactions with SN-CDP, specificity effector dATP (carbons in green), and water molecules (magenta spheres with gray mesh, threshold set to sd level 1). Water molecules mediate the interaction between loop 2 of α, K638 and D649 of the hairpin in α, R57 in β, and the β-tail loop (residues 342-362) that closes off the active site pocket from the aqueous environment. Hydrogen bond interactions are indicated by dashed black lines (distance ≤ 3.5 Å).

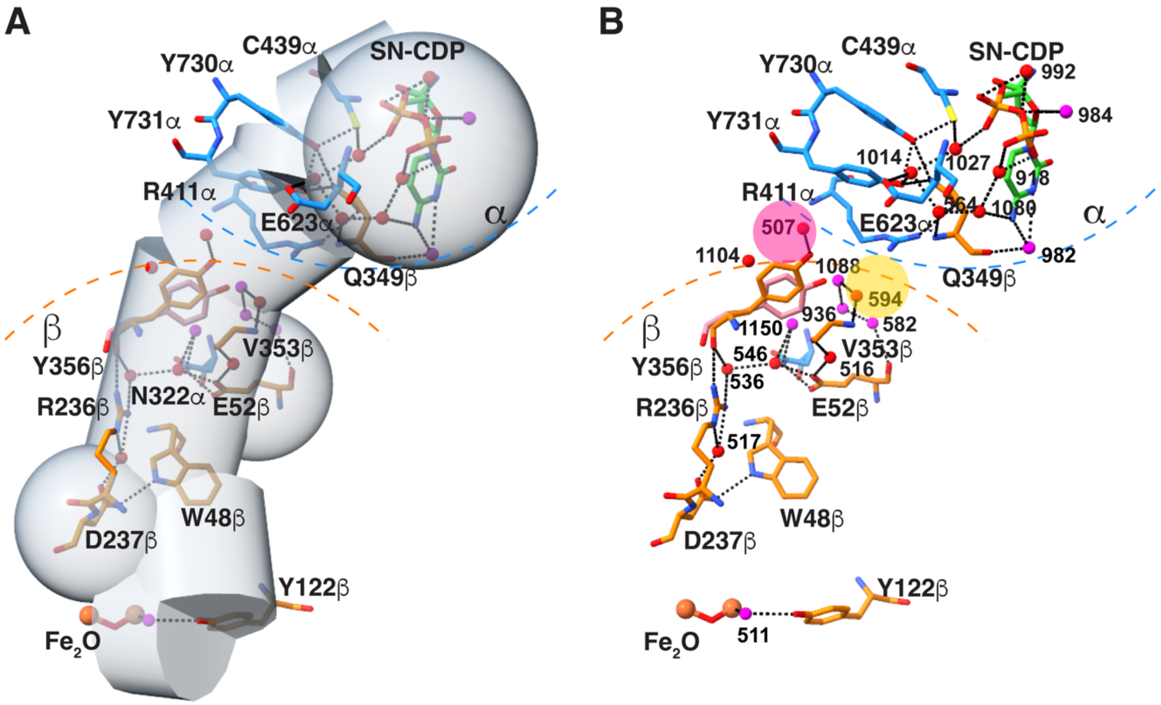


**Figure S14. PCET Waters. (A)** Visualization of "PCET waters," which we defined as those molecules either within a 5-Å radius of $Fe_2O$β, Y122β, D237β, W48β, E52(Q)β, Y356β, Y731α, Y730α, and C439α, and a 7-Å radius of the nucleotide or within 5 Å of the shortest path between electron transfer residue pairs. β residues are shown with orange carbons, Y356β from the previous work (PDB 6W4X) is shown with dark pink carbons, α residues are shown with dark blue carbons, and the SN-CDP adduct is shown with green carbons. The transparent gray surface encompasses the region containing PCET waters, as described in the main text. Water molecules are shown as either red spheres (previously visualized in crystal structures, **Tables S3-S6**) or magenta spheres (newly observed in this work). The black dashed lines indicate potential hydrogen bonds along the radical transfer pathway (distance ≤ 3.8 Å). **(B)** The same image as in **(A)** with water molecules labeled by number. Water molecules numbered 901-1196 correspond to chain B (α) and water molecules numbered 501-608 correspond to chain C (β). Additional hydrogen bonding interactions not depicted to preserve clarity are indicated in **Table S15**. The pink and yellow spheres indicate the direction of the start of two different water channels leading to the surface of the complex and depicted in detail in **Figure 5** and **Figure S16**.

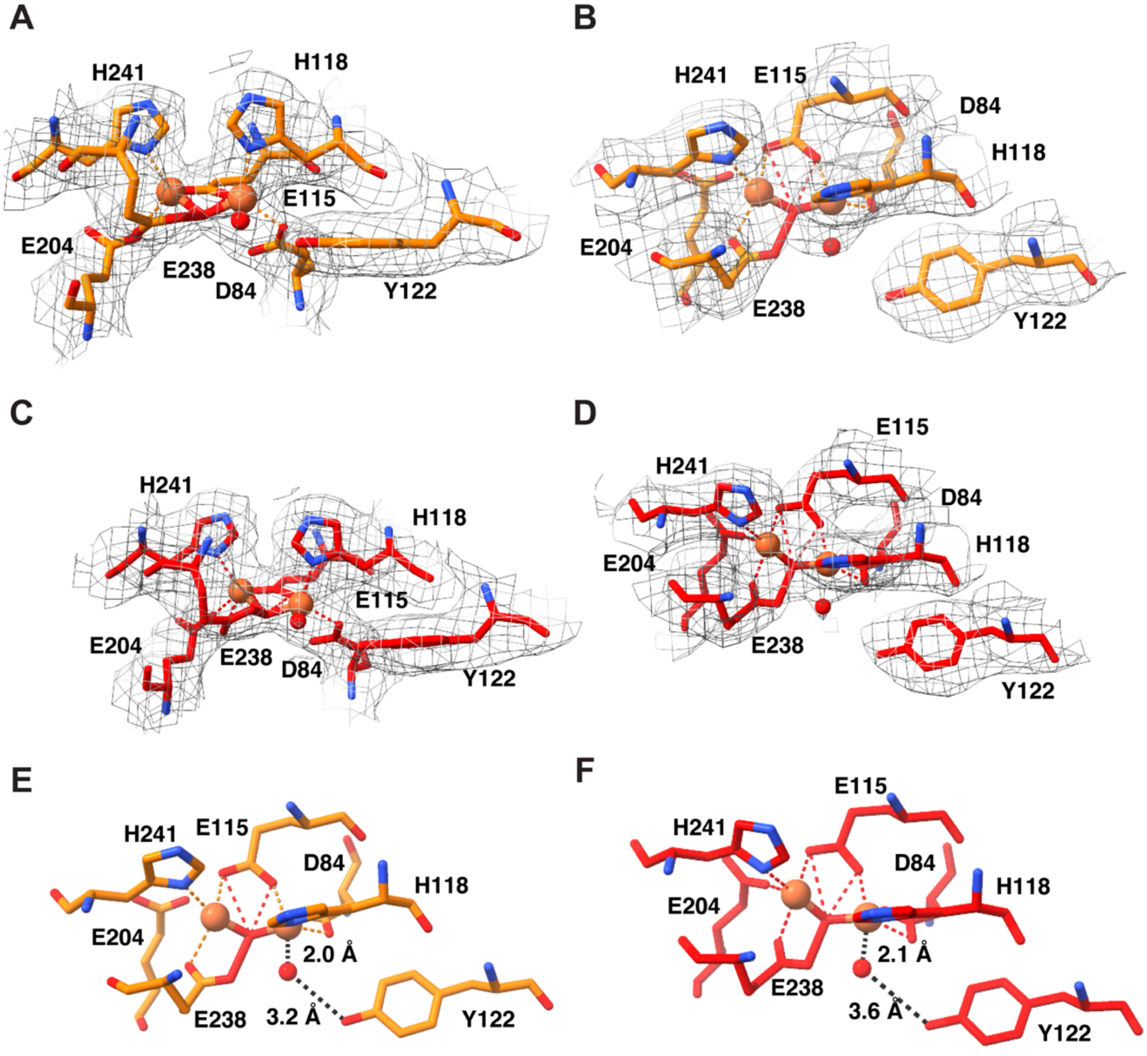


**Figure S15. Cryo-EM density for the di-iron site of β2. (A)** Cryo-EM map (grey mesh; sd level 1) for the di-iron site of β. The iron molecules are shown as orange spheres, the β residues are shown with orange carbons, and the water molecule is shown as a red sphere. **(B)** An orthogonal view of the di-iron site shown in **(A)**. **(C)** Cryo-EM map (grey mesh; sd level 1) for the di-iron site of β′. The iron molecules are shown as orange spheres, the β′ residues are shown with red carbons, and the water molecule is shown as a red sphere. **(D)** An orthogonal view of the di-iron site shown in **(C)**. **(E)** Interactions between the di-iron site of β (orange spheres) with the residues of β (orange carbons) and a single water molecule (red sphere). Potential hydrogen bonding interactions (distance ≤ 3.5 Å) are shown with dashed black lines and the hydrogen bonding distances for the water molecule are shown. The view is the same as in **(B)**. **(F)** Interactions between the di-iron site of β′ (orange spheres) with the residues of β′ (red carbons) and a single water molecule (red sphere). Potential hydrogen bonding interactions (distance ≤ 3.5 Å) are shown with dashed black lines and the hydrogen bonding distances for the water molecule are shown. The view is the same as in **(D)**.

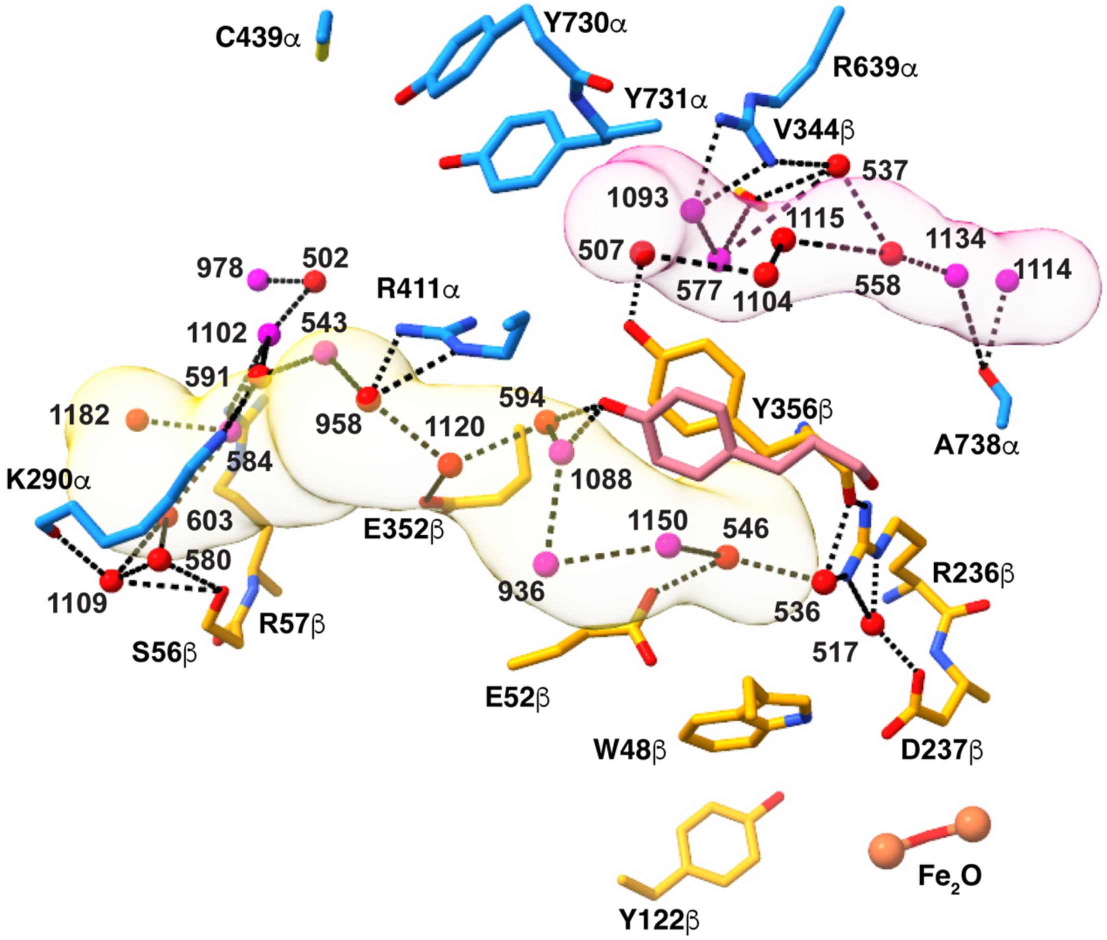


**Figure S16. Water channels and networks in more detail.** Figure is reproduced from **Fig. 5A** of the main text with water numbers added. The 35-Å-long yellow channel begins at Y356β, passes by E52β and E352β and exits at the back of the complex past loop 2 residue K290α. α residues are shown with carbons in dark blue and β residues are shown with carbons in orange. Previously observed water molecules are shown as red spheres and newly observed water molecules are shown as magenta spheres. The 20-Å-long pink channel runs along β-tail residues 342-356 and exits to bulk solvent between the β-tail residue D342β and G737α. Dashed black lines indicate possible hydrogen bonding interactions (distance ≤ 5 Å). Water molecules numbered 901-1196 correspond to chain B (α) and water molecules numbered 501-608 correspond to chain C (β).

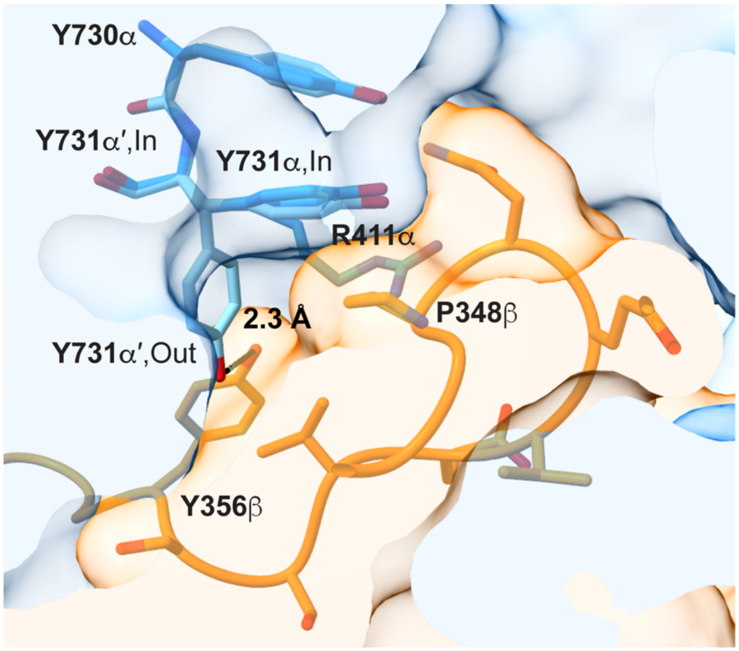


**Figure S17. Surface view of α and β showing the packing of the β-tail (orange) with Y731 and Y730 of α (dark blue).** Y731 and Y730 of α′ (light blue) are overlaid, including both the “flipped in” and “flipped out” conformation of Y731. The packing of P348β and R411α against Y731α appears to restrict the movement of Y731α, which is not “flipped out” when β is bound in either the previous pre-turnover or current mid-turnover structures. If Y731α “flips out” during reverse PCET, its hydroxyl group would be close to the hydroxyl group of Y356β, ~2.3-Å apart. A “flipped out” Y731α is observed post-turnover following β dissociation.

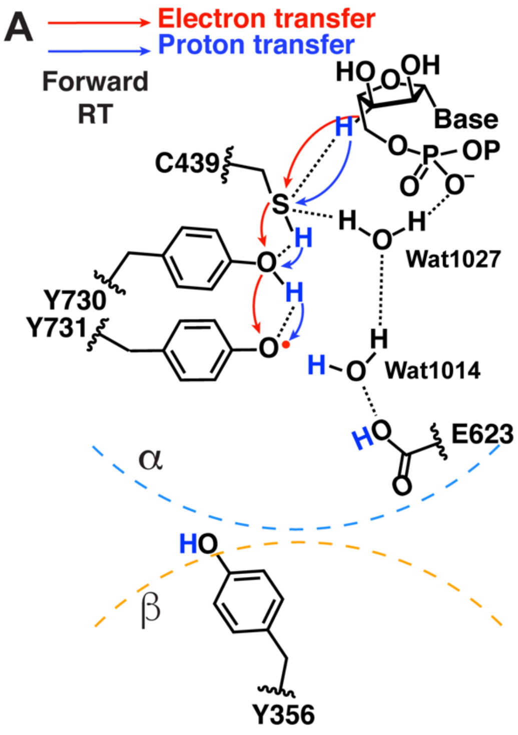


**Figure S18. The water-mediated hydrogen bond network in α could facilitate colinear PCET from Y731α to the substrate. (A)** Depiction of forward RT showing possible deprotonation of Y731α by E623α leading to formation of Y731α•. E623α is shown as flipped out as observed in the mid-turnover structure and water molecule Wat1014 has moved in to take its place. The presence of these water molecules (Wat1014 and Wat1027, both in chain B (α)) and this hydrogen bonding network would appear to restrict the positioning of protons of Y730α and C439α, potentially allowing for facile proton-coupled ET. **(B)** Depiction of reverse RT showing reduction of product radical to form Y356β•. The water-mediated hydrogen bond network within α appears arranged to facilitate rapid reverse radical transfer to Y356β.

**Table S1.** Cryo-EM data collection, reconstruction, and refinement statistics

| **Accession codes** | **PDB: 9DB2; EMDB: EMD-46711; EMPIAR: EMPIAR-12249** |
|---|---|
| **Data collection** | |
| Microscope | FEI Titan Krios |
| Camera | Gatan K2 |
| Acceleration voltage (kV) | 300 |
| Magnification (X) | 105,000 |
| Pixel size (Å) | 0.415 |
| Defocus range ($\mu$m) | -1 to -2.3 |
| Number of frames | 30 |
| Exposure time (s) | 1.2 |
| Total exposure ($e^-$/Å) | 50.946 |
| Shots per hole | 2 |
| Total micrographs collected | 6,007 |
| Particles | |
| Micrographs used for selection | 5,973 |
| Total particles | 1,029,442 |
| **Reconstruction** | |
| Software | RELION-4.0 |
| Final number of refined particles | 440,549 |
| Symmetry imposed | C1 |
| Resolution of unmasked map (Å) | 3.4 |
| Resolution of masked map (Å) | 2.6 |
| FSC threshold | 0.143 |
| Local resolution range (Å) | 2.46-4.45 |
| **Refinement** | |
| Refined model resolution (Å) | 2.6 |
| dFSC model threshold | 0.143 |
| CC (mask)/CC (box) | 0.90/0.66 |
| Model composition | |
| Residues | 2206 |
| Water | 624 |
| SN-CDP adduct from $N_3$CDP | 1 |
| ATP/dATP | 4/2 |
| Fe-O-Fe | 2 |
| $Mg^{2+}$ | 4 |
| Nitrogen resulting from N• | 1 |
| RMSD | |
| Bond lengths (Å) | 0.005 |
| Bond angles (°) | 0.791 |
| Molprobity score | 1.74 |
| Clashscore | 11.84 |
| Ramachandran plot (%) | |
| Favored | 97.13 |
| Allowed | 2.73 |
| Outliers | 0.14 |
| Rotamer Outliers (%) | 0.56 |
| CaBLAM outliers (%) | 1.10 |
| B-factors (min/max/mean) | |

| | |
|---|---|
| Protein | 30.65/184.84/82.82 |
| SN-CDP adduct from $N_3$CDP | 64.60/83.79/73.65 |
| ATP | 78.75/197.01/136.02 |
| dATP | 57.71/74.60/66.22 |
| Fe-O-Fe | 82.93/99.18/92.58 |
| $Mg^{2+}$ | 76.99/166.17/106.50 |
| Nitrogen resulting from N• | 59.93/59.93/59.93 |
| Water | 41.53/142.47/81.68 |
| EMRinger Score | 3.65 |

**Table S2.** The number of residues visualized in the cryo-EM structure of the RNR $\alpha2\beta2$ complex (the $\alpha$ subunit has 761 residues and the $\beta$ subunit has 375 residues).

| Subunit | Chain A ($\alpha'$) | Chain B ($\alpha$) | Chain C ($\beta$) | Chain D ($\beta'$) |
|---|---|---|---|---|
| $\alpha$ | 7–739 | 2–745 | NA | NA |
| $\beta$ | NA | NA | 1–375 | 1–339/360-373 |

**Table S3. List of waters bound to chain A residues ($\alpha$') that have corresponding water molecules in one or more crystal structures.**

| Water Number in Chain A (a')* | Location (relative to Chain A) | 8VHN – Water Number in Chain A* (Chain A aligned by C-alphas) | 2X0X – Water Number in Chain A (Chain A aligned by C-alphas) | 1R1R – Water Number in Chain A (Chain A aligned by C-alphas) | 2XAP – Water Number in Chain A (Chain A aligned by C-alphas) | 2XAV – Water Number in Chain A (Chain A aligned by C-alphas) | 2XAX – Water Number in Chain A (Chain A aligned by C-alphas) | 2XAY – Water Number in Chain A (Chain A aligned by C-alphas) | 2XAZ – Water Number in Chain A (Chain A aligned by C-alphas) | 2XO4 – Water Number in Chain A (Chain A aligned by C-alphas) | 2XO5 – Water Number in Chain A (Chain A aligned by C-alphas) | Qscore |
|---|---|---|---|---|---|---|---|---|---|---|---|---|
| 902 | Chain A internal | 1019 | 2094 | 835 | 2131 | 2075 | 2061 | 2071 | 2027 | 2094 | 2101 | 0.785654 |
| 903 | Chain A surface | N/A | 2234 | N/A | 2320 | 2151 | 2148 | 2184 | 2069 | 2084 | 2215 | 0.693233 |
| 904 | Chain A internal | 992 | 2134 | 780 | 2400 | N/A | N/A | N/A | N/A | 2280 | 2142 | 0.826739 |
| 906 | Chain A surface | 1060 | 2073 | 842 | 2109 | N/A | N/A | 2060 | N/A | 2083 | 2090 | 0.731875 |
| 907 | Chain A surface | 1135 | 2188 | N/A | 2352 | N/A | N/A | N/A | N/A | 2246 | N/A | 0.868261 |
| 908 | Chain A surface | N/A | N/A | N/A | N/A | N/A | 2087 | N/A | N/A | N/A | N/A | 0.664395 |
| 909 | Chain A/B interface | 941 | 2116 | N/A | 2157 | 2080 | N/A | 2094 | 2037 | 2112 | 2121 | 0.882817 |
| 910 | Chain A surface | N/A | N/A | N/A | N/A | N/A | 2168 | N/A | N/A | N/A | N/A | 0.804106 |
| 911 | Chain A surface | N/A | N/A | N/A | N/A | N/A | N/A | 2178 | N/A | N/A | N/A | 0.850262 |
| 912 | Chain A surface | 1047 | 2033 | N/A | 2048 | N/A | N/A | N/A | N/A | N/A | 2051 | 0.72025 |
| 914 | Chain A internal | N/A | 2117 | N/A | 2159 | N/A | N/A | N/A | N/A | N/A | N/A | 0.792557 |
| 915 | Chain A/B interface near dATP site in Chain A | 1073 | N/A | N/A | 2148 in Chain B | N/A | N/A | N/A | N/A | N/A | N/A | 0.64521 |
| 916 | Chain A surface | 963 | 2260 | N/A | 2362 | N/A | N/A | 2213 | N/A | 2252 | 2256 | 0.743428 |
| 917 | Chain A internal | 920 | 2108 | 762 | 2178 | 2098 | 2066 | 2119 | 2045 | 2139 | 2110 | 0.882335 |
| 918 | Chain A surface | N/A | N/A | N/A | 2265 | 2131 | N/A | 2157 | N/A | N/A | N/A | 0.681624 |
| 919 | Chain A surface near dATP site in Chain A | 1057 | N/A | N/A | N/A | N/A | N/A | N/A | N/A | N/A | N/A | 0.854175 |
| 920 | Chain A surface | 1022 | 2143 | N/A | 2195 | N/A | 2100 | N/A | N/A | N/A | 2151 | 0.646932 |
| 921 | Chain A surface | 1044 | 2147 | N/A | 2197 | 2101 | N/A | N/A | N/A | N/A | N/A | 0.75953 |
| 922 | Chain A surface | 1173 | 2020 | N/A | 2035 | 2026 | 2025 | 2027 | 2009 | 2033 | 2038 | 0.592626 |
| 923 | Chain A surface | N/A | 2228 | N/A | 2307 | N/A | 2004 | 2002 | 2002 | 2003 | 2005 | 0.785767 |
| 924 | Chain A internal | 1006 | 2257 | 862 | 2358 | N/A | N/A | N/A | N/A | N/A | N/A | 0.895725 |
| 925 | Chain A internal | 1027 | 2139 | N/A | 2189 | N/A | N/A | N/A | N/A | N/A | N/A | 0.778061 |
| 927 | Chain A surface | 959 | 2138 | N/A | 2188 | N/A | N/A | N/A | N/A | N/A | 2148 | 0.808472 |

| | | | | | | | | | | | | |
|---|---|---|---|---|---|---|---|---|---|---|---|---|
| 928 | Chain A internal | 1039 | 2160 | 776 | 2217 | 2174 | 2111 | 2133 | 2050 | 2158 | 2167 | 0.856547 |
| 929 | Chain A surface | 1021 | 2162 | 806 | 2220 | N/A | 2110 | 2135 | 2052 | 2159 | 2168 | 0.670943 |
| 930 | Chain A surface | N/A | 2201 | 819 | 2270 | 2134 | 2133 | 2162 | 2061 | 2196 | 2193 | 0.707091 |
| 932 | Chain A surface | 1079 | 2294 | 767 | 2390 | N/A | N/A | N/A | N/A | 2249 | N/A | 0.828024 |
| 933 | Chain A internal | 999 | 2044 | 769 | 2079 | 2050 | 2042 | 2047 | 2018 | 2061 | 2066 | 0.661522 |
| 934 | Chain A surface | N/A | 2197 | 860 | 2266 | N/A | 2128 | 2156 | N/A | 2190 | 2189 | 0.806088 |
| 935 | Chain A surface | N/A | N/A | N/A | N/A | N/A | 2079 | N/A | N/A | N/A | N/A | 0.620163 |
| 936 | Chain A surface | N/A | 2064 | N/A | 2095 | N/A | N/A | N/A | N/A | N/A | N/A | 0.512302 |
| 937 | Chain A surface | 1059 | 2166 | N/A | 2224 | 2113 | 2114 | N/A | 2053 | N/A | 2171 | 0.845694 |
| 938 | Chain A/B interface | 979 | 2100 | 800 | 2144 | 2069 | N/A | 2077 | 2031 | 2097 | 2105 | 0.775077 |
| 939 | Chain A surface | 911 | N/A | N/A | N/A | 2110 | N/A | 2131 | N/A | N/A | 2164 | 0.825768 |
| 940 | Chain A surface | 1005 | 2152 | N/A | 2202 | 2102 | 2091 | 2122 | N/A | N/A | N/A | 0.863964 |
| 941 | Chain A internal | 1002 | 2179 | 771 | 2234 | 2132 | 2131 | 2159 | 2059 | 2192 | 2190 | 0.82746 |
| 942 | Chain A surface | N/A | 2232 | N/A | 2316 | N/A | N/A | 2181 | N/A | 2224 | N/A | 0.844911 |
| 943 | Chain A internal | 1037 | 2088 | 827 | 2127 | N/A | 2060 | 2067 | 2024 | 2090 | N/A | 0.787331 |
| 945 | Y730/731 | 944 | 2132 | 845 | 2177 | N/A | N/A | N/A | N/A | 2138 | N/A | 0.89132 |
| 946 | Chain A surface | 1053 | N/A | N/A | 2357 | N/A | N/A | N/A | N/A | N/A | N/A | 0.808946 |
| 947 | Chain A surface | 1076 | 2032 | N/A | 2044 | N/A | N/A | N/A | N/A | N/A | N/A | 0.464073 |
| 948 | Chain A surface | 951 | 2161 | 809 | 2223 | 2112 | 2113 | 2134 | 2051 | 2163 | 2170 | 0.832905 |
| 949 | Chain A surface | N/A | 2059 | N/A | 2087 | N/A | 2048 | 2051 | N/A | N/A | 2082 | 0.841729 |
| 950 | Chain A surface | 1131 | N/A | N/A | N/A | N/A | 2032 | N/A | N/A | N/A | N/A | 0.563393 |
| 952 | Chain A internal | 1029 | 2296 | 781 | 2394 | N/A | 2180 | 2223 | N/A | 2275 | 2271 | 0.836936 |
| 953 | Chain A surface | 1101 | 2217 | N/A | 2292 | 2141 | 2139 | N/A | N/A | 2210 | N/A | 0.590693 |
| 955 | ATP site | 964 | N/A | N/A | N/A | N/A | N/A | N/A | N/A | N/A | N/A | 0.901812 |
| 956 | Chain A internal | N/A | 2126 | N/A | 2169 | N/A | N/A | N/A | N/A | 2123 | N/A | 0.766351 |
| 957 | Chain A surface | 973 | 2123 | N/A | 2165 | 2089 | N/A | 2104 | N/A | 2124 | 2133 | 0.887337 |
| 959 | Chain A internal | 1000 | 2170 | 773 | 2229 | 2116 | N/A | 2143 | N/A | 2168 | 2175 | 0.914041 |
| 960 | Chain A internal | 1034 | 2250 | 829 | 2345 | N/A | N/A | 2175 | 2068 | 2240 | 2232 | 0.784187 |
| 961 | Chain A surface | 966 | 2283 | 843 | 2398 | N/A | N/A | N/A | N/A | N/A | 2260 | 0.715516 |
| 962 | Chain A/C interface | 1043 | 2125 | 804 | 2167 | N/A | 2082 | 2103 | 2041 | 2126 | 2134 | 0.852138 |
| 963 | dATP site in Chain C and A/B interface | 1117 | N/A | 836 | 2134 | N/A | N/A | 2073 | 2029 | N/A | N/A | 0.857832 |

| | | | | | | | | | | | | |
|---|---|---|---|---|---|---|---|---|---|---|---|---|
| 964 | Chain A surface | 1133 | 2236 | N/A | 2322 | N/A | 2150 | N/A | N/A | 2227 | 2266 | 0.857072 |
| 966 | Chain A surface | 955 | N/A | N/A | 2153 | N/A | 2108 | N/A | N/A | N/A | N/A | 0.8266 |
| 967 | Chain A/C interface | 1024 | 2129 | N/A | 2172 | N/A | 2090 | 2111 | N/A | 2127 | 2138 | 0.827116 |
| 968 | Chain A internal | 954 | 2093 | 772 | 2130 | 2065 | N/A | 2070 | 2026 | 2093 | 2100 | 0.897656 |
| 969 | Chain A surface, near Chain B | N/A | N/A | N/A | 2120 | N/A | N/A | N/A | N/A | N/A | N/A | 0.69598 |
| 970 | Chain A internal | 1144 | N/A | N/A | N/A | N/A | N/A | N/A | N/A | N/A | N/A | 0.768177 |
| 971 | Chain A surface | 1182 | N/A | N/A | N/A | N/A | N/A | N/A | N/A | N/A | N/A | 0.752754 |
| 972 | Chain A/C interface | 1017 | 2124 | 813 | 2166 | N/A | 2084 | 2105 | 2042 | 2125 | N/A | 0.800749 |
| 973 | Chain A internal | 1009 | 2169 | 777 | 2228 | 2115 | N/A | 2219 | 2082 | 2167 | 2267 | 0.870093 |
| 975 | Chain A surface, near dATP site in Chain A | 1158 | N/A | N/A | N/A | N/A | N/A | N/A | N/A | N/A | N/A | 0.707536 |
| 976 | Chain A internal | 967 | 2120 | N/A | 2160 | 2082 | 2075 | 2096 | 2038 | 2115 | 2125 | 0.90376 |
| 977 | Chain A/B interface | N/A | 2122 | N/A | 2162 | N/A | N/A | 2106 in Chain B | N/A | 2142 in Chain B | 2127 | 0.818447 |
| 978 | Chain A internal | 987 | 2300 | 803 | 2393 | N/A | 2183 | 2220 | 2083 | 2155 | 2268 | 0.850388 |
| 979 | Chain A/B interface | 949 | 2099 | 807 | 2138 | 2068 | 2064 | 2076 | 2030 | 2096 | 2104 | 0.937777 |
| 980 | dATP site in Chain A | 1192 | N/A | N/A | N/A | N/A | N/A | N/A | N/A | N/A | N/A | 0.63099 |
| 981 | Chain A surface | 1157 | 2243 | 854 | 2331 | 2145 | 2143 | 2189 | N/A | 2213 | 2222 | 0.840002 |
| 983 | Chain A surface near ATP site | 1120 | 2030 | N/A | 2041 | N/A | N/A | 2033 | 2012 | 2042 | 2044 | 0.382403 |
| 985 | Chain A surface | N/A | 2263 | 857 | 2363 | N/A | N/A | N/A | N/A | N/A | N/A | 0.813933 |
| 986 | Chain A surface | N/A | 2068 | 850 | 2100 | N/A | 2053 | N/A | N/A | 2080 | N/A | 0.815836 |
| 988 | Chain A surface | 1054 | 2141 | N/A | 2192 | N/A | 2097 | N/A | N/A | 2143 | N/A | 0.916273 |
| 989 | Chain A/B interface | 1072 in Chain B | 2121 in Chain B | 1195 in Chain B | 2158 | N/A | 2059 in Chain B | 2090 in Chain B | 2036 in Chain B | 2123 in Chain B | 2122 | 0.852755 |
| 990 | Chain A surface | N/A | 2070 | N/A | 2308 | 2149 | N/A | N/A | N/A | N/A | 2210 | 0.892792 |
| 991 | Chain A internal | 1196 | N/A | N/A | N/A | N/A | N/A | N/A | N/A | N/A | N/A | 0.842984 |
| 992 | Chain A internal | 1004 | 2289 | 778 | 2387 | N/A | 2182 | 2224 | N/A | 2271 | 2265 | 0.894157 |
| 993 | Chain A surface | N/A | N/A | N/A | 2036 | N/A | N/A | N/A | N/A | N/A | N/A | 0.786828 |
| 994 | Chain A surface | 989 | 2103 | N/A | N/A | N/A | N/A | N/A | N/A | N/A | N/A | 0.837289 |
| 996 | Chain A internal | 1062 | 2231 | 821 | 2317 | N/A | N/A | N/A | N/A | 2223 | N/A | 0.857796 |
| 997 | Chain A/B interface | N/A | 2110 in Chain B | N/A | 2146 in Chain B | N/A | N/A | N/A | N/A | N/A | N/A | 0.791944 |

| | | | | | | | | | | | | |
|---|---|---|---|---|---|---|---|---|---|---|---|---|
| | near dATP in Chain A | | | | | | | | | | | |
| 998 | Chain A/B interface | 1093 | 2098 | 832 | 2136 | N/A | N/A | 2074 | N/A | N/A | N/A | 0.823458 |
| 999 | Chain A surface | N/A | N/A | N/A | N/A | N/A | N/A | N/A | N/A | N/A | 2161 | 0.643422 |
| 1000 | Chain A surface | N/A | N/A | N/A | N/A | N/A | N/A | 2118 | N/A | N/A | 2145 | 0.767028 |
| 1001 | Chain A surface | 1095 | N/A | N/A | N/A | N/A | 2105 | 2125 | N/A | 2151 | 2158 | 0.715045 |
| 1002 | Chain A surface | N/A | 2235 | N/A | 2321 | 2152 | 2149 | 2185 | 2070 | 2226 | 2216 | 0.813629 |
| 1003 | Chain A surface near dATP in Chain A | 1052 | N/A | N/A | N/A | N/A | N/A | N/A | N/A | N/A | N/A | 0.862214 |
| 1004 | Chain A/B interface | N/A | N/A | N/A | N/A | N/A | N/A | N/A | N/A | N/A | 2120 | 0.821012 |
| 1005 | Chain A/B interface | N/A | N/A | N/A | N/A | N/A | 2073 | N/A | N/A | N/A | N/A | 0.687051 |
| 1008 | Chain A surface | 1068 | N/A | N/A | 2236 | N/A | N/A | N/A | N/A | N/A | N/A | 0.608874 |
| 1009 | Chain A surface | N/A | 2199 | N/A | 2268 | N/A | N/A | N/A | N/A | N/A | N/A | 0.822701 |
| 1010 | Chain A surface | 1066 | N/A | N/A | N/A | N/A | N/A | N/A | N/A | N/A | N/A | 0.840904 |
| 1011 | Chain A surface | N/A | 2103 | N/A | 2163 | N/A | 2081 | 2102 | 2040 | 2099 | 2132 | 0.776025 |
| 1012 | Chain A surface | 1137 | N/A | N/A | N/A | N/A | N/A | N/A | N/A | N/A | N/A | 0.484934 |
| 1013 | Chain A/B interface | 1074 | 2096 | N/A | 2135 | N/A | N/A | N/A | N/A | N/A | N/A | 0.826983 |
| 1016 | Chain A surface | 1138 | 2095 | 844 | 2132 | 2066 | 2062 | 2087 | N/A | 2095 | 2102 | 0.847849 |
| 1017 | Chain A surface | N/A | 2151 | 826 | 2203 | N/A | N/A | N/A | N/A | 2147 | 2153 | 0.772199 |
| 1018 | Chain A surface | 1156 | 2245 | N/A | 2333 | N/A | N/A | 2190 | 2072 | 2233 | 2223 | 0.690652 |
| 1020 | Chain A surface | N/A | 2284 | N/A | 2384 | N/A | N/A | 2216 | N/A | N/A | 2261 | 0.547331 |
| 1021 | Chain A surface | 1236 | N/A | N/A | 2276 | N/A | N/A | N/A | N/A | N/A | N/A | 0.712903 |
| 1022 | Chain A surface | 1167 | 2119 | 763 | 2174 | 2081 | N/A | 2113 | 2044 | 2134 | 2124 | 0.843816 |
| 1023 | Chain A surface | 1028 | 2106 | N/A | N/A | 2088 | N/A | 2083 | 2033 | 2101 | N/A | 0.893164 |
| 1024 | Chain A surface | N/A | 2240 | N/A | 2328 | N/A | 2154 | 2188 | 2071 | 2229 | 2219 | 0.633664 |
| 1025 | Chain A surface | 1110 | N/A | N/A | N/A | 2043 | N/A | 2042 | N/A | N/A | N/A | 0.795674 |
| 1026 | Chain A surface | N/A | 2253 | N/A | N/A | N/A | N/A | 2196 | N/A | 2245 | N/A | 0.836521 |
| 1027 | Chain A surface | N/A | 2154 | N/A | 2205 | N/A | N/A | N/A | N/A | N/A | 2154 | 0.832856 |
| 1028 | Chain A/C interface | N/A | N/A | N/A | 2171 | N/A | N/A | 2110 | N/A | 2132 | 2137 | 0.817144 |
| 1029 | Chain A surface | N/A | 2304 | N/A | 2408 | N/A | N/A | N/A | N/A | 2287 | N/A | 0.78381 |
| 1030 | Chain A surface | 1121 | 2255 | 766 | 2355 | 2160 | 2162 | 2198 | N/A | 2247 | 2241 | 0.810324 |
| 1031 | Chain A internal | 1075 | 2233 | 852 | 2319 | N/A | 2181 | 2183 | N/A | 2225 | 2214 | 0.825019 |

| | | | | | | | | | | | | |
|---|---|---|---|---|---|---|---|---|---|---|---|---|
| 1032 | Chain A surface | N/A | N/A | N/A | 2084 | N/A | N/A | N/A | N/A | 2189 | N/A | 0.773776 |
| 1035 | Chain A surface | N/A | 2214 | 802 | 2287 | 2140 | 2138 | 2169 | 2064 | 2207 | 2200 | 0.919886 |
| 1036 | Chain A/B interface | 1123 | 2180 | N/A | 2235 | N/A | N/A | N/A | N/A | N/A | N/A | 0.857156 |
| 1037 | Chain A surface | 1116 | 2145 | N/A | 2193 | 2099 | N/A | 2120 | 2047 | 2145 | 2152 | 0.835655 |
| 1038 | Chain A surface | 995 | 2055 | 846 | 2085 | N/A | N/A | 2049 | N/A | 2066 | N/A | 0.892225 |
| 1039 | Chain A surface | N/A | 2136 | N/A | 2180 | N/A | 2092 | 2116 | 2046 | 2140 | 2143 | 0.854425 |
| 1041 | Chain A surface | 1210 | 2077 | N/A | 2116 | 2059 | N/A | N/A | N/A | N/A | 2094 | 0.79807 |
| 1042 | Chain A internal | 1144 | 2203 | 799 | 2272 | 2125 | 2134 | 2149 | N/A | 2179 | 2176 | 0.844352 |
| 1045 | Chain A internal | 1007 | 2076 | 811 | 2107 | 2058 | 2054 | N/A | N/A | 2082 | N/A | 0.890018 |
| 1046 | Chain A surface near Chain C | 905 | N/A | N/A | 2168 | N/A | 2083 | N/A | N/A | N/A | N/A | 0.831224 |
| 1047 | Chain A surface | N/A | 2146 | N/A | 2196 | N/A | N/A | 2121 | 2048 | 2146 | N/A | 0.886278 |
| 1049 | Chain A/B interface | N/A | 2039 in Chain B | N/A | 2054 in Chain B | N/A | N/A | 2035 in Chain B | N/A | 2119 | N/A | 0.852794 |
| 1050 | Chain A surface | N/A | 2038 | N/A | 2057 | N/A | N/A | N/A | N/A | N/A | N/A | 0.789683 |
| 1008 in Chain B | Chain A/B interface | 1014 | 2118 | 807 | 2137 | N/A | 2074 | 2075 | N/A | 2114 | 2123 | 0.863672 |
| 1169 in Chain B | Chain A/B interface | 1138 in Chain B | N/A | N/A | 2238 | N/A | N/A | N/A | N/A | N/A | N/A | 0.830731 |
| 526 in Chain C | Chain A/C interface | 1223 | N/A | N/A | N/A | N/A | N/A | N/A | N/A | N/A | N/A | 0.845592 |
| 608 in Chain C | Chain A/C interface | 1226 | N/A | N/A | N/A | N/A | N/A | N/A | N/A | N/A | N/A | 0.89588 |
| 550 in Chain D | Chain A/D interface | N/A | 2158 | N/A | N/A | N/A | N/A | N/A | N/A | 2001 | N/A | 0.753131 |

*unless the Chain identity is otherwise specified

**Table S4. List of waters bound to chain B residues (α) that have corresponding water molecules in one or more crystal structures.**

| Water Number in Chain B (α)* | Location (relative to Chain B) | 8VHN – Water Number in Chain B (Chain B aligned by C-alphas) | 2X0X – Water Number in Chain B (Chain B aligned by C-alphas) | 1R1R – Water Number in Chain B (Chain B aligned by C-alphas) | 2XAP – Water Number in Chain B (Chain B aligned by C-alphas) | 2XAV – Water Number in Chain B (Chain B aligned by C-alphas) | 2XAX – Water Number in Chain B (Chain B aligned by C-alphas) | 2XAY – Water Number in Chain B (Chain B aligned by C-alphas) | 2XAZ – Water Number in Chain B (Chain B aligned by C-alphas) | 2XO4 – Water Number in Chain B (Chain B aligned by C-alphas) | 2XO5 – Water Number in Chain B (Chain B aligned by C-alphas) | 5CNS - Water Number in Chain B (Chain B aligned by C-alphas) | Qscore |
|---|---|---|---|---|---|---|---|---|---|---|---|---|---|
| 901 | Chain B internal | 950 | 2197 | 863 | 2260 | 2060 | N/A | N/A | N/A | 2087 | 2076 | N/A | 0.738701 |
| 902 | Chain B internal | 1142 | 2251 | N/A | N/A | N/A | N/A | N/A | N/A | N/A | N/A | N/A | 0.89395 |
| 903 | Chain B internal | 982 | 2173 | 791 | 2234 | 2067 | 2049 | 2144 | 2049 | 2106 | 2085 | 902 | 0.761051 |
| 904 | Chain B surface near dATP site in Chain B | 1046 | 2150 | 2194 | N/A | N/A | N/A | N/A | N/A | 2153 | 2130 | N/A | 0.816604 |
| 906 | Chain B internal | 968 | 2114 | 762 | 2151 | 2068 | 2071 | 2080 | N/A | 2146 | 2086 | N/A | 0.839548 |
| 907 | Chain B internal | 1078 | 2208 | 765 | 2267 | 2151 | 2115 | 2172 | N/A | 2206 | N/A | N/A | 0.83544 |
| 908 | Chain B internal | 961 | 2256 | N/A | 2335 | N/A | 2164 | N/A | N/A | 2287 | 2243 | N/A | 0.812621 |
| 909 | Chain B surface | 1200 | 2155 | N/A | 2200 | 2101 | 2085 | 2124 | N/A | 2157 | 2133 | N/A | 0.848451 |
| 910 | Chain B internal | 1066 | 2160 | N/A | 2213 | N/A | N/A | 2131 | N/A | N/A | N/A | N/A | 0.787544 |
| 911 | Chain B surface | 1074 | 2048 | 822 | 2070 | N/A | N/A | N/A | N/A | N/A | N/A | N/A | 0.761132 |
| 912 | Chain B internal | 1121 | 2235 | 838 | 2304 | N/A | N/A | N/A | N/A | N/A | 2203 | N/A | 0.859426 |
| 913 | Chain B internal | 1061 | N/A | N/A | N/A | N/A | N/A | N/A | N/A | N/A | N/A | N/A | 0.904837 |
| 914 | Chain B internal | 903 | 2089 | N/A | 2115 | N/A | N/A | N/A | N/A | 2083 | 2212 | N/A | 0.695238 |
| 916 | Chain B surface | 993 | 2161 | N/A | N/A | N/A | N/A | N/A | N/A | N/A | N/A | N/A | 0.69012 |
| 917 | Chain B internal | 1043 | 2223 | 782 | 2315 | N/A | N/A | 2186 | 2071 | 2224 | 2196 | 904 | 0.623789 |
| 918 | Chain B internal, in active site of Chain B | 1116 | N/A | N/A | 2329 | N/A | N/A | N/A | N/A | 2248 | N/A | N/A | 0.491714 |
| 919 | Chain B internal | 959 | 2252 | 786 | 2336 | 2155 | 2143 | 2210 | 2083 | 2251 | 2214 | N/A | 0.85819 |
| 920 | Chain A/B interface, near dATP site in Chain B | 1025 | N/A | N/A | N/A | N/A | N/A | N/A | N/A | N/A | N/A | N/A | 0.659248 |
| 921 | Chain B surface near Chain C | 951 | 2261 | N/A | 2341 | 2169 | N/A | 2214 | N/A | 2255 | 2235 | N/A | 0.896796 |

| | | | | | | | | | | | | |
|---|---|---|---|---|---|---|---|---|---|---|---|---|
| 922 | Chain B internal | 952 | 2064 | 770 | 2091 | 2039 | 2033 | 2046 | 2018 | 2064 | 2060 | 903 | 0.828868 |
| 923 | Chain B internal near Chain C | 931 | 2257 | 862 | 2334 | N/A | N/A | N/A | N/A | 2252 | N/A | N/A | 0.853353 |
| 925 | Chain B surface | 942 | 2297 | 789 | 2380 | 2177 | 2167 | N/A | N/A | 2293 | 2249 | 901 | 0.847138 |
| 927 | Chain B/C interface | 1088 | 2255 | N/A | 2337 | N/A | 2144 | 2211 | N/A | N/A | N/A | N/A | 0.87707 |
| 928 | Chain B surface | 1232 | N/A | N/A | 2061 | N/A | N/A | N/A | N/A | N/A | N/A | N/A | 0.783364 |
| 929 | Chain B internal | 924 | 2298 | 852 | 2372 | 2178 | 2168 | 2237 | 2095 | 2295 | 2250 | N/A | 0.892962 |
| 931 | Chain B internal | 1023 | 2068 | 846 | 2096 | N/A | 2035 | N/A | N/A | N/A | N/A | N/A | 0.849053 |
| 933 | Chain B internal | 1093 | 2144 | 780 | 2185 | 2097 | 2073 | 2113 | 2042 | 2148 | 2126 | N/A | 0.841757 |
| 934 | Chain B surface | 1075 | N/A | N/A | 2197 | 2099 | N/A | N/A | 2044 | N/A | N/A | N/A | 0.875564 |
| 935 | Chain B surface | 1103 | 2214 | 784 | 2277 | 2134 | 2123 | 2180 | 2065 | 2212 | 2185 | N/A | 0.737449 |
| 937 | Chain B internal | 1031 | 2290 | 803 | 2389 | 2108 | N/A | N/A | N/A | 2300 | 2242 | N/A | 0.802152 |
| 939 | Chain B internal | 1026 | 2301 | 779 | 2387 | N/A | N/A | 2241 | 2097 | 2284 | 2241 | N/A | 0.796228 |
| 940 | Chain B surface | N/A | N/A | N/A | 2262 | 2128 | N/A | N/A | N/A | N/A | N/A | N/A | 0.801413 |
| 941 | Chain B surface | 1047 | 2230 | 797 | 2301 | N/A | N/A | N/A | N/A | 2231 | N/A | N/A | 0.846497 |
| 942 | Chain B surface | 1118 | N/A | N/A | N/A | N/A | N/A | N/A | N/A | N/A | N/A | N/A | 0.86814 |
| 943 | Chain B surface | 1147 | 2273 | N/A | 2355 | N/A | 2154 | 2223 | N/A | N/A | 2229 | N/A | 0.796431 |
| 944 | Chain B surface | 1272 | N/A | N/A | N/A | N/A | 2004 | 2007 | N/A | N/A | N/A | N/A | 0.688242 |
| 945 | Chain B surface | 1278 | N/A | N/A | N/A | N/A | N/A | N/A | N/A | N/A | N/A | N/A | 0.74647 |
| 946 | Chain B internal | 943 | 2053 | 769 | 2092 | 2046 | N/A | 2053 | 2019 | 2065 | 2053 | N/A | 0.832047 |
| 947 | Chain B surface | 1192 | N/A | N/A | N/A | N/A | N/A | N/A | N/A | N/A | N/A | N/A | 0.617738 |
| 949 | Chain B surface | 1157 | 2178 | N/A | 2302 | N/A | 2127 | N/A | N/A | N/A | N/A | N/A | 0.582454 |
| 950 | Chain B surface | N/A | 2243 | N/A | N/A | N/A | 2135 | 2201 | N/A | 2242 | N/A | N/A | 0.661157 |
| 951 | Chain B/C interface | N/A | 2267 | N/A | 2079 | N/A | N/A | 2047 | N/A | N/A | 2225 | N/A | 0.714153 |
| 952 | Chain B surface near Chain C | 1003 | N/A | N/A | N/A | N/A | N/A | N/A | N/A | N/A | N/A | N/A | 0.73239 |
| 953 | Chain B internal | 902 | N/A | N/A | N/A | N/A | N/A | N/A | N/A | N/A | N/A | N/A | 0.854074 |
| 954 | Chain B internal, in active site of Chain B | 912 | N/A | N/A | N/A | N/A | N/A | N/A | N/A | N/A | N/A | N/A | 0.826124 |
| 957 | Chain B/C interface | 989 | N/A | N/A | 2221 | N/A | N/A | N/A | N/A | N/A | N/A | N/A | 0.853547 |
| 958 | Chain B/C interface | 1099 | N/A | N/A | N/A | 2093 | N/A | N/A | N/A | N/A | N/A | N/A | 0.848245 |

| | | | | | | | | | | | | | |
|---|---|---|---|---|---|---|---|---|---|---|---|---|---|
| 959 | Chain B internal | 919 | N/A | N/A | N/A | N/A | N/A | N/A | N/A | N/A | N/A | N/A | 0.853978 |
| 960 | Chain B internal | 953 | 2205 | 829 | 2324 | N/A | N/A | 2171 | N/A | 2205 | 2180 | N/A | 0.848027 |
| 961 | Chain B/C interface | N/A | 2095 | N/A | 2128 | N/A | N/A | N/A | N/A | N/A | N/A | N/A | 0.792468 |
| 962 | Chain B surface | 967 | 2271 | 851 | 2352 | N/A | N/A | N/A | N/A | N/A | N/A | N/A | 0.832754 |
| 964 | Chain B surface | 1055 | 2220 | 783 | 2286 | 2139 | 2125 | 2185 | N/A | 2223 | 2195 | N/A | 0.870325 |
| 966 | Chain B surface | 1133 | N/A | N/A | N/A | N/A | N/A | N/A | N/A | N/A | N/A | N/A | 0.777318 |
| 967 | Chain B internal | 972 | 2287 | 778 | 2373 | 2180 | N/A | 2239 | 2096 | 2298 | 2240 | N/A | 0.844779 |
| 968 | ATP site | 937 | N/A | N/A | N/A | N/A | N/A | N/A | N/A | N/A | N/A | N/A | 0.908857 |
| 969 | Chain B/C interface | 1040 | 2262 | 857 | 2342 | N/A | 2146 | 2215 | N/A | 2256 | 2228 | N/A | 0.863627 |
| 970 | ATP site | 935 | N/A | N/A | N/A | N/A | N/A | N/A | N/A | N/A | N/A | 914 | 0.527132 |
| 971 | Chain B internal | 1038 | 2209 | N/A | 2268 | N/A | N/A | N/A | 2062 | 2207 | 2181 | N/A | 0.713339 |
| 972 | Chain B surface | 1110 | N/A | N/A | N/A | N/A | N/A | N/A | N/A | N/A | N/A | N/A | 0.643336 |
| 973 | Chain B internal | 921 | N/A | N/A | 2160 | N/A | N/A | N/A | N/A | N/A | N/A | N/A | 0.915434 |
| 974 | Chain B surface | 1015 | 2168 | 831 | 2225 | 2112 | 2094 | N/A | N/A | 2176 | N/A | N/A | 0.919044 |
| 975 | Chain B surface | 1246 | 2008 | N/A | 2010 | N/A | N/A | 2010 | N/A | N/A | 2014 | N/A | 0.688661 |
| 976 | Chain B surface | N/A | 2281 | N/A | N/A | N/A | N/A | N/A | N/A | N/A | N/A | N/A | 0.802981 |
| 980 | Chain B surface near ATP site | 1206 | 2037 | N/A | 2053 | N/A | N/A | N/A | N/A | N/A | 2047 | N/A | 0.803467 |
| 983 | Chain B surface | N/A | 2148 | N/A | 2215 | N/A | 2078 | N/A | N/A | 2151 | N/A | N/A | 0.716411 |
| 985 | Chain B internal | 975 | 2180 | 771 | 2237 | 2115 | 2102 | 2152 | 2054 | 2186 | 2159 | 906 | 0.783604 |
| 986 | Chain B surface | 1194 | 2015 | N/A | 2017 | 2013 | 2010 | 2016 | 2005 | 2019 | 2019 | N/A | 0.518359 |
| 987 | Chain B surface | 962 | 2204 | 819 | 2266 | 2131 | 2114 | 2170 | N/A | 2204 | 2179 | N/A | 0.837626 |
| 988 | Chain A/B interface | 998 in Chain A | 2118 | N/A | 2159 | 2078 | 2058 | 2088 | N/A | 2121 | 2102 | N/A | 0.813145 |
| 989 | Chain B surface near chain D | 907 | 2164 | N/A | 2216 | N/A | 2088 | 2135 | N/A | 2167 | 2143 | N/A | 0.823397 |
| 991 | Chain B internal | 1073 | 2177 | 773 | 2233 | N/A | N/A | 2147 | 2052 | 2181 | 2155 | 907 | 0.882765 |
| 992 | Chain B internal, in active site of Chain B | N/A | N/A | 793 | 2007 | N/A | N/A | N/A | N/A | N/A | N/A | N/A | 0.823473 |
| 993 | Chain B surface | 920 | 2195 | N/A | 2257 | N/A | N/A | N/A | N/A | N/A | N/A | N/A | 0.814889 |
| 995 | Chain B internal | 1137 | 2293 | 801 | 2377 | N/A | N/A | N/A | N/A | 2289 | 2245 | N/A | 0.879423 |
| 996 | Chain A/B interface | 1010 | N/A | N/A | N/A | N/A | N/A | N/A | N/A | N/A | N/A | N/A | 0.830866 |

| | | | | | | | | | | | | |
|---|---|---|---|---|---|---|---|---|---|---|---|---|
| 998 | Chain B internal | 998 | 2292 | 781 | 2376 | N/A | N/A | N/A | N/A | 2288 | 2244 | N/A | 0.866307 |
| 999 | Chain A/B interface near dATP site in Chain B | 992 | N/A | 836 | 2140 | 2070 | 2048 | 2078 | 2032 | 2117 | 2096 | N/A | 0.749727 |
| 1000 | Chain B surface | 947 | 2295 | N/A | 2363 | 2176 | 2158 | 2226 | N/A | 2290 | 2236 | N/A | 0.731393 |
| 1001 | Chain B internal | 1009 | 2285 | 812 | 2367 | 2171 | 2163 | 2229 | 2092 | 2279 | 2239 | N/A | 0.851709 |
| 1002 | Chain B internal | 1062 | 2288 | 775 | 2385 | 2174 | N/A | 2240 | N/A | 2283 | 2150 | N/A | 0.789614 |
| 1003 | Chain B surface | 1020 | 2250 | N/A | 2331 | N/A | N/A | N/A | 2082 | N/A | N/A | N/A | 0.81577 |
| 1004 | Chain B surface near Chain C | N/A | 2024 | N/A | N/A | N/A | 2015 | 2022 | N/A | 2024 | 2030 | N/A | 0.887079 |
| 1005 | Chain B surface | 956 | N/A | N/A | 2318 | N/A | N/A | N/A | N/A | N/A | N/A | N/A | 0.789051 |
| 1006 | Chain B internal | 1146 | N/A | N/A | 2117 | N/A | N/A | N/A | N/A | N/A | N/A | N/A | 0.865334 |
| 1007 | Chain B internal | 1079 | 2184 | 772 | 2246 | N/A | N/A | 2155 | 2056 | 2190 | 2079 | N/A | 0.911185 |
| 1009 | Chain B surface | N/A | 2203 | N/A | 2265 | 2129 | 2112 | 2168 | 2060 | 2202 | 2178 | N/A | 0.658598 |
| 1010 | Chain B internal | 1065 | 2176 | 777 | 2232 | N/A | 2098 | 2146 | 2051 | 2180 | 2154 | N/A | 0.840747 |
| 1011 | Chain B surface | 1115 | 2210 | N/A | 2272 | N/A | N/A | N/A | N/A | N/A | N/A | N/A | 0.926666 |
| 1012 | ATP site | 908 | N/A | N/A | N/A | N/A | 2001 | N/A | N/A | N/A | N/A | N/A | 0.60108 |
| 1013 | Chain B surface | N/A | 2069 | N/A | 2099 | N/A | N/A | N/A | N/A | 2071 | N/A | N/A | 0.530241 |
| 1014 | Chain B/C interface, in active site of Chain B | 1170 | N/A | N/A | N/A | N/A | N/A | N/A | N/A | N/A | N/A | N/A | 0.814412 |
| 1015 | Chain A/B interface | N/A | 2119 | N/A | N/A | N/A | N/A | N/A | N/A | N/A | 2103 | N/A | 0.834008 |
| 1017 | Chain B internal | 1053 | 2275 | 849 | 2359 | N/A | 2157 | N/A | 2062 | 2276 | 2233 | N/A | 0.681621 |
| 1019 | Chain B/C interface | N/A | 2259 | 792 | 2340 | 2156 | N/A | 2213 | 2085 | 2254 | 2217 | N/A | 0.76307 |
| 1020 | Chain B surface | N/A | N/A | N/A | 2089 | N/A | N/A | N/A | N/A | N/A | 2043 | N/A | 0.834909 |
| 1021 | Chain B surface | N/A | N/A | N/A | N/A | N/A | N/A | N/A | N/A | 2031 | N/A | 922 | 0.705861 |
| 1022 | Chain B surface near Chain D | N/A | N/A | N/A | N/A | 2091 | N/A | N/A | N/A | 2143 | 2121 | N/A | 0.572681 |
| 1023 | Chain B surface | N/A | N/A | N/A | N/A | N/A | N/A | N/A | N/A | 2034 | 2057 | N/A | 0.808067 |
| 1024 | Chain B surface | 941 | 2131 | N/A | 2172 | 2083 | 2065 | 2100 | N/A | 2134 | 2113 | N/A | 0.857824 |
| 1025 | Chain B internal, in active site of Chain B | 1076 | 2174 | N/A | 2150 | 2113 | 2097 | 2145 | 2050 | 2104 | 2152 | 905 | 0.911641 |
| 1026 | Chain B internal | 1081 | 2269 | 768 | 2349 | 2042 | 2151 | 2220 | 2021 | 2267 | 2227 | N/A | 0.880544 |

| | | | | | | | | | | | | |
|---|---|---|---|---|---|---|---|---|---|---|---|---|
| 1027 | Chain B/C interface, in active site of Chain B | N/A | N/A | N/A | N/A | N/A | N/A | N/A | N/A | N/A | N/A | 923 | 0.812897 |
| 1028 | Chain B internal | 1105 | 2254 | 767 | 2333 | N/A | N/A | N/A | N/A | N/A | 2215 | N/A | 0.860179 |
| 1029 | Chain B/C interface | 1195 | 2258 | N/A | 2338 | N/A | 2145 | N/A | 2084 | 2253 | N/A | N/A | 0.808222 |
| 1030 | Chain B surface | N/A | N/A | N/A | 2106 | N/A | N/A | N/A | 2078 | N/A | N/A | N/A | 0.62857 |
| 1031 | Chain B surface | N/A | N/A | N/A | 2174 | N/A | N/A | N/A | N/A | N/A | N/A | N/A | 0.823983 |
| 1032 | Chain B/C/D interface | N/A | N/A | N/A | N/A | N/A | 2090 | N/A | N/A | N/A | N/A | N/A | 0.867879 |
| 1033 | Chain B surface | N/A | 2082 | N/A | 2039 | N/A | N/A | N/A | 2026 | 2079 | 2070 | N/A | 0.825628 |
| 1034 | Chain B surface | 1131 | N/A | N/A | N/A | N/A | N/A | N/A | N/A | 2198 | N/A | N/A | 0.868278 |
| 1035 | Chain B surface | 1158 | 2056 | 820 | 2077 | 2041 | N/A | 2209 | 2020 | 2059 | 2056 | N/A | 0.880316 |
| 1036 | Chain B surface | 1259 | 2200 | 860 | 2263 | N/A | N/A | N/A | N/A | N/A | 2176 | N/A | 0.810512 |
| 1039 | Chain B surface | N/A | 2084 | 818 | 2109 | N/A | N/A | 2064 | 2027 | N/A | 2072 | N/A | 0.792416 |
| 1040 | Chain B/C interface | 1234 | 2123 | N/A | 2163 | 2094 | 2070 | 2011 | 2037 | 2145 | N/A | N/A | 0.821964 |
| 1041 | Chain B surface | 1017 | 2282 | 843 | 2368 | N/A | N/A | N/A | N/A | 2280 | N/A | N/A | 0.779527 |
| 1043 | Chain B internal | 1128 | 2299 | 798 | 2384 | 2179 | 2169 | 2238 | N/A | 2296 | N/A | N/A | 0.893644 |
| 1044 | Chain B internal, near active site in Chain B | N/A | 2187 | N/A | 2251 | 2120 | N/A | 2158 | 2058 | 2194 | 2166 | 909 | 0.866504 |
| 1045 | Chain B surface | 1036 | 2244 | 854 | 2319 | N/A | N/A | 2202 | N/A | 2243 | 2208 | N/A | 0.599221 |
| 1046 | Chain B/C interface | N/A | 2147 | N/A | 2189 | 2098 | 2077 | 2117 | N/A | 2150 | 2128 | N/A | 0.893947 |
| 1047 | Chain B surface | 964 | 2083 | 850 | 2108 | 2050 | N/A | N/A | N/A | N/A | N/A | N/A | 0.874061 |
| 1049 | Chain B surface | 1175 | N/A | N/A | 2149 | 2065 | 2110 | N/A | N/A | N/A | N/A | N/A | 0.826325 |
| 1050 | Chain B internal | 1002 | 2300 | 776 | 2386 | 2110 | 2092 | N/A | 2048 | 2297 | 2148 | 913 | 0.865613 |
| 1051 | Chain B surface | N/A | 2163 | N/A | N/A | 2107 | N/A | 2132 | N/A | 2164 | N/A | N/A | 0.79693 |
| 1053 | ATP site | 1139 | N/A | N/A | N/A | N/A | N/A | N/A | N/A | N/A | N/A | N/A | 0.683631 |
| 1054 | Chain B surface | N/A | 2159 | 826 | 2206 | 2103 | N/A | 2126 | N/A | 2159 | 2136 | N/A | 0.873995 |
| 1056 | Chain B/C interface | N/A | N/A | N/A | N/A | N/A | N/A | 2207 | 2080 | 2250 | N/A | N/A | 0.841839 |
| 1058 | Chain B surface | N/A | N/A | N/A | 2059 | N/A | 2021 | N/A | N/A | N/A | N/A | N/A | 0.907003 |
| 1059 | Chain B surface | 1243 | 2202 | N/A | 2264 | N/A | N/A | N/A | N/A | N/A | N/A | N/A | 0.768359 |
| 1061 | Chain B surface | 1124 | 2169 | 806 | 2223 | 2109 | 2093 | 2143 | N/A | 2177 | 2149 | N/A | 0.802367 |
| 1062 | Chain B surface | 1172 | 2154 | N/A | 2201 | N/A | 2084 | 2123 | 2045 | 2156 | 2134 | N/A | 0.779686 |
| 1063 | Chain B internal | 1071 | 2227 | N/A | 2292 | 2125 | 2126 | 2189 | N/A | 2227 | 2199 | 925 | 0.860836 |

| | | | | | | | | | | | | |
|---|---|---|---|---|---|---|---|---|---|---|---|---|
| 1064 | Chain B surface | 1257 | 2219 | N/A | 2285 | 2137 | N/A | 2184 | 2069 | 2219 | 2193 | N/A | 0.935318 |
| 1065 | Chain B surface | 1213 | N/A | N/A | N/A | N/A | N/A | N/A | N/A | 2050 | N/A | N/A | 0.723453 |
| 1066 | Chain B surface | 990 | 2156 | N/A | 2203 | 2102 | N/A | 2125 | N/A | 2158 | 2135 | N/A | 0.698498 |
| 1067 | Chain B internal | 981 | 2185 | 839 | 2244 | N/A | 2099 | N/A | 2057 | 2189 | 2156 | N/A | 0.786205 |
| 1068 | Chain B internal | N/A | 2207 | 799 | 2247 | 2118 | N/A | N/A | N/A | 2191 | 2164 | N/A | 0.859488 |
| 1072 | Chain B surface | N/A | 2233 | N/A | N/A | N/A | N/A | N/A | N/A | N/A | N/A | N/A | 0.906241 |
| 1074 | Chain B internal | 1173 | 2245 | 766 | 2325 | N/A | N/A | 2212 | N/A | 2246 | N/A | N/A | 0.855388 |
| 1075 | Chain B internal | 944 | 2238 | 821 | 2309 | N/A | N/A | N/A | N/A | 2235 | N/A | N/A | 0.785574 |
| 1077 | Chain B surface | N/A | 2188 | N/A | 2252 | N/A | N/A | N/A | N/A | N/A | N/A | N/A | 0.871595 |
| 1078 | Chain B internal | 1014 | 2086 | 764 | 2112 | 2054 | N/A | 2065 | N/A | 2081 | 2073 | N/A | 0.905924 |
| 1079 | Chain B surface | N/A | 2032 | N/A | 2045 | N/A | N/A | N/A | N/A | 2032 | 2037 | N/A | 0.770573 |
| 1080 | Chain B/C interface, in active site of Chain B | N/A | 2247 | N/A | N/A | N/A | N/A | N/A | N/A | N/A | N/A | N/A | 0.836946 |
| 1082 | Chain A/B interface | 1107 | 2132 | 804 | 2173 | 2084 | N/A | 2101 | N/A | 2135 | 2112 | N/A | 0.763866 |
| 1085 | Chain B surface | 1114 | 2153 | N/A | 2198 | N/A | N/A | N/A | N/A | 2052 | 2132 | N/A | 0.764913 |
| 1089 | Chain B surface | N/A | 2215 | 833 | 2279 | N/A | 2119 | N/A | N/A | 2216 | 2186 | N/A | 0.763415 |
| 1091 | Chain A/B interface | 1186 | N/A | N/A | N/A | N/A | N/A | N/A | N/A | N/A | N/A | N/A | 0.839061 |
| 1092 | Chain B surface, near Chains A and C | 1190 | 2139 | 763 | 2182 | 2092 | N/A | N/A | N/A | N/A | 2123 | 924 | 0.942153 |
| 1094 | Chain B internal | 1210 | 2067 | 788 | N/A | N/A | N/A | N/A | N/A | N/A | N/A | N/A | 0.741976 |
| 1095 | Chain B surface | 1186 | 2152 | N/A | 2199 | N/A | 2083 | 2122 | N/A | N/A | 2131 | N/A | 0.853415 |
| 1096 | Chain B surface | 1179 | 2165 | N/A | 2222 | N/A | N/A | 2141 | N/A | 2173 | 2147 | N/A | 0.783163 |
| 1097 | Chain B/C interface | 1199 | 2263 | N/A | N/A | N/A | N/A | N/A | N/A | 2257 | N/A | 911 | 0.915354 |
| 1098 | Chain B surface | 1082 | N/A | N/A | 2258 | N/A | 2108 | N/A | N/A | N/A | N/A | N/A | 0.797896 |
| 1099 | Chain B internal | 1060 | N/A | N/A | N/A | N/A | N/A | N/A | N/A | N/A | N/A | N/A | 0.866521 |
| 1100 | Chain B surface near Chain A | 1039 | 2138 | N/A | 2181 | 2090 | N/A | N/A | N/A | N/A | N/A | N/A | 0.865435 |
| 1101 | ATP site | 1229 | 2035 | N/A | 2052 | N/A | N/A | 2034 | 2016 | 2039 | 2040 | N/A | 0.807395 |
| 1104 | Chain B/C interface, near Y356 in Chain C | N/A | 2304 | N/A | 2391 | N/A | N/A | N/A | N/A | N/A | N/A | N/A | 0.738173 |
| 1105 | Chain A/B interface | N/A | 2092 | N/A | 2120 | 2059 | N/A | 2071 | N/A | 2085 | N/A | N/A | 0.882834 |

| | | | | | | | | | | | | | |
|---|---|---|---|---|---|---|---|---|---|---|---|---|---|
| 1106 | Chain B surface near Chain C | 1205 | 2240 | N/A | 2311 | 2145 | 2132 | 2198 | 2074 | 2239 | 2205 | N/A | 0.600165 |
| 1107 | Chain B surface | 1167 | N/A | N/A | 2068 | N/A | N/A | N/A | N/A | N/A | N/A | N/A | 0.86576 |
| 1108 | Chain B surface | 1102 | 2041 | N/A | 2056 | N/A | N/A | N/A | N/A | N/A | N/A | N/A | 0.823467 |
| 1109 | Chain A/B/C interface | 1051 | 2124 | N/A | 2162 | N/A | N/A | N/A | N/A | N/A | 2105 | N/A | 0.891692 |
| 1110 | Chain B/C/D interface | N/A | N/A | N/A | N/A | N/A | N/A | N/A | N/A | 2169 | N/A | N/A | 0.807915 |
| 1111 | Chain B surface near Chain C | 1252 | 2031 | N/A | 2006 | 2025 | 2017 | 2029 | 2014 | 2030 | 2035 | N/A | 0.85353 |
| 1112 | Chain B surface | 901 | 2055 | N/A | 2076 | 2040 | 2031 | N/A | N/A | N/A | 2055 | N/A | 0.85184 |
| 1113 | Chain B internal | 1070 | 2085 | 811 | 2111 | N/A | N/A | N/A | N/A | 2080 | N/A | N/A | 0.877702 |
| 1115 | Chain B/C interface | N/A | N/A | N/A | N/A | 2171 | N/A | N/A | N/A | N/A | N/A | N/A | 0.807249 |
| 1116 | Chain B surface | 1117 | 2054 | 842 | 2075 | N/A | N/A | N/A | N/A | 2058 | N/A | N/A | 0.920671 |
| 1117 | Chain B/C interface | N/A | 2028 | N/A | 2038 | 2024 | N/A | 2028 | N/A | N/A | N/A | N/A | 0.883843 |
| 1118 | Chain B surface near dATP site in Chain B | 1123 | 2104 | 844 | 2137 | N/A | N/A | N/A | N/A | 2097 | 2081 | N/A | 0.823314 |
| 1120 | Chain B/C interface | N/A | 2136 | N/A | 2177 | N/A | N/A | 2102 | N/A | 2137 | 2115 | N/A | 0.794066 |
| 1121 | Chain B internal | 1301 | N/A | N/A | N/A | N/A | N/A | N/A | N/A | N/A | N/A | N/A | 0.869759 |
| 1122 | Chain B surface | 1100 | N/A | N/A | N/A | N/A | N/A | N/A | N/A | N/A | N/A | N/A | 0.905213 |
| 1123 | Chain B/C interface | N/A | 2098 | 840 | 2344 | 2158 | 2018 | 2217 | 2086 | 2092 | 2036 | N/A | 0.627721 |
| 1125 | Chain B surface | N/A | N/A | N/A | N/A | N/A | N/A | N/A | N/A | 2166 | 2142 | N/A | 0.90189 |
| 1126 | Chain B surface | N/A | N/A | N/A | 2057 | N/A | N/A | N/A | N/A | N/A | N/A | N/A | 0.82522 |
| 1127 | Chain B surface | 960 | 2241 | N/A | 2313 | N/A | 2133 | 2199 | 2075 | 2240 | 2206 | N/A | 0.815075 |
| 1128 | ATP site | 1165 | N/A | N/A | N/A | N/A | N/A | N/A | N/A | N/A | 2092 | N/A | 0.827876 |
| 1129 | dATP site in Chain B | 1187 | N/A | N/A | N/A | N/A | N/A | N/A | N/A | N/A | N/A | N/A | 0.800928 |
| 1130 | Chain B internal | 1160 | N/A | N/A | N/A | N/A | N/A | N/A | N/A | N/A | N/A | N/A | 0.800537 |
| 1131 | Chain B surface | 1129 | N/A | N/A | 2051 | 2028 | 2019 | N/A | N/A | N/A | N/A | N/A | 0.739785 |
| 1133 | Chain B surface | 1135 | 2229 | N/A | 2295 | 2126 | 2107 | N/A | N/A | 2197 | 2173 | N/A | 0.913668 |
| 1135 | Chain B internal | 948 | N/A | N/A | N/A | N/A | N/A | N/A | N/A | N/A | N/A | N/A | 0.799553 |
| 1136 | Chain B/C interface | 1183 | 2289 | N/A | 2388 | N/A | N/A | N/A | N/A | 2285 | N/A | N/A | 0.870009 |
| 1137 | Chain B/C interface | N/A | 2019 | N/A | 2024 | N/A | 2012 | N/A | N/A | N/A | N/A | N/A | 0.886737 |
| 1138 | Chain A/B interface | 1087 | 2106 | N/A | 2140 | 2064 | N/A | 2078 | 2029 | 2100 | N/A | N/A | 0.900034 |

| | | | | | | | | | | | | | |
|---|---|---|---|---|---|---|---|---|---|---|---|---|---|
| 1139 | Chain B/C interface | N/A | N/A | N/A | N/A | N/A | N/A | N/A | N/A | 2028 | N/A | N/A | 0.849285 |
| 1140 | Chain B/C interface | 973 | N/A | N/A | N/A | N/A | 2066 | N/A | N/A | N/A | N/A | N/A | 0.745589 |
| 1142 | Chain A/B interface near dATP site in Chain A | 1141 | 2111 | N/A | 2148 | N/A | N/A | N/A | N/A | N/A | 2083 | N/A | 0.754389 |
| 1143 | Chain B internal | 980 | 2239 | N/A | 2310 | N/A | N/A | N/A | N/A | 2237 | N/A | N/A | 0.89543 |
| 1144 | Chain A/B interface and dATP site in Chain B | 1241 | 2110 | N/A | N/A | N/A | N/A | 2079 | N/A | 2116 | N/A | N/A | 0.846162 |
| 1145 | Chain B surface | N/A | 2158 | N/A | 2208 | N/A | N/A | 2128 | N/A | 2160 | 2125 | N/A | 0.8945 |
| 1146 | Chain B surface | 1089 | 2224 | N/A | 2289 | N/A | N/A | 2187 | N/A | 2225 | 2197 | N/A | 0.801791 |
| 1148 | Chain B/C interface | N/A | 2025 | N/A | 2034 | N/A | N/A | N/A | N/A | 2026 | N/A | N/A | 0.90239 |
| 1149 | Chain B internal | N/A | N/A | N/A | 2093 | N/A | N/A | N/A | N/A | 2066 | 2074 | N/A | 0.832765 |
| 1152 | Chain B surface | 1045 | 2284 | N/A | 2369 | 2172 | 2162 | 2230 | 2093 | 2281 | 2238 | N/A | 0.750837 |
| 1153 | Chain B surface | 1270 | 2107 | N/A | 2141 | N/A | N/A | N/A | N/A | 2101 | N/A | N/A | 0.850887 |
| 1157 | Chain B/C interface | N/A | 2268 | N/A | 2036 | 2023 | 2152 | 2219 | 2089 | 2266 | 2224 | N/A | 0.89614 |
| 1158 | Chain B surface | 1239 | N/A | N/A | 2202 | N/A | N/A | N/A | N/A | N/A | N/A | N/A | 0.902143 |
| 1159 | Chain B surface near chain C | N/A | 2296 | N/A | 2378 | N/A | N/A | N/A | N/A | 2291 | N/A | N/A | 0.900754 |
| 1160 | Chain B surface | N/A | N/A | N/A | 2269 | N/A | N/A | N/A | N/A | N/A | N/A | N/A | 0.800748 |
| 1161 | Chain A/B interface | 1207 | 2108 | 832 | 2143 | N/A | N/A | N/A | N/A | N/A | N/A | 921 | 0.853925 |
| 1163 | Chain B surface | 1282 | N/A | N/A | N/A | N/A | N/A | N/A | N/A | N/A | N/A | N/A | 0.798455 |
| 1166 | Chain B surface near Chain C | N/A | N/A | N/A | N/A | N/A | N/A | N/A | N/A | N/A | 2256 | N/A | 0.896872 |
| 1167 | Chain B surface | 1143 | N/A | N/A | 2245 | N/A | N/A | N/A | N/A | N/A | N/A | N/A | 0.674744 |
| 1170 | Chain A/B interface | 1111 | 2181 | 800 | 2238 | N/A | N/A | 2151 | 2055 | 2185 | N/A | N/A | 0.773511 |
| 1171 | Chain B surface | 1244 | N/A | N/A | N/A | N/A | N/A | N/A | N/A | N/A | N/A | N/A | 0.819553 |
| 1172 | Chain B surface | N/A | 2072 | 815 | 2287 | 2138 | N/A | N/A | 2070 | 2209 | 2065 | N/A | 0.91076 |
| 1173 | Chain B surface | 1306 | N/A | N/A | 2175 | N/A | N/A | N/A | N/A | N/A | 2111 | N/A | 0.812918 |
| 1175 | Chain B/C interface | N/A | 2096 | N/A | 2129 | N/A | N/A | N/A | N/A | N/A | N/A | N/A | 0.73625 |
| 1176 | Chain B internal | 1231 | 2170 | 814 | 2228 | N/A | N/A | N/A | N/A | N/A | N/A | N/A | 0.722855 |
| 1179 | Chain B internal in active site of Chain B | 1269 | 2090 | N/A | N/A | N/A | N/A | N/A | N/A | N/A | N/A | N/A | 0.890412 |

| | | | | | | | | | | | | | |
|---|---|---|---|---|---|---|---|---|---|---|---|---|---|
| 1180 | Chain B surface | 1230 | N/A | N/A | N/A | N/A | N/A | N/A | N/A | N/A | N/A | N/A | 0.679166 |
| 1181 | Chain B/D interface | N/A | N/A | N/A | N/A | N/A | N/A | 2108 | N/A | N/A | N/A | N/A | 0.787115 |
| 1182 | Chain A/B/C interface | 1228 | N/A | N/A | N/A | N/A | N/A | N/A | N/A | N/A | N/A | N/A | 0.720529 |
| 1183 | Chain B surface | 1317 | N/A | 790 | N/A | 2133 | 2039 | 2177 | 2063 | N/A | N/A | N/A | 0.742122 |
| 1184 | Chain B surface | N/A | N/A | N/A | N/A | 2123 | N/A | N/A | N/A | N/A | N/A | N/A | 0.635074 |
| 1185 | Chain B surface | 1309 | N/A | N/A | N/A | N/A | 2117 | N/A | N/A | N/A | N/A | N/A | 0.870323 |
| 1188 | Chain B surface | 1083 | 2228 | N/A | 2294 | 2142 | N/A | 2191 | 2072 | N/A | 2200 | N/A | 0.741399 |
| 1189 | Chain B surface | N/A | N/A | N/A | N/A | 2124 | N/A | 2162 | N/A | N/A | 2172 | N/A | 0.741347 |
| 1192 | Chain B surface | 1314 | N/A | N/A | N/A | N/A | N/A | N/A | N/A | N/A | N/A | N/A | 0.940014 |
| 1193 | Chain B surface | 1244 | N/A | N/A | N/A | N/A | N/A | N/A | N/A | N/A | N/A | N/A | 0.832208 |
| 1194 | Chain B surface | 1298 | N/A | N/A | N/A | N/A | N/A | N/A | N/A | N/A | N/A | N/A | 0.939378 |
| 1195 | Chain A/B interface near Chain C | 1313 | 2120 | N/A | 2042 | N/A | N/A | 2036 | N/A | N/A | 2046 | N/A | 0.812362 |
| 1196 | Chain B surface | 1292 | N/A | N/A | N/A | N/A | N/A | N/A | N/A | N/A | N/A | N/A | 0.842742 |
| 901 in Chain A | Chain A/B interface | N/A | N/A | N/A | N/A | 2076 | N/A | N/A | N/A | N/A | N/A | N/A | 0.814448 |
| 974 in Chain A | Chain A/B interface near dATP site in Chain B | 1186 | N/A | N/A | N/A | N/A | N/A | N/A | N/A | N/A | N/A | N/A | 0.815534 |
| 982 in Chain A | Chain A/B interface | 1042 | 2109 | N/A | 2139 | N/A | N/A | N/A | N/A | 2102 | N/A | N/A | 0.792635 |
| 1006 in Chain A | Chain A/B/C interface | N/A | N/A | N/A | N/A | N/A | N/A | N/A | N/A | N/A | N/A | 912 | 0.873525 |
| 1007 in Chain A | Chain A near dATP site in Chain A | 1091 | N/A | N/A | N/A | N/A | N/A | N/A | N/A | N/A | N/A | N/A | 0.874661 |
| 1043 in Chain A | Chain A/B interface | N/A | 2012 | N/A | N/A | N/A | N/A | N/A | N/A | N/A | N/A | N/A | 0.735162 |
| 1048 in Chain A | Chain A surface near dATP site in Chain A | N/A | N/A | N/A | N/A | N/A | N/A | N/A | N/A | N/A | 2114 in Chain A | N/A | 0.759753 |
| 503 in Chain C | Chain B/C interface | 1261 | 2130 | N/A | 2171 | N/A | N/A | N/A | N/A | N/A | N/A | N/A | 0.851865 |
| 506 in Chain C | Chain B/C interface | 1164 | 2001 in Chain E | N/A | 2002 in Chain E | N/A | N/A | 2234 | N/A | 2001 | 2002 in Chain E | N/A | 0.813433 |
| 507 in Chain C | Chain B/C interface near Y356 in Chain C | N/A | 2303 | N/A | N/A | 2182 | 2170 | 2140 | N/A | 2302 | 2252 | N/A | 0.815922 |
| 514 in Chain C | Chain B/C interface | N/A | N/A | N/A | 2005 in Chain E | N/A | N/A | N/A | N/A | N/A | N/A | N/A | 0.69363 |
| 537 in Chain C | Chain B/C interface | 1161 | 2277 | N/A | 2362 | N/A | 2159 | 2227 | 2091 | 2277 | N/A | N/A | 0.745016 |

| | | | | | | | | | | | | | |
|---|---|---|---|---|---|---|---|---|---|---|---|---|---|
| 544 in Chain C | Chain B/C interface | N/A | N/A | N/A | 2029 | N/A | N/A | N/A | N/A | N/A | 2023 | N/A | 0.855975 |
| 553 in Chain C | Chain A/B/C interface | N/A | N/A | N/A | 2058 | N/A | N/A | N/A | N/A | N/A | N/A | N/A | 0.775084 |
| 558 in Chain C | Chain B/C interface | N/A | N/A | N/A | 2393 | N/A | N/A | 2244 | N/A | 2305 | 2234 | N/A | 0.730446 |
| 564 in Chain C | Chain B/C interface in the active site of Chain B | N/A | N/A | N/A | N/A | N/A | N/A | N/A | N/A | 2299 | N/A | N/A | 0.794003 |
| 573 in Chain C | Chain B/C/D interface | N/A | N/A | N/A | 2218 | N/A | N/A | 2137 | N/A | 2171 | N/A | N/A | 0.873128 |
| 580 in Chain C | Chain A/B/C interface | 1030 | N/A | N/A | N/A | N/A | N/A | N/A | N/A | N/A | N/A | N/A | 0.731432 |
| 591 in Chain C | Chain B/C interface in active site of Chain B | 1305 | N/A | N/A | N/A | N/A | N/A | 2098 | N/A | 2131 | N/A | N/A | 0.915874 |
| 594 in Chain C | Chain B/C interface near Y356 in Chain B | N/A | N/A | N/A | N/A | N/A | N/A | 2139 | N/A | 2172 | 2146 | N/A | 0.896096 |
| 603 in Chain C | Chain A/B/C interface | N/A | 2127 | 816 | N/A | N/A | 2061 | 2094 | N/A | 2127 | N/A | N/A | 0.811685 |
| 604 in Chain C | Chain B/C interface | N/A | N/A | N/A | N/A | 2163 | N/A | N/A | N/A | N/A | N/A | N/A | 0.904331 |

*unless the Chain identity is otherwise specified

**Table S5. List of waters bound to chain C residues (β) that have corresponding water molecules in one or more crystal structures.**

| Water Number in Chain C (β)* | Location (relative to Chain C) | 2XOF – Water Number in Chain A (Chain A aligned by C-alphas) | 5CI0 – Water Number in Chain A (Chain A aligned by C-alphas) | 5CI1 – Water Number in Chain A (Chain A aligned by C-alphas) | 5CI2 – Water Number in Chain A (Chain A aligned by C-alphas) | 5CI3 – Water Number in Chain A (Chain A aligned by C-alphas) | 5CI4 – Water Number in Chain A (Chain A aligned by C-alphas) | 5CNS – Water Number in Chain H (Chain H aligned by C-alphas) | 5CNS – Water Number in Chain G (Chain G aligned by C-alphas) | Qscore |
|---|---|---|---|---|---|---|---|---|---|---|
| 501 | Chain C surface | 2155 | 635 | 621 | 640 | 646 | 669 | N/A | N/A | 0.69721 |
| 504 | Chain C surface | 2052 | N/A | N/A | N/A | N/A | N/A | N/A | N/A | 0.748096 |
| 505 | Chain C internal | N/A | 533 | 509 | 516 | 508 | 506 | N/A | N/A | 0.827597 |
| 508 | Chain C internal | 2119 | 562 | 517 | 532 | 553 | 527 | 608 | 607 | 0.843476 |
| 510 | Chain B/C interface | 2189 | N/A | N/A | N/A | N/A | N/A | N/A | N/A | 0.882875 |
| 512 | Chain C internal | 2072 | 511 | 548 | 525 | 569 | 516 | N/A | N/A | 0.812895 |
| 515 | Chain C internal | 2013 | 588 | 645 | 595 | 586 | 577 | N/A | 610 | 0.8421 |
| 516 | Chain B/C interface | N/A | 678 | 505 | 581 | 679 | 710 | N/A | N/A | 0.827894 |
| 517 | Chain C internal | 2135 | 535 | 545 | 539 | 552 | 558 | 609 | 604 | 0.743141 |
| 518 | Chain C surface | N/A | N/A | 669 | N/A | N/A | 633 | N/A | N/A | 0.885654 |
| 522 | Chain C internal | 2132 | 605 | 549 | 592 | 604 | 603 | 603 | N/A | 0.907498 |
| 523 | Chain C surface | N/A | 671 | 698 | N/A | N/A | 680 | N/A | N/A | 0.864723 |
| 524 | Chain C surface | N/A | 670 | 676 | 650 | 579 | 667 | N/A | N/A | 0.761002 |
| 525 | Chain C surface | 2177 | 617 | 510 | 537 | 551 | 637 | N/A | N/A | 0.789202 |
| 527 | Chain C surface | 2063 | 568 | 531 | 582 | 581 | 624 | N/A | N/A | 0.790612 |
| 528 | Chain C surface near Chain A | N/A | N/A | N/A | N/A | 512 | 502 | N/A | N/A | 0.799775 |
| 529 | Chain B/C interface | 2163 | 508 | 558 | 509 | 583 | 574 | N/A | N/A | 0.82611 |
| 530 | Chain C internal | 2137 | 586 | 597 | 605 | 601 | 610 | N/A | 612 | 0.894852 |
| 531 | Chain C/D interface | 2078 | 513 | 546 | 571 | 572 | 524 | N/A | N/A | 0.879885 |
| 532 | Chain C surface | 2058 | 536 | N/A | 562 | 518 | 553 | N/A | N/A | 0.725901 |
| 533 | Chain C/D interface | N/A | 679 | 530 | 558 | 532 | 568 | N/A | N/A | 0.838667 |
| 534 | Chain C surface | 2178 | 650 | 639 | 600 | 574 | 530 | N/A | N/A | 0.567009 |
| 536 | Chain B/C interface | N/A | 570 | 584 | 594 | 635 | 534 | N/A | N/A | 0.840254 |
| 538 | Chain C surface | 2151 | 645 | 659 | 644 | 642 | 617 | N/A | N/A | 0.74493 |
| 541 | Chain C internal | 2116 | 623 | 623 | 626 | 585 | 627 | N/A | N/A | 0.803239 |

| | | | | | | | | | |
|---|---|---|---|---|---|---|---|---|---|
| 545 | Chain B/C interface | 2162 | 614 | 577 | 646 | 641 | 620 | N/A | N/A | 0.863681 |
| 546 | Chain B/C interface | N/A | 689 | 666 | 688 | 684 | 608 | N/A | N/A | 0.84668 |
| 547 | Chain C internal | 2023 | 607 | 652 | 658 | 628 | 609 | N/A | 613 | 0.889909 |
| 548 | Chain C/D interface | N/A | 555 | 628 | 623 | 503 | 645 | N/A | N/A | 0.903091 |
| 551 | Chain B/C interface | 2133 | N/A | N/A | N/A | N/A | N/A | N/A | N/A | 0.846489 |
| 552 | Chain C surface near Chain B | 2032 | 578 | 604 | 611 | 576 | 660 | N/A | N/A | 0.90908 |
| 554 | Chain C surface | 2031 | 675 | 660 | 659 | 634 | 643 | N/A | N/A | 0.559846 |
| 555 | Chain C surface | 2165 | 616 | 646 | 616 | 645 | 592 | N/A | N/A | 0.88504 |
| 557 | Chain C/D interface | 2081 | 629 | N/A | N/A | 589 | N/A | N/A | N/A | 0.886191 |
| 559 | Chain C surface | 2067 | 661 | 624 | 548 | 677 | 677 | N/A | N/A | 0.806189 |
| 560 | Chain C surface | 2051 | N/A | N/A | N/A | N/A | N/A | N/A | N/A | 0.845962 |
| 566 | Chain C internal | 2143 | 549 | 611 | 565 | 538 | 541 | N/A | N/A | 0.753506 |
| 568 | Chain C/D interface | N/A | 630 | 626 | 641 | 682 | 666 | N/A | N/A | 0.813004 |
| 569 | Chain C internal | 2117 | 564 | 593 | 535 | 570 | 536 | N/A | N/A | 0.848924 |
| 571 | Chain C/B interface | 2014 | 677 | 690 | 687 | 698 | 676 | N/A | N/A | 0.861487 |
| 574 | Chain C surface | 2138 | 636 | 544 | 609 | 575 | 539 | N/A | N/A | 0.681999 |
| 575 | Chain B/C interface | 2018 | 565 | 529 | 577 | 632 | 551 | N/A | N/A | 0.887211 |
| 576 | Chain B/C interface | N/A | 632 | 550 | 612 | 588 | 569 | N/A | N/A | 0.84851 |
| 579 | Chain C surface | 2101 | 503 | 603 | 512 | 511 | 511 | N/A | N/A | 0.869885 |
| 581 | Chain C surface | 2171 | 639 | 671 | 645 | 622 | 646 | N/A | N/A | 0.851588 |
| 583 | Chain B/C interface | 2175 | 600 | 600 | 578 | 547 | 586 | N/A | N/A | 0.782821 |
| 585 | Chain C/D interface | N/A | 596 | 585 | 637 | 623 | 616 | N/A | N/A | 0.817923 |
| 587 | Chain C/D interface | N/A | 633 | 683 | 675 | 671 | 638 | N/A | N/A | 0.866949 |
| 588 | Chain C internal | 2149 | 527 | 552 | 544 | 599 | 543 | N/A | N/A | 0.809273 |
| 590 | Chain C surface | 2110 | 631 | 658 | 663 | 656 | 651 | N/A | N/A | 0.835024 |
| 593 | Chain C surface | 2026 | 667 | 633 | 649 | 666 | 682 | N/A | N/A | 0.789744 |
| 595 | Chain C/D interface | 2083 | 581 | 504 | 575 | 540 | 533 | N/A | N/A | 0.903847 |
| 598 | Chain C internal | 2144 | N/A | N/A | N/A | N/A | N/A | N/A | N/A | 0.792783 |

| | | | | | | | | | | |
|---|---|---|---|---|---|---|---|---|---|---|
| 601 | Chain C internal | 2036 | 643 | 650 | 629 | 560 | 566 | N/A | N/A | 0.891445 |
| 1028 in Chain A | Chain A/C interface | 2046 | 609 | 632 | 614 | 611 | N/A | N/A | N/A | 0.817144 |
| 1048 in Chain B | Chain B/C interface | 2042 | N/A | N/A | N/A | N/A | N/A | N/A | N/A | 0.921024 |
| 1057 in Chain B | Chain B/C interface | N/A | 697 | 701 | 700 | 706 | 703 | N/A | N/A | 0.868964 |
| 1073 in Chain B | Chain B/C interface | N/A | 693 | 689 | 694 | 694 | 700 | N/A | N/A | 0.77985 |
| 506 in Chain D | Chain C/D interface | 2093 | N/A | N/A | N/A | N/A | N/A | N/A | N/A | 0.793798 |
| 525 in Chain D | Chain C/D interface | 2079 | N/A | 620 | 632 | N/A | 658 | N/A | N/A | 0.869497 |
| 527 in Chain D | Chain C/D interface | 2104 | N/A | 572 | 536 | 521 | N/A | N/A | N/A | 0.907282 |
| 537 in Chain D | Chain C/D interface | N/A | 602 | 516 | 608 | 607 | 601 | N/A | N/A | 0.924188 |
| 559 in Chain D | Chain C/D interface | 2043 | N/A | N/A | N/A | N/A | N/A | N/A | N/A | 0.905344 |

*unless the Chain identity is otherwise specified

**Table S6. List of waters bound to chain D residues (β') that have corresponding water molecules in one or more crystal structures.**

| Water Number in Chain D (β')* | Location (relative to Chain D) | 2XOF – Water Number in Chain B (Chain B aligned by C-alphas) | 5CI0 – Water Number in Chain A (Chain A aligned by C-alphas) | 5CI1 – Water Number in Chain A (Chain A aligned by C-alphas) | 5CI2 – Water Number in Chain A (Chain A aligned by C-alphas) | 5CI3 – Water Number in Chain A (Chain A aligned by C-alphas) | 5CI4 – Water Number in Chain A (Chain A aligned by C-alphas) | 5CNS – Water Number in Chain E (Chain E aligned by C-alphas) | 5CNS – Water Number in Chain F (Chain F aligned by C-alphas) | Qscore |
|---|---|---|---|---|---|---|---|---|---|---|
| 501 | Chain D surface | 2001 | N/A | N/A | N/A | N/A | N/A | N/A | N/A | 0.739029 |
| 502 | Chain D internal | 2066 | 511 | 548 | 525 | 569 | 516 | N/A | N/A | 0.842776 |
| 507 | Chain D internal | 2115 | 535 | 545 | 539 | 552 | 558 | 609 | 609 | 0.755458 |
| 508 | Chain D surface | 2167 | 644 | 649 | 671 | 667 | 642 | N/A | N/A | 0.589915 |
| 510 | Chain C/D interface | 2074 | 679 | 530 | 558 | 532 | 568 | N/A | N/A | 0.868074 |
| 511 | Chain D surface | N/A | 645 | 659 | 644 | 642 | 617 | N/A | N/A | 0.775572 |
| 512 | Chain C/D interface | 2026 | 615 | 653 | 653 | 624 | 593 | N/A | N/A | 0.79454 |
| 513 | Chain D internal | 2105 | 569 | 599 | 603 | 592 | 596 | N/A | N/A | 0.849179 |
| 514 | Chain D surface | 2127 | 549 | 611 | 565 | 538 | 541 | N/A | N/A | 0.840527 |
| 515 | Chain D surface | 2122 | 695 | 694 | 690 | 704 | 694 | N/A | N/A | 0.60209 |
| 516 | Chain D surface near Chain B | N/A | 578 | 604 | 611 | 576 | 660 | N/A | N/A | 0.878622 |
| 517 | Chain D surface | 2169 | 622 | 622 | 555 | 643 | N/A | N/A | N/A | 0.774024 |
| 518 | Chain D surface | 2089 | N/A | N/A | N/A | N/A | N/A | N/A | N/A | 0.781212 |
| 519 | Chain D surface | 2103 | 564 | 593 | 535 | 570 | 536 | N/A | 610 | 0.752963 |
| 520 | Chain D surface | 2170 | N/A | N/A | N/A | N/A | N/A | N/A | N/A | 0.617013 |
| 521 | Chain B/C/D interface | 2041 | 658 | 648 | 634 | 664 | 632 | N/A | N/A | 0.785889 |
| 522 | Chain D surface | 2113 | 681 | 687 | 666 | 647 | 670 | N/A | N/A | 0.922288 |
| 523 | Chain D surface | 2112 | 605 | 549 | 592 | 604 | 603 | N/A | N/A | 0.914072 |
| 524 | Chain D internal | 2118 | 586 | 597 | 605 | 601 | 610 | N/A | N/A | 0.742284 |
| 525 | Chain C/D interface | 2079 in Chain A | 629 | N/A | N/A | 589 | N/A | N/A | N/A | 0.869497 |
| 526 | Chain D surface | 2080 | 680 | 670 | 639 | 637 | 625 | N/A | N/A | 0.804296 |
| 527 | Chain C/D interface | 2104 in Chain A | 574 | N/A | N/A | N/A | 580 | N/A | N/A | 0.907282 |
| 528 | Chain D internal | 2018 | 507 | 567 | 502 | 554 | 515 | 604 | N/A | 0.811967 |

| | | | | | | | | | | |
|---|---|---|---|---|---|---|---|---|---|---|
| 530 | Chain D internal | 2021 | 588 | 645 | 595 | 586 | 577 | 608 | N/A | 0.837587 |
| 533 | Chain C/D interface | 2075 | 596 | 585 | 637 | 623 | 616 | N/A | 612 | 0.855745 |
| 534 | Chain C/D interface | 2020 | 628 | 601 | 601 | 595 | 647 | N/A | N/A | 0.834769 |
| 535 | Chain B/D interface | N/A | 537 | 634 | 615 | 662 | 681 | N/A | N/A | 0.767379 |
| 536 | Chain C/D interface | 2064 | 603 | 602 | 622 | 587 | 635 | N/A | N/A | 0.880767 |
| 537 | Chain C/D interface | 2069 | 602 | 516 | 608 | 607 | 601 | N/A | N/A | 0.924188 |
| 539 | Chain D surface | 2152 | 593 | 655 | 654 | 631 | 659 | N/A | N/A | 0.74035 |
| 540 | Chain D surface | N/A | 665 | 608 | 618 | 633 | 679 | N/A | N/A | 0.611122 |
| 541 | Chain D surface | 2097 | 503 | 603 | 512 | 511 | 511 | N/A | N/A | 0.801963 |
| 542 | Chain D surface | N/A | 510 | 561 | 513 | 559 | 549 | N/A | N/A | 0.849633 |
| 544 | Chain D surface | 2053 | 522 | 524 | 523 | 509 | 531 | N/A | N/A | 0.758194 |
| 545 | Chain D surface | 2136 | 635 | 621 | 640 | 646 | 669 | N/A | N/A | 0.629037 |
| 547 | Chain C/D interface | 2009 | 686 | 642 | 682 | 691 | 685 | N/A | N/A | 0.836917 |
| 549 | Chain C/D interface | 2086 | 659 | 644 | 617 | 640 | 673 | N/A | N/A | 0.670254 |
| 551 | Chain B/D interface | N/A | 570 | 584 | 594 | 635 | 534 | N/A | N/A | 0.735835 |
| 552 | Chain D surface | 2161 | 600 | 600 | 578 | 547 | 586 | N/A | N/A | 0.775324 |
| 553 | Chain C/D interface | N/A | 555 | 628 | 623 | 503 | 645 | N/A | N/A | 0.871854 |
| 554 | Chain D surface | 2114 | N/A | N/A | N/A | N/A | N/A | N/A | N/A | 0.817378 |
| 555 | Chain D internal | 2098 | N/A | N/A | N/A | N/A | N/A | N/A | N/A | 0.631756 |
| 556 | Chain D surface | 2111 | 546 | 590 | 563 | 598 | 599 | N/A | N/A | 0.826425 |
| 557 | Chain D surface | 2015 | 631 | 658 | 663 | 656 | 651 | N/A | N/A | 0.90321 |
| 558 | Chain D internal | 2012 | 607 | 652 | 658 | 628 | 609 | N/A | N/A | 0.871309 |
| 559 | Chain C/D interface | 2043 in Chain A | 633 | 683 | 675 | 671 | 638 | N/A | N/A | 0.905344 |
| 560 | Chain D internal | 2035 | 643 | 650 | 629 | 560 | 566 | N/A | N/A | 0.814984 |
| 561 | Chain D surface near Chain C | 2005 | N/A | N/A | N/A | N/A | N/A | N/A | N/A | 0.863626 |
| 563 | Chain D surface | 2100 | 623 | 623 | 626 | 585 | 627 | N/A | N/A | 0.811471 |

| | | | | | | | | | | |
|---|---|---|---|---|---|---|---|---|---|---|
| 564 | Chain C/D interface | 2093 in Chain A | N/A | N/A | N/A | N/A | N/A | N/A | N/A | 0.711731 |
| 565 | Chain D near Chain B | N/A | 689 | 666 | 688 | 684 | 608 | N/A | N/A | 0.78647 |
| 567 | Chain C/D interface | N/A | 505 | N/A | 528 | N/A | N/A | N/A | N/A | 0.814825 |
| 569 | Chain C/D interface | N/A | 704 | 699 | 699 | 702 | 704 | N/A | N/A | 0.7974 |
| 570 | Chain D surface near Chain B | N/A | 678 | 505 | 581 | 679 | 710 | N/A | N/A | 0.683878 |

*unless the Chain identity is otherwise specified

**Table S7. List of waters bound to any chain in the structure found using the "Find Waters" function in Coot (see SI for details) that do not have corresponding water molecules in the crystal structures analyzed above.**

| Water Number and Chain from "Find Waters" not accounted for by crystal structures | Location | Qscore |
|---|---|---|
| 1015 in Chain A | Chain A internal | 0.834147 |
| 936 in Chain B | Chain B/C interface | 0.758996 |
| 955 in Chain B | Chain B/C interface | 0.902583 |
| 963 in Chain B | Chain B surface | 0.749244 |
| 982 in Chain B | Chain B/C interface and Chain B active site | 0.912371 |
| 984 in Chain B | Chain B active site | 0.864741 |
| 1037 in Chain B | Chain B surface | 0.893012 |
| 1038 in Chain B | Chain B surface | 0.831865 |
| 1076 in Chain B | Chain B/C interface | 0.902976 |
| 1081 in Chain B | Chain B/D interface | 0.925554 |
| 1086 in Chain B | Chain B surface | 0.84617 |
| 1093 in Chain B | Chain B/C interface | 0.781017 |
| 1102 in Chain B | Chain B/C interface | 0.893715 |
| 1124 in Chain B | Chain B surface near chain D | 0.850708 |
| 1156 in Chain B | Chain B/C interface | 0.809696 |
| 1165 in Chain B | Chain B/C interface | 0.823532 |
| 1177 in Chain B | Chain B/C interface | 0.836857 |
| 1178 in Chain B | Chain A/B/C interface | 0.860573 |
| 509 in Chain C | Chain A/B/C interface | 0.750023 |
| 549 in Chain C | Chain C internal | 0.852757 |
| 561 in Chain C | Chain A/C interface | 0.877152 |
| 572 in Chain C | Chain C/D interface | 0.78696 |
| 577 in Chain C | Chain B/C interface | 0.741394 |
| 582 in Chain C | Chain B/C interface | 0.866382 |
| 589 in Chain C | Chain B/C interface | 0.84306 |
| 597 in Chain C | Chain C surface | 0.805753 |
| 503 in Chain D | Chain A/D interface | 0.776443 |
| 531 in Chain D | Chain D internal | 0.881503 |
| 546 in Chain D | Chain B/D interface | 0.828317 |

**Table S8. List of waters bound to any chain in the structure that do not have corresponding water molecules in the crystal structures analyzed above found by manually walking through the structure.**

| Water Number and Chain for manually added waters | Location | Qscore |
|---|---|---|
| 905 in Chain A | Chain A surface | 0.61765 |
| 913 in Chain A | Chain A surface | 0.869532 |
| 926 in Chain A | Chain A surface | 0.734997 |
| 951 in Chain A | Chain A/B interface | 0.830304 |
| 954 in Chain A | Chain A surface | 0.834943 |
| 958 in Chain A | Chain A surface | 0.84718 |
| 965 in Chain A | Chain A/B interface | 0.832192 |
| 987 in Chain A | Chain A surface | 0.730228 |
| 995 in Chain A | Chain A surface | 0.801112 |
| 1014 in Chain A | Chain A surface | 0.841313 |
| 1019 in Chain A | Chain A surface | 0.815642 |
| 1033 in Chain A | Chain A surface | 0.868972 |
| 1034 in Chain A | Chain A surface | 0.848711 |
| 1040 in Chain A | Chain A surface near dATP site in Chain A | 0.721635 |
| 1044 in Chain A | Chain A/D interface | 0.815237 |
| 905 in Chain B | Chain B/C interface | 0.647573 |
| 915 in Chain B | Chain B/C interface | 0.754857 |
| 924 in Chain B | Chain B near ATP site | 0.402601 |
| 926 in Chain B | Chain B/C interface | 0.8405 |
| 930 in Chain B | Chain B surface | 0.915631 |
| 932 in Chain B | Chain A/B interface | 0.839068 |
| 938 in Chain B | Chain B surface | 0.654941 |
| 948 in Chain B | Chain B/C interface | 0.868882 |
| 956 in Chain B | Chain B/C interface | 0.781581 |
| 965 in Chain B | Chain B/C interface | 0.875007 |
| 977 in Chain B | Chain B/C interface | 0.789176 |
| 978 in Chain B | Chain B/C interface | 0.815292 |
| 979 in Chain B | Chain B/C interface | 0.800549 |
| 981 in Chain B | Chain B surface | 0.817583 |
| 994 in Chain B | Chain B surface | 0.635165 |
| 997 in Chain B | Chain B surface | 0.794173 |
| 1016 in Chain B | Chain B/C interface | 0.797765 |
| 1018 in Chain B | Chain B/C interface | 0.836799 |
| 1042 in Chain B | Chain B surface | 0.826292 |
| 1052 in Chain B | ATP site | 0.83499 |
| 1055 in Chain B | Chain B/D interface | 0.80752 |

| | | |
|---|---|---|
| 1060 in Chain B | Chain B/C interface | 0.810573 |
| 1069 in Chain B | Chain B surface | 0.727205 |
| 1070 in Chain B | Chain B/C interface | 0.819343 |
| 1071 in Chain B | Chain B surface | 0.786581 |
| 1083 in Chain B | Chain B/C interface | 0.788178 |
| 1084 in Chain B | Chain B/C interface | 0.860176 |
| 1088 in Chain B | Chain B/C interface | 0.855077 |
| 1103 in Chain B | Chain B surface | 0.807777 |
| 1114 in Chain B | Chain B/C interface | 0.903401 |
| 1119 in Chain B | Chain B surface near dATP site in Chain B | 0.820286 |
| 1132 in Chain B | Chain B surface | 0.755591 |
| 1134 in Chain B | Chain B/C interface | 0.808806 |
| 1141 in Chain B | Chain B/C interface | 0.788936 |
| 1147 in Chain B | Chain B/C interface | 0.800612 |
| 1150 in Chain B | Chain B/C interface | 0.75509 |
| 1151 in Chain B | Chain B surface | 0.792592 |
| 1154 in Chain B | Chain B surface near dATP site in Chain B | 0.779656 |
| 1155 in Chain B | Chain B surface | 0.835391 |
| 1162 in Chain B | Chain B surface | 0.754283 |
| 1164 in Chain B | Chain A/B surface | 0.857396 |
| 1168 in Chain B | Chain B surface | 0.811115 |
| 1174 in Chain B | Chain B surface | 0.765418 |
| 1186 in Chain B | Chain B surface near dATP site in Chain B | 0.748357 |
| 1187 in Chain B | Chain B surface near Chain C | 0.742427 |
| 1190 in Chain B | Chain A/B/C interface | 0.808241 |
| 1191 in Chain B | Chain B surface | 0.788587 |
| 502 in Chain C | Chain B/C interface | 0.766246 |
| 511 in Chain C | $Fe_2O$ site | 0.544792 |
| 513 in Chain C | Chain B/C interface | 0.813343 |
| 519 in Chain C | Chain B/C interface | 0.823887 |
| 520 in Chain C | Chain C/D interface | 0.626192 |
| 521 in Chain C | Chain B/C interface | 0.760901 |
| 535 in Chain C | Chain B/C interface | 0.845528 |
| 539 in Chain C | Chain A/C interface | 0.732206 |
| 540 in Chain C | Chain C surface | 0.794983 |
| 542 in Chain C | Chain B/C interface | 0.832889 |
| 543 in Chain C | Chain B/C interface | 0.763121 |
| 550 in Chain C | Chain C surface | 0.798474 |

| | | |
|---|---|---|
| 556 in Chain C | Chain B/C interface | 0.897779 |
| 562 in Chain C | Chain B/C interface | 0.746886 |
| 563 in Chain C | Chain B/C/D interface | 0.736722 |
| 565 in Chain C | Chain B/C interface | 0.886891 |
| 567 in Chain C | Chain B/C interface | 0.859275 |
| 570 in Chain C | Chain C/D interface | 0.882411 |
| 578 in Chain C | Chain C surface | 0.875736 |
| 584 in Chain C | Chain B/C interface | 0.87105 |
| 586 in Chain C | Chain B/C interface | 0.76788 |
| 592 in Chain C | Chain A/C interface | 0.744232 |
| 596 in Chain C | Chain A/C interface | 0.881656 |
| 599 in Chain C | Chain B/C interface | 0.804872 |
| 600 in Chain C | Chain A/B/C interface | 0.808845 |
| 602 in Chain C | Chain A/B/C interface | 0.800009 |
| 605 in Chain C | Chain B/C interface | 0.891546 |
| 606 in Chain C | Chain C surface | 0.812837 |
| 607 in Chain C | Chain A/B/C interface | 0.79152 |
| 504 in Chain D | $Fe_2O$ site | 0.669508 |
| 505 in Chain D | Chain C/D interface | 0.684958 |
| 509 in Chain D | Chain A/D interface | 0.803428 |
| 529 in Chain D | Chain D surface | 0.555955 |
| 532 in Chain D | Chain D surface near Chain B | 0.841204 |
| 538 in Chain D | Chain A/D interface | 0.831821 |
| 543 in Chain D | Chain B/C/D interface | 0.793863 |
| 548 in Chain D | Chain D surface near Chain B | 0.892428 |
| 562 in Chain D | Chain B/D interface near Chain C | 0.739774 |
| 566 in Chain D | Chain D surface | 0.698096 |
| 568 in Chain D | Chain B/D interface near Chain C | 0.773393 |

**Table S9. Direct interactions across β′–α, β–α′, β′–α′ subunit interfaces.** Distances less than or equal to 3.5 Å between key residues in the indicated subunits.

| *β'-α Interactions* | |
|---|---|
| *Residues* | *Distance (Å)* |
| K229β' (NZ) – E397α (OE1) | 3.4 |
| K229β' (NZ) – E397α (OE2) | 2.5 |
| *β-α' Interactions* | |
| N131β – E314α′ (backbone O) | 3.2 |
| R127β – N322α′ (backbone O) | 2.8 |
| R127β – R323α′ (backbone O) | 3.3 |
| R127β (backbone O) – N328α′ | 3.3 |
| S134β – R323α′ | 2.9 |
| *β'-α' Interactions* | |
| D362β′ (backbone N) – V709α′ (backbone O) | 3.2 |
| D366β′ (backbone N) – Q712α′ | 3.1 |
| D29β′ – R323α′ (NE) | 3.3 |
| D29β′ – R323α′ (NH2) | 3.4 |
| Q30β′ – R323α′ | 2.7 |
| Q360β′ (backbone O) – V709α′ (backbone N) | 3.3 |
| D362β′ (backbone N) – V709α′ (backbone O) | 3.2 |
| D362β′ (backbone O) – M711α′ (backbone N) | 2.5 |
| D368β′ (OD1) – Q712α′ | 2.5 |
| D368β′ (OD2) – Q712α′ | 3.4 |

**Table S10. Direct interactions across the α–β interface.** Distances less than or equal to 3.5 Å between key residues in the β and α subunits. β-C-terminus interactions not observed in the previously published α2β2 structure are bolded. Note that not all observed interactions were reported in **Table S2** of the previous publication.

| *β-C-Terminus* | |
|---|---|
| *Residues* | *Distance (Å)* |
| **D342β (backbone O) – W670α** | 3.3 |
| D342β (OD1) – R735α (NE) | 2.6 |
| D342β (OD2) – R735α (NH2) | 3.0 |
| D342β (OD1) – R735α (NH2) | 3.4 |
| N343β (backbone O)– I644α (backbone N) | 3.0 |
| N343β – V642α (backbone O) | 2.8 |
| Q345β (backbone N) – I644α (backbone O) | 2.6 |
| Q345β (backbone O) – A646α (backbone N) | 3.5 |
| **Q345β – E623α** | 2.5 |
| A347β (backbone N) – A646α (backbone O) | 3.0 |
| **P348β (backbone O) – R411α** (NH1) | 2.5 |
| **P348β (backbone O) – R411α** (NH2) | 3.1 |
| **Q349β – R411α** | 3.4 |
| **Q349β – Y731α** | 3.1 |
| Q349β – Y730α | 3.4 |
| Q349β – D334α | 3.1 |
| E350β – K154α | 3.1 |
| **E350β – T624α** | 3.4 |
| **S355β – T734α (backbone O)** | 2.8 |
| Q360β – G707α (backbone O) | 3.4 |
| **Q360β (backbone O) – V709α (backbone N)** | 3.1 |
| D362β (backbone N) – V709α (backbone O) | 2.8 |
| **D362β – K708α** | 3.5 |
| D366β (backbone N) – Q712α | 2.8 |
| D366β (backbone O) – Q712α | 3.3 |
| **L375β (backbone O) – K584α** | 2.4 |
| *Observed Previously (PDB 6W4X) but Not Present in Current Structure* | |
| *Residues* | *Distance in Current Structure (Å) (Distance in Previous Structure (Å))* |
| Q349β – Y413α | 3.7 (3.2) |
| E350β – S647α | 4.2 (2.4) |
| *Other Region* | |
| *Residues* | *Distance (Å)* |
| K42β – D736α (backbone O) | 3.3 |

| | |
|---|---|
| R49β – N322α (backbone O) | 3.2 |
| R49β – G324α (backbone O) | 2.7 |
| E52β – N322α | 2.8 |
| E220β – K64α | 3.1 |
| R236β – D736α | 3.5 |
| N300β – I47α (backbone O) | 3.0 |
| Y307β – Q742α | 2.7 |
| Y310β – D743α (backbone O) | 2.2 |
| Y310β – D743α (backbone N) | 3.0 |
| R315β (NH1) – D740α (OD1) | 3.3 |
| R315β (NH2) – D740α (OD2) | 2.9 |
| N330β – Q742α (backbone O) | 3.0 |
| N336β – Q742α (backbone O) | 2.6 |
| N336β – Q742α (backbone N) | 2.8 |
| L339β (backbone O) – D740α (backbone N) | 3.3 |
| S341β (backbone N) – A738α (backbone O) | 3.3 |
| S341β – D736α | 2.7 |

**Table S11. Water-bridged interactions across the α–β interface.** Waters newly identified in this structure are bolded.

| *β-C-Terminus* | |
|---|---|
| *Residues* | *Bridging Water Number and Chain* |
| E350β, SN-CDPα | 1080 in Chain B |
| E350β, G299α (backbone O), SN-CDPα | **982 in Chain B** |
| S350β (backbone O), D736α | **955 in Chain B** |
| G359β (backbone N), Q404α | 573 in Chain C |
| G349β, R411α, D334α | 503 in Chain C |
| V346β (backbone O), R639α | **562 in Chain C** |
| E352β, K320α | 1120 in Chain B |
| Q345β, R639α (backbone O) | 1019 in Chain B |
| S355β, T734α (backbone O) | 558 in Chain C |
| E350β, T624α | 1056 in Chain B |
| D342β (backbone O), V344β (backbone O), W670α, R639α | 537 in Chain C |
| S363β, S400α | 1110 in Chain B |
| E350β, D649α | **502 in Chain C** |
| E364β (backbone O, N), M711α (backbone N) | 506 in Chain C |
| Q345β, E623α | **1141 in Chain B** |
| D342β, D736α, D737α (backbone N) | **1134 in Chain B** |
| E352β (backbone N), E225β, K648α | **521 in Chain C** |
| *Other Region* | |
| *Residues* | *Bridging Water Number* |
| D226β (backbone N), E225β (backbone N), D58β, K648α | 1057 in Chain B |
| M296β (backbone O), S295β, Q48α (backbone O) | 545 in Chain C |
| E220β (backbone O), I651α (backbone O) | 576 in Chain C |
| R57β, V297α (backbone O) | **519 in Chain C** |
| P333β (backbone O), R653α | **1147 in Chain B** |
| R315β, A741α (backbone O) | **905 in Chain B** |

**Table S12. Water-bridged interactions across β′–α, β–α′, β′–α′ subunit interfaces.** Waters newly identified in this structure are bolded.

| *β'-α Interactions* | |
|---|---|
| *Residues* | *Bridging Water Number and Chain* |
| E52β′, Q404α | 1032 in Chain B |
| F47β′ (backbone O), R323α | **546 in Chain D** |
| E52β′ (backbone O), Q404α | **543 in Chain D** |
| *β-α' Interactions* | |
| R59(C)β, E312α′, T276α′ (backbone O), C278α′ (backbone N), I279α′ (backbone N) | 962 in Chain A |
| R59(B)β, N328α′ (backbone O), S315α′ (backbone O), S315α′ | 967 in Chain A |
| R59(C)β, S315α′, E312α′ | 972 in Chain A |
| R59(B)β, N131β, R329α′ | 526 in Chain C |
| *β'-α' Interactions* | |
| D362β′, D362β′ (backbone O), E364β′ (backbone O), E364β′ (backbone N), M711α′ (backbone N), Q712α′ (backbone N) | **503 in Chain D** |
| Q360β′, A407α′ (backbone O) | 550 in Chain D |
| D29β′ (backbone O), R323α′ | **1044 in Chain A** |

**Table S13. List of PCET pathway interactions with water molecules in α and β.** Interaction distances are listed and waters newly identified in this structure are bolded.

| Interacting Residue or Water | Water Number and Chain | Distance (Å) |
|---|---|---|
| $Fe_2O$ (FE1) | **511 in Chain C** | 2.0 |
| Y122β (OH) | **511 in Chain C** | 3.2 |
| D237β (OD1) | 517 in Chain C | 2.4 |
| R236β (NE) | 517 in Chain C | 2.3 |
| R236β (NH2) | 517 in Chain C | 2.8 |
| R236β (NH2) | 536 in Chain C | 3.5 |
| Y356β (backbone O) | 536 in Chain C | 2.7 |
| Water 26 | 536 in Chain C | 3.2 |
| E52β (OE2) | 546 in Chain C | 2.7 |
| **Water 1150 in Chain B** | 546 in Chain C | 3.4 |
| V353β (backbone O) | 516 in Chain C | 2.3 |
| E52β (OE2) | 516 in Chain C | 2.9 |
| Y356β (OH) | 507 in Chain C | 2.2 |
| N322α (ND2) | **1150 in Chain B** | 3.3 |
| E52β (backbone O) | **582 in Chain C** | 3.2 |
| **Water 936 in Chain B** | **582 in Chain C** | 2.3 |
| **Water 1088 in Chain B** | **936 in Chain B** | 3.5 |
| Water 594 in Chain C | **1088 in Chain B** | 2.9 |
| V353β (backbone N) | 594 in Chain C | 3.4 |
| Y731α (OH) | 1014 in Chain B | 2.7 |
| E623α (OE2) | 1014 in Chain B | 3.8 |
| Water 1027 in Chain B | 1014 in Chain B | 3.0 |
| C439α (SG) | 1027 in Chain B | 3.0 |
| SN-CDP phosphate group (O1A) | 1027 in Chain B | 2.7 |
| SN-CDP base (N4) | **982 in Chain B** | 2.5 |
| Q349β (backbone O) | **982 in Chain B** | 2.7 |
| SN-CDP base (N3) | **982 in Chain B** | 3.5 |
| Water 1080 in Chain B | 564 in Chain C | 2.6 |
| Q349β (backbone N) | 564 in Chain C | 2.9 |
| Water 1014 in Chain B | 564 in Chain C | 2.9 |
| SN-CDP base (N4) | 1080 in Chain B | 2.9 |
| Water 918 in Chain B | 1080 in Chain B | 3.1 |
| SN-CDP phosphate group (O1B) | 918 in Chain B | 2.3 |
| SN-CDP phosphate group (O2B) | 918 in Chain B | 2.2 |
| SN-CDP phosphate group (O3A) | **984 in Chain B** | 3.1 |

| | | |
|---|---|---|
| SN-CDP ribose oxygen (O4') | **984 in Chain B** | 2.8 |
| SN-CDP phosphate group (O3A) | 992 in Chain B | 2.6 |
| SN-CDP phosphate group (O2A) | 992 in Chain B | 3.2 |

**Table S14. List of PDB IDs for crystal structures used to identify previously observed water structures within the current cryo-EM density.**

| PDB ID | Resolution (Å) | Subunit |
|---|---|---|
| 1R1R | 2.90 | $\alpha$2 |
| 2X0X | 2.30 | $\alpha$2 |
| 2XAP | 2.10 | $\alpha$2 |
| 2XAX | 2.75 | $\alpha$2 |
| 2XAY | 2.65 | $\alpha$2 |
| 2XAZ | 2.60 | $\alpha$2 |
| 2XO4 | 2.50 | $\alpha$2 |
| 2XO5 | 2.70 | $\alpha$2 |
| 5CNS | 2.98 | $\alpha$2 |
| 8VHN | 2.62 | $\alpha$2 |
| 2XOF | 2.20 | $\beta$2 |
| 5CI0 | 2.25 | $\beta$2 |
| 5CI1 | 1.95 | $\beta$2 |
| 5CI2 | 2.25 | $\beta$2 |
| 5CI3 | 2.40 | $\beta$2 |
| 5CI4 | 2.05 | $\beta$2 |

**Table S15. Additional hydrogen bonding interactions by PCET waters not depicted in Figure S14.** Interaction distances are listed and waters newly identified in this structure are bolded.

| Water Number and Chain | Interacting Residue or Water | Distance (Å) |
|---|---|---|
| **Water 511 in Chain C** | E238β | 2.5 |
| **Water 511 in Chain C** | D84β | 2.6 |
| Water 517 in Chain C | L233β (backbone O) | 3.1 |
| Water 517 in Chain C | F46β (backbone O) | 3.5 |
| Water 536 in Chain C | F47β (backbone O) | 2.6 |
| Water 546 in Chain C | N/A | N/A |
| **Water 1150 in Chain B** | S408α (backbone O) | 3.4 |
| **Water 1150 in Chain B** | Water 1140 in Chain B | 2.3 |
| **Water 936 in Chain B** | K320α (backbone O) | 2.3 |
| **Water 1088 in Chain B** | T409α (backbone O) | 3.0 |
| Water 594 in Chain C | Water 1120 in Chain B | 3.3 |
| Water 507 in Chain C | N/A | N/A |
| Water 1104 in Chain B | Water 1115 in Chain B | 2.2 |
| Water 1104 in Chain B | T734α (backbone N) | 3.0 |
| Water 1104 in Chain B | S355β (backbone O) | 3.3 |
| Water 1014 in Chain B | N/A | N/A |
| Water 1080 in Chain B | Q350β | 3.1 |
| Water 918 in Chain B | T624α (backbone N) | 3.3 |
| Water 918 in Chain B | R298α | 3.5 |
| Water 918 in Chain B | T624α | 2.7 |
| Water 918 in Chain B | SN-CDP phosphate group (O1A) | 3.0 |
| Water 1027 in Chain B | N/A | N/A |
| **Water 982 in Chain B** | G299α (backbone O) | 2.7 |
| **Water 984 in Chain B** | S224α | 2.3 |
| Water 992 in Chain B | S625α | 3.5 |
| Water 992 in Chain B | Water 1179 in Chain B | 2.7 |

**Table S16. List of waters found within the two water channels.** Interaction distances are listed and waters newly identified in this structure are bolded.

| **Yellow Channel** | | |
|---|---|---|
| **Waters located within channel** | **Interacting Residue or Water** | **Distance (Å)** |
| Water 546 in Chain C | Water 536 in Chain C | 3.2 |
| Water 546 in Chain C | **Water 1150 in Chain B** | 3.4 |
| Water 546 in Chain C | E52β | 2.7 |
| Water 536 in Chain C | F47β (backbone O) | 2.6 |
| Water 536 in Chain C | Y356β (backbone O) | 2.7 |
| Water 536 in Chain C | R236β | 3.5 |
| Water 1140 in Chain B | **Water 1150 in Chain B** | 2.3 |
| Water 1140 in Chain B | S408α (backbone O) | 3.2 |
| Water 1140 in Chain B | S408α | 3.5 |
| Water 1140 in Chain B | R323α | 3.3 |
| Water 1140 in Chain B | N322α (OD1) | 3.4 |
| Water 1140 in Chain B | N322α (ND2) | 3.5 |
| Water 1140 in Chain B | N322α (backbone N) | 3.5 |
| Water 1109 in Chain B | Water 1006 in Chain A | 3.5 |
| Water 1109 in Chain B | Water 580 in Chain C | 2.1 |
| Water 1109 in Chain B | K290α (backbone O) | 3.1 |
| Water 1006 in Chain A | **Water 1178 in Chain B** | 2.6 |
| Water 1006 in Chain A | Water 1109 in Chain B | 3.5 |
| Water 1006 in Chain A | K290α (backbone O) | 3.2 |
| Water 1006 in Chain A | S291α (backbone O) | 3.2 |
| Water 1006 in Chain A | T276α' | 3.0 |
| **Water 582 in Chain C** | **Water 936 in Chain B** | 2.3 |
| **Water 582 in Chain C** | E52β (backbone O) | 3.2 |
| **Water 1178 in Chain B** | **Water 1038 in Chain B** | 2.5 |
| **Water 1178 in Chain B** | Water 603 in Chain C | 3.1 |
| **Water 1178 in Chain B** | Water 1182 in Chain B | 2.4 |
| Water 591 in Chain C | Q349β (backbone O) | 3.3 |
| **Water 1038 in Chain B** | **Water 1178 in Chain B** | 2.5 |
| **Water 1038 in Chain B** | C292α (backbone O) | 2.8 |
| **Water 1038 in Chain B** | G300α (backbone N) | 3.0 |
| Water 594 in Chain C | Water 1120 in Chain B | 3.3 |
| Water 594 in Chain C | **Water 1088 in Chain B** | 2.9 |
| Water 594 in Chain C | V353β (backbone N) | 3.4 |
| **Water 936 in Chain B** | **Water 1088 in Chain B** | 3.5 |
| **Water 936 in Chain B** | K320α | 2.3 |

| | | |
|---|---|---|
| Water 603 in Chain C | Water 1182 in Chain B | 3.4 |
| Water 603 in Chain C | **Water 602 in Chain C** | 3.3 |
| Water 1120 in Chain B | Water 958 in Chain B | 3.3 |
| Water 1120 in Chain B | E352β | 3.4 |
| Water 1120 in Chain B | K320α | 3.1 |
| Water 1040 in Chain B | K290α | 2.8 |
| Water 1182 in Chain B | **Water 584 in Chain C** | 3.2 |
| Water 1182 in Chain B | **Water 602 in Chain C** | 3.0 |
| Water 1182 in Chain B | **Water 600 in Chain C** | 2.3 |
| **Water 584 in Chain C** | R57β | 3.2 |
| Water 958 in Chain B | Water 503 in Chain C | 3.3 |
| Water 958 in Chain B | R411α (NE) | 3.3 |
| Water 958 in Chain B | R411α (NH2) | 2.4 |
| Water 958 in Chain B | D334α | 2.9 |
| Water 580 in Chain C | S56β | 3.2 |
| **Water 1150 in Chain B** | S408α (backbone O) | 3.4 |
| **Water 1150 in Chain B** | N322α | 3.3 |
| **Water 1088 in Chain B** | T409α (backbone O) | 3.0 |
| **Water 567 in Chain C** | **Water 543 in Chain C** | 2.4 |
| **Water 567 in Chain C** | P348β (backbone O) | 3.0 |
| **Water 602 in Chain C** | See previous entries | See previous entries |
| **Water 600 in Chain C** | **Water 519 in Chain C** | 3.3 |
| **Water 513 in Chain C** | R57β (backbone N) | 3.4 |
| **Water 513 in Chain C** | S56β | 3.2 |
| **Water 513 in Chain C** | D54β | 2.3 |
| **Water 513 in Chain C** | D54β (backbone O) | 3.4 |
| **Water 543 in Chain C** | Q349β (backbone O) | 2.7 |
| Water 516 in Chain C | V353β (backbone O) | 2.3 |
| Water 516 in Chain C | E52β | 2.9 |
| **Pink Channel** | | |
| **Waters located within channel** | **Interacting Residue or Water** | **Distance (Å)** |
| Water 507 in Chain C | Y356β | 2.2 |
| **Water 1093 in Chain B** | Water 1115 in Chain B | 3.1 |
| **Water 1093 in Chain B** | **Water 577 in Chain C** | 2.9 |
| **Water 1093 in Chain B** | **Water 562 in Chain C** | 2.3 |
| **Water 1093 in Chain B** | R639α (NH1) | 3.1 |
| **Water 1093 in Chain B** | R639α (NH2) | 3.0 |
| **Water 577 in Chain C** | **Water 586 in Chain C** | 2.3 |
| **Water 577 in Chain C** | V344β (backbone O) | 3.1 |

| | | |
|---|---|---|
| **Water 562 in Chain C** | Water 1129 in Chain B | 2.6 |
| **Water 562 in Chain C** | R639α | 3.4 |
| **Water 562 in Chain C** | V346β (backbone O) | 2.9 |
| Water 558 in Chain C | Water 537 in Chain C | 3.5 |
| Water 558 in Chain C | **Water 1134 in Chain B** | 2.4 |
| Water 558 in Chain C | T734α (backbone O) | 3.4 |
| Water 558 in Chain C | S355β | 3.0 |
| Water 558 in Chain C | S341β (backbone O) | 2.9 |
| Water 537 in Chain C | W670α | 3.2 |
| Water 537 in Chain C | R639α | 2.9 |
| Water 537 in Chain C | V344β (backbone O) | 3.4 |
| Water 537 in Chain C | D342β (backbone O) | 2.6 |
| **Water 1134 in Chain B** | G737α (backbone N) | 3.4 |
| **Water 1134 in Chain B** | D736α | 3.2 |
| **Water 1134 in Chain B** | D342β | 3.4 |
| **Water 1134 in Chain B** | S341β (backbone N) | 3.4 |
| **Water 1134 in Chain B** | S341β | 3.3 |
| **Water 1114 in Chain B** | A738α (backbone O) | 3.1 |
| **Water 1114 in Chain B** | D342β (backbone N) | 3.3 |
| **Water 586 in Chain C** | S354β (backbone N) | 3.2 |
